# A Microfluidic Platform for Spatiotemporal Dissection of Neurodegeneration Across Hierarchical Human Neural Circuits

**DOI:** 10.64898/2026.07.10.737646

**Authors:** Joël Küchler, Areli Balderas Maza, Giulia Amos, Luc Jordi, Elena Durante, Yara Roth, Vaiva Vasiliauskaitė, Benedikt Maurer, Magdalini Polymenidou, Kan Cao, Nicolai Winter-Hjelm

## Abstract

Understanding how neurodegenerative diseases initiate and propagate through neural circuits remains a fundamental challenge in neuroscience. The earliest stages occur years before symptoms emerge, making them inaccessible to study in patients. Microfluidic platforms, where neurons communicate across chambers through microchannels accessible only to their axons, have opened new experimental avenues. However, existing models lack the complexity and precision needed to track how individual circuit components respond to focal pathological changes over time. Here we present a 33-chamber cortical network-on-chip integrating human iPSC-derived excitatory neurons, inhibitory neurons, and astrocytes in a six-layer feedforward architecture recapitulating the laminar structure of the neocortex. Amyloid-β is applied globally, while progerin-induced accelerated ageing in a single chamber establishes a defined disease core. Continuous recordings using high-density microelectrode arrays reveal progressive, layer-dependent changes in firing dynamics and network topology. Machine-learning-based feature analysis identifies a multiparametric electrophysiological signature distinguishing healthy from disease-affected chambers, enabling studies of the earliest timepoint at which pathology becomes detectable. This establishes a scalable framework for mechanistic studies of neurodegeneration and identification of electrophysiological biomarkers of disease progression.

## Introduction

Neurodegenerative diseases are among the most devastating challenges in medicine, yet directly studying their onset in the human brain at the time and cellular resolution at which it occurs remains impossible. In Alzheimer’s disease (AD), pathological changes are thought to emerge in cortical and hippocampal circuits years before clinical symptoms appear (1, 2). By the time of diagnosis, widespread neurodegeneration has already occurred, rendering the earliest and most therapeutically relevant stages of the disease inaccessible to direct study in patients. During this preclinical period, neural circuits adapt and reorganise in response to ongoing pathology, and these changes may represent both early indicators of disease and targets for therapeutic intervention (3–5). Understanding how pathology spreads through a circuit, and how the surrounding network responds, is therefore essential for detecting the disease at its earliest stages, identifying electrophysiological biomarkers of disease progression, and developing targeted interventions. Experimental platforms that can recapitulate these dynamics in a controlled human context are critically needed.

The use of microfluidic platforms to structure neural networks *in vitro* has emerged as a powerful approach for studying neural network topology, function, and dysfunction under controlled conditions at cellular resolution. In these platforms, distinct neural populations are segregated in interconnected chambers linked by microchannels that are only accessible to their neurites, allowing neurons to communicate through defined axonal connections while their cell bodies remain spatially segregated (6, 7). This promotes the emergence of modular network dynamics that balance segregated and integrated activity akin to that observed across brain regions *in vivo* (8–10). Anatomically relevant circuit hierarchies between distinct neural populations can be recapitulated by controlling the cell type composition of individual chambers (11–13). Geometrical constraints can furthermore be implemented in the microchannels to promote unidirectional axonal outgrowth between populations (14–16). Tesla valve-inspired designs, for example, guide axons growing in the non-preferred direction back toward their chamber of origin, enforcing feedforward hierarchical connectivity between defined neural populations (8, 17). Interfacing these platforms with microelectrode arrays (MEAs) enables the study of functional network dynamics across the full circuit, and recent advances have shown that structured networks can be established directly on high-density complementary metal-oxide-semiconductor (CMOS) MEAs, enabling stimulation and recording with single-unit spatiotemporal resolution (18).

A critical advantage of segregating populations into defined chambers is that pathological perturbations can be introduced into a single chamber while the surrounding network remains unperturbed. This enables direct study of how pathology spreads and how connected populations respond, a question central to many neurodegenerative diseases (12, 19–22). Recent work has demonstrated that multi-layer, multi-node microfluidic circuits show spatially structured functional reorganisation in response to localised perturbations, with adaptive changes detectable across directly connected and distal populations (17). However, existing platforms have been limited to a small number of chambers containing thousands of cells each, providing only a sparse and population-averaged view of network activity. Most studies have also relied on rodent-derived neurons, limiting translational relevance to human disease (17). Furthermore, existing studies have typically relied on discrete recording sessions, limiting the temporal resolution at which network dynamics can be tracked longitudinally.

Here we present a 33-chamber microfluidic platform integrating human iPSC-derived neurons and astrocytes in a six-layer feedforward architecture with defined directional connectivity. The platform is interfaced with High-Density MEAs (HD-MEAs) providing continuous electrophysiological recordings from approximately 1000 electrodes across the full network for weeks. Progerin-induced accelerated ageing in a single chamber, combined with a global amyloid-β perturbation, establishes a defined disease core within the network. Continuous recordings reveal progressive, layer-dependent disruption of network dynamics emanating from the disease-affected chamber, captured through transfer entropy-based functional connectivity analysis, graph-theoretic mapping, and machine learning classification of individual chamber states. This platform establishes a scalable framework for mechanistic studies of neurodegeneration and for the identification of electrophysiological biomarkers of disease progression.

## Results

### The six-layer, 33-node microfluidic platform establishes defined directional connectivity and physiological cell-type composition across a human neural circuit

A six-layer, 33-node microfluidic platform was developed to construct layered neural circuits with directional connectivity across the layers **(Fig. 1A)**. Spheroids expressing either GFP or RFP were seeded into the chambers (hereafter referred to as nodes) in alternating layers. Nodes within each layer were interconnected by bidirectional straight channels permitting symmetric axonal exchange, while interlayer feedforward connections were mediated by asymmetric Tesla valve microchannels. In these unidirectional channels, funnel-shaped openings at the presynaptic side guide axons into the channel entrance, while sawtooth structures at the postsynaptic side misdirect neurites from the postsynaptic spheroid away from the channel entrances. Neurites entering from the non-preferred direction are redirected by the Tesla valve geometry back toward their chamber of origin, ensuring that only presynaptic axons successfully traverse the channel to innervate the postsynaptic spheroid **(Fig. 1B)** (8). Fluorescence imaging at 14 days *In Vitro* (DIV) confirmed successful network formation across all 33 nodes **(Fig. 1A)**, and magnified views at the single-channel level confirmed effective redirection of neurites from the postsynaptic side by the Tesla valves **(Fig. 1C)**. Quantification of axon density along the channel axis at DIV 14 revealed significantly higher fluorescence intensity in the preferred versus non-preferred direction in unidirectional channels, while bidirectional channels showed no significant directional bias **(Fig. 1D–E; Fig. S1;** * * * * *p <* 0.0001; ns, not significant). Axon bundles travelling in the undesired direction were significantly narrower than those in the desired direction in unidirectional channels, while no significant difference in neurite width was observed in bidirectional channels **(Fig. 1F;** * * * * *p <* 0.0001; ns, not significant). This directional bias was maintained through DIV 35, confirming that asymmetric connectivity was established early and sustained throughout network maturation **(Fig. S2)**.

**Figure 1.**
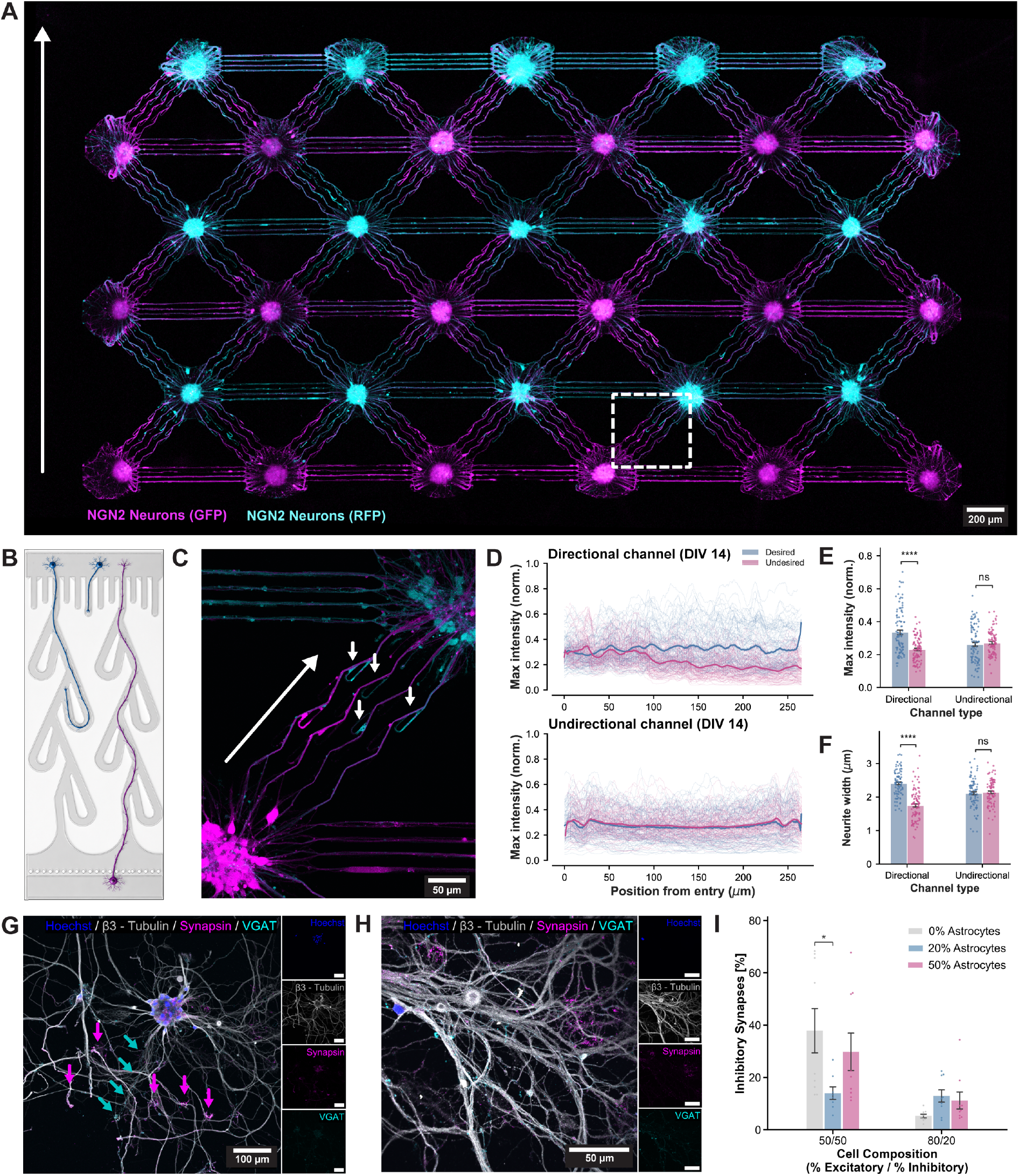
A 33-node feedforward human neural circuit establishes directional inter-layer connectivity and physiological excitatory/inhibitory composition. **(A)** Fluorescence overview image of the 33-node platform at DIV 14, showing NGN2-induced neurons expressing GFP (magenta) and RFP (cyan) seeded in alternating layers. Within each layer, nodes were connected by bidirectional (straight) channels, while feedforward connections across layers were mediated by unidirectional channels. The white arrow indicates the feedforward direction. **(B)** Schematic top view of a unidirectional microfluidic channel illustrating the asymmetric Tesla valve geometry designed to bias axonal growth in a preferred direction. Tesla valves redirected neurites from the non-preferred (blue) cell population back toward their chamber of origin. **(C)** Magnified view of the dashed region in **(A)**, demonstrating unidirectional connectivity from the magenta to the cyan spheroid at DIV 14. Small arrows highlight neurites from the cyan population being redirected by the Tesla valves back toward their chamber of origin. **(D)** Full intensity profiles along the channel axis (0–270 µm from channel entry) for desired (blue) and undesired (pink) directions in unidirectional (top) and bidirectional (bottom) channels at DIV 14, showing individual profiles (thin lines) and mean (thick lines). **(E)** Unidirectional channels exhibited significantly higher axon growth in the preferred versus non-preferred direction at DIV 14 (max fluorescence intensity, normalised), while bidirectional channels showed no significant directional bias. \*\*\*\**p <* 0.0001; ns, not significant. **(F)** Axon bundles travelling in the undesired direction were significantly narrower than those travelling in the desired direction in unidirectional channels, consistent with selective passage of axons from a single population through the Tesla valve geometry. No significant difference in neurite width was observed in bidirectional channels. \*\*\*\**p <* 0.0001; ns, not significant. **(G–H)** Immunofluorescence of neurons co-stained for Hoechst (blue), *β*3-Tubulin (gray), Synapsin (magenta), and VGAT (cyan) confirmed the formation of both excitatory and inhibitory synapses within the network. VGAT puncta co-localising with Synapsin identified the inhibitory fraction of the total synapse population. **(I)** Quantification of inhibitory synapse proportions across cell compositions and astrocyte co-culture conditions. At a 50/50 excitatory/inhibitory composition with 20% astrocytes, inhibitory synapses constituted a mean of 14.0 ± 2.4% (SE) of total synapses, consistent with the reported range of inhibitory synapses in the human cortex (∼15–20%).

To establish a physiologically relevant excitatory/inhibitory balance, the cell type ratio and astrocyte fraction were systematically varied across spheroids of 50 cells. Astrocytes were included in all conditions given their well-established roles in synapse formation, maturation, and network stability (23, 24). Excessively high astrocyte fractions were avoided, as an overabundance of astrocytes risks occluding microchannel entrances and shielding electrodes from neurons during recordings (25, 26). Immunostaining for *β*3-Tubulin, Synapsin, and the inhibitory synapse marker VGAT confirmed the presence of both excitatory and inhibitory synapses across all conditions **(Fig. 1G–H)**. An astrocyte fraction of 20% yielded consistent inhibitory synapse percentages irrespective of E/I ratio **(Fig. 1I)**, broadly matching estimates in the human cortex (∼ 15–20%) (27). A 50/50 E/I seeding ratio was therefore selected to compensate for the lower viability of GABAergic neurons following dissociation and seeding, ensuring sufficient inhibitory neuron representation across all spheroids. At this composition, inhibitory synapses constituted 14.0 ± 2.4% (SE) of total synapses. This condition was adopted as the standard spheroid composition for all subsequent experiments.

### Progerin expression in a single network node combined with a global amyloid-*β* perturbation establishes a human neural circuit model for AD-relevant network perturbation

Having established a 33-node feedforward human neural circuit with defined directional connectivity and physiological cell composition, AD-relevant disease perturbations were introduced to enable spatially resolved interrogation of network-level responses. Ageing represents the single greatest risk factor for AD, and ageing-associated molecular changes occur in a cell-type-specific manner across the human brain (28, 29). To introduce a site of accelerated ageing-associated vulnerability within the platform, progerin expression was induced in a single node of the 33-node network via lentiviral transduction **(Fig. 2A– B)**. Progerin is a truncated form of Lamin A that drives accelerated cellular ageing and has been used to establish *in vitro* models recapitulating early AD-relevant pathological features, including amyloid accumulation and tau hyper-phosphorylation (30, 31). While progerin expression does not fully recapitulate the complexity of physiological neural ageing, it provides a tractable means of introducing defined ageing-associated stress into individual nodes of a structured network. CellROX staining indicated a modest increase in oxidative stress in progerin-expressing spheroids **(Fig. 2C)**, consistent with the increased reactive oxygen species reported in progerin-expressing cells (30, 32).

**Figure 2.**
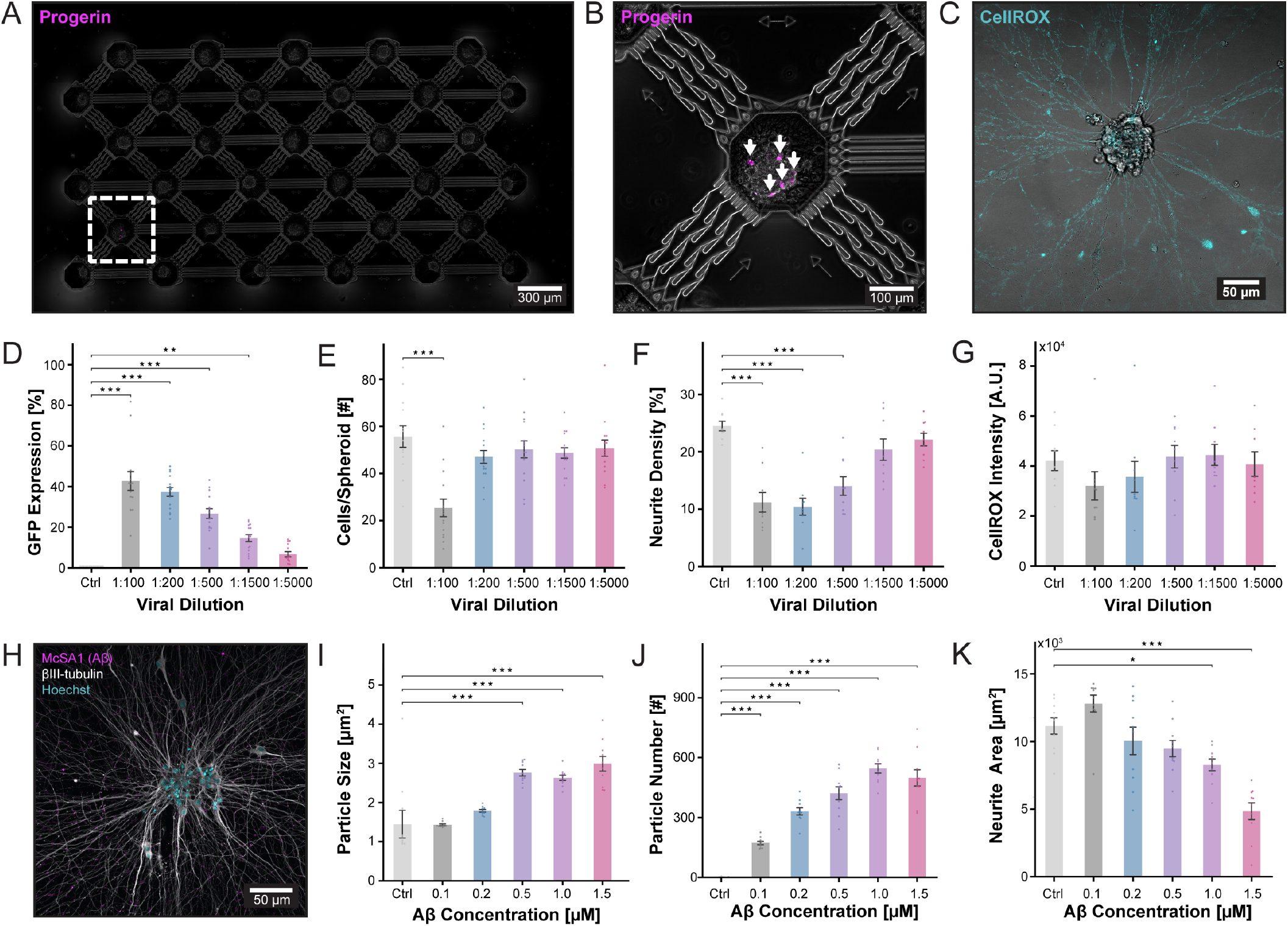
Spatially targeted progerin expression and global amyloid-*β* perturbation establish a human neural circuit model for AD-relevant network perturbation. **(A)** Confocal overview image of the 33-node platform following lentiviral induction of progerin expression in a single spheroid (dashed box). **(B)** Magnified view of the progerin-expressing node, with arrows indicating GFP-positive (magenta) cells, confirming successful lentiviral transduction within a single node of the network. **(C)** Representative CellROX fluorescence image of a progerin-expressing spheroid, showing a trend toward elevated oxidative stress consistent with progerin-induced cellular stress (30, 32). **(D–G)** Lentiviral dilution titration across five concentrations (1:100–1:5000) revealed dose-dependent changes in GFP-progerin expression **(D)**, cell number per spheroid, indicating viability **(E)**, and neurite density **(F)**, alongside a non-significant trend toward increasing oxidative stress at higher viral loads **(G)**, informing selection of an optimal viral dose. Statistical comparisons are shown against the control condition only; full pairwise statistics are reported in Supplementary Tables 3–10. \*\*\**p <* 0.001; \*\**p <* 0.01; \**p <* 0.05; ns, not significant. **(H)** Representative immunofluorescence image of McSA1 (magenta) and *β*III-tubulin (gray) at DIV 37, 14 days after global amyloid-*β* addition at a working concentration of 0.2 µM at DIV 23, confirming the formation of perineuronal A*β* aggregates (arrows) within the neurite network. **(I–K)** Concentration-dependent effects of A*β* treatment (0.1–1.5 µM) on average aggregate size **(I)**, perineuronal aggregate count **(J)**, and neurite mesh area **(K)**, demonstrating significant dose-dependent aggregate accumulation and progressive neurite loss, informing selection of a working concentration for subsequent network experiments. Only statistical comparisons to the control condition are shown. Full pairwise statistics are reported in Supplementary Tables 11–13. \*\*\**p <* 0.001; \*\**p <* 0.01; \**p <* 0.05; ns, not significant.

To determine an optimal viral dose that induced progerin expression without causing immediate network degradation, a lentiviral dilution titration was performed across a range of concentrations **(Fig. 2D–G)**. GFP-progerin expression decreased monotonically with increasing dilution **(Fig. 2D); Fig. S3)**, while cell number per spheroid was significantly reduced only at the highest viral concentrations **(Fig. 2E)**. Neurite density declined significantly at a 1:500 dilution and below **(Fig. 2F); Fig. S4)**, and a non-significant trend toward increasing oxidative stress was observed at higher viral loads **(Fig. 2G); Fig. S5)**. Based on these results, a dilution of 1:1000 was selected, yielding approximately 20% GFP-positive cells per spheroid (∼ 10 cells per spheroid of 50). This fraction represents a balance between introducing a defined cellular perturbation and preserving network integrity during the maturation period preceding the amyloid-*β* perturbation.

Amyloid-*β* (A*β*) accumulation is a key pathological hallmark of AD (3), and was introduced as a global network perturbation at DIV 23 to model the diffuse nature of A*β* pathology in the brain. Immunofluorescence staining with the McSA1 antibody, which detects A*β* peptides (amino acids 1–12), confirmed the formation of perineuronal A*β* aggregates within the neurite network 14 days after addition **(Fig. 2H; Figs. S6–S7)**. A concentration titration was performed to determine a working concentration that allowed sustained study of network-level responses over prolonged periods **(Fig. 2I– K; Fig. S8)**. Aggregate count and average aggregate size increased significantly with concentration, while neurite area declined significantly at 1.0 µM and beyond **(Fig. 2I–K; Fig. S8)**. A working concentration of 0.2 µM was therefore selected for all subsequent experiments, as it induced detectable perineuronal A*β* aggregate formation while preserving network viability over the prolonged recording period.

Overall, progerin-induced cellular perturbation in a single node and a global A*β* perturbation established a defined perturbation paradigm within the 33-node network, which was subsequently interrogated using continuous HD-MEA recordings.

### Single-node progerin expression and global amyloid-β perturbation produce progressive, layer-dependent disruption of network firing dynamics

HD-MEA recordings were performed longitudinally from DIV 15 to 45 across both control and progerin/A*β* networks, spanning the period of network maturation, A*β* addition at DIV 23, and subsequent network evolution. To resolve spatiotemporal activity patterns and their changes in response to the combined perturbation, network responses were analysed both by layer and by hop distance from the progerin-expressing node, as illustrated in **Fig. 3A**. Hop distance is defined as the minimum number of nodes that must be traversed to reach a given node from the progerin-expressing node within the network topology.

**Figure 3.**
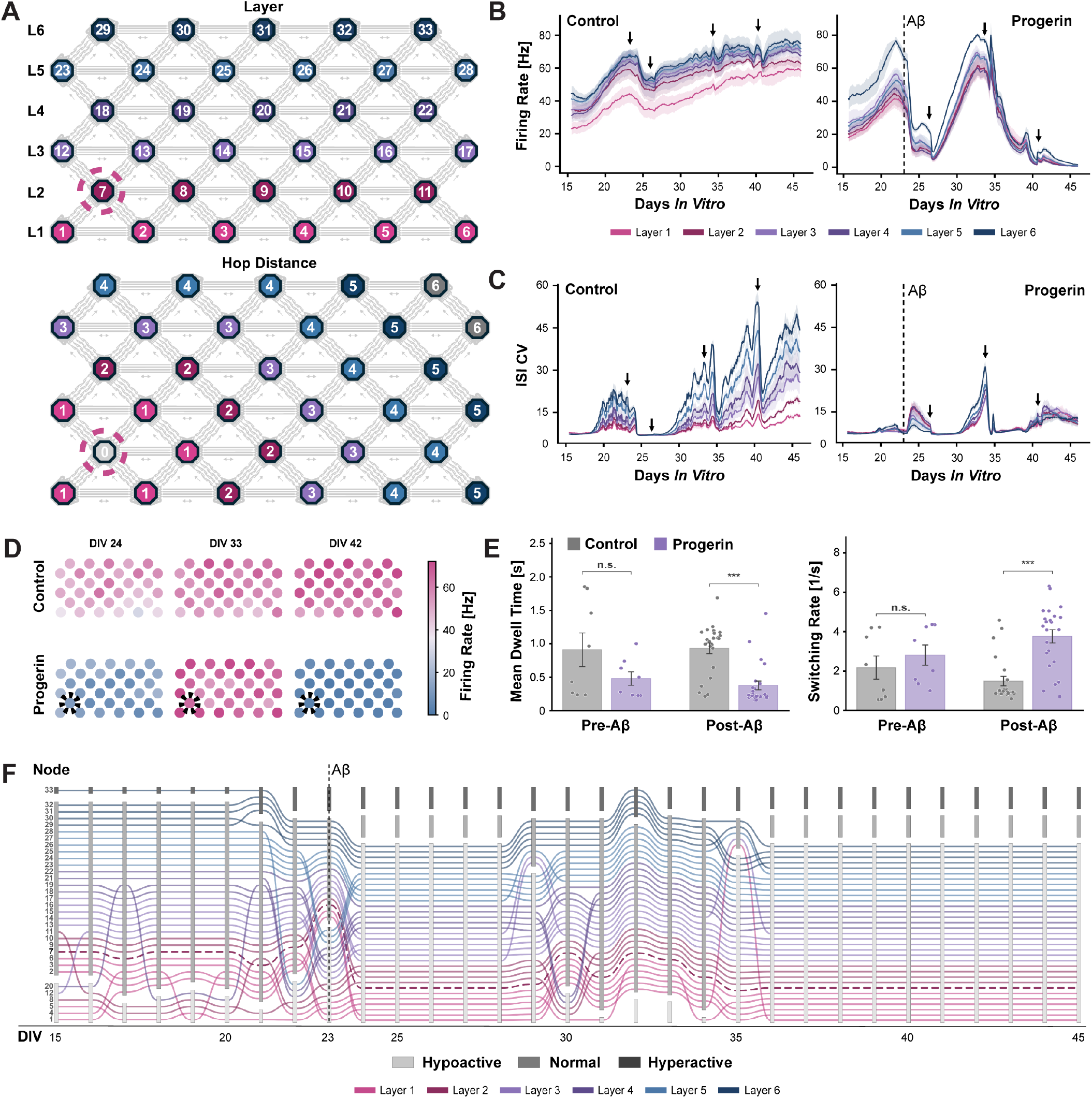
Single-node progerin expression and global amyloid-β perturbation produced progressive, layer-dependent disruption of network firing dynamics. **(A)** Schematics of the 33-node network architecture illustrating the two spatial metrics used to resolve network responses. Nodes are indexed 1–33, assigned starting from the lower-left node, increasing from left to right within each layer and continuing layer by layer from bottom to top. Top: nodes coloured by layer (1–6), with the progerin-expressing node indicated by a dashed circle. Bottom: nodes coloured by hop distance from the progerin-expressing node, defined as the minimum number of nodes separating a given node from the disease core. **(B)** Continuous firing rate trajectories for each layer (mean ± confidence interval) in control (left) and progerin/A*β* (right) networks from DIV 15 to 45. Both conditions exhibited a clear layer-dependent hierarchy, with higher layers displaying consistently elevated activity. A transient reduction following the media exchange at DIV 23, with A*β* added to the progerin network, was observed in both conditions, but was markedly more pronounced in the progerin/A*β* network. Firing rate in the progerin/A*β* network declined significantly after DIV 33. A dashed vertical line indicates the timing of A*β* addition. Arrows indicate media changes. **(C)** Longitudinal CV_ISI_ for each layer in control (left) and progerin/A*β* (right) networks. The control network showed a progressive layer-dependent increase in CV_ISI_ over time with transient reductions following media changes. The progerin/A*β* network displayed markedly reduced layer-dependent differentiation and a flatter trajectory. A dashed vertical line indicates the timing of A*β* addition. Arrows indicate media changes. **(D)** Node-level mean firing rate maps at DIV 24, 33, and 42 for control (top) and progerin/A*β* (bottom) networks. The dashed circle indicates the progerin-expressing node. Control networks showed a consistent network-wide increase in firing rate over time, while the progerin/A*β* network showed an initial increase followed by a progressive decline. **(E)** Mean dwell time (left) and switching rate (right) for control and progerin/A*β* networks pre- and post-A*β* addition. No significant difference was observed between conditions prior to A*β* addition. Following A*β* perturbation, the progerin/A*β* network exhibited significantly reduced mean dwell time and increased switching rate. \*\*\**p <* 0.001; ns, not significant. **(F)** Alluvial plot of firing rate classification for each node in the progerin/A*β* network over time (DIV 15–45). Nodes were classified as hypoactive, normal, or hyperactive based on the firing rate distribution of the control network. The y-axis denotes the node index (1–33), assigned starting from the lower-left node of the platform, increasing from left to right within each layer, and continuing layer by layer from bottom to top. Node lines are coloured by layer; the progerin-expressing node is indicated by a dashed line. Following A*β* addition, the network broadly transitioned into a hypoactive state, after which nodes in higher layers progressively recovered toward the normal range before the network returned to a broadly hypoactive state from DIV 35 onward. A dashed vertical line indicates the timing of A*β* addition.

Mean firing rate was computed as the per-electrode spike count divided by the duration of a recording block and averaged across the node’s electrodes. Each block had a length of 15 minutes. In control networks, the firing rate increased consistently across all layers throughout the recording period, with a clear layer-dependent hierarchy in which higher layers displayed persistently elevated activity **(Fig. 3B)**. This layer-dependent stratification was also present in the progerin/A*β* network prior to A*β* addition, suggesting that accelerated ageing in a single node does not disrupt the emergent hierarchical organisation of network activity during maturation. At DIV 23, both networks underwent a media change (with A*β* addition in the progerin/A*β* network), producing a transient reduction in firing rate that was markedly more pronounced in the progerin/A*β* network **(Fig. 3B)**. From DIV 33 onward, firing rate in the progerin/A*β* network declined significantly, with a concurrent reduction in the separation between layers compared to the control **(Fig. 3B–D)**.

The coefficient of variation of inter-spike intervals (CV_ISI_ = SD/mean of ISIs) is a firing-rate-independent measure of the irregularity of neural firing. In control networks, CV_ISI_ showed a progressive, layer-dependent increase over time, consistent with network maturation. Transient reductions in CV_ISI_ were observed following media changes, with recovery over the following hours **(Fig. 3C)**. In contrast, the progerin/A*β* network displayed markedly reduced layer-dependent differentiation in CV_ISI_ and a flatter trajectory over time. The stability of network activity states was further characterised using mean dwell time and switching rate, derived from a hidden Markov model fit to the network activity time series. Model selection and fit quality were validated using BIC, posterior state confidence, and heldout cross-validation, supporting K=3 as the optimal number of states **(Fig. S9)**. Dwell time quantifies how long the network remains in a given activity state, while switching rate reflects how frequently it transitions between states. Both showed no significant difference between conditions prior to A*β* addition **(Fig. 3E)**, but following A*β* perturbation, the progerin/A*β* network exhibited significantly reduced mean dwell time and increased switching rate compared to controls, indicating a shift toward more transient and less stable activity states. When resolved by layer, control networks showed a graded increase in dwell time and decrease in switching rate from lower to higher layers, whereas the progerin/A*β* network showed weaker layer dependence for both measures, particularly following A*β* addition **(Fig. S10)**. Burst frequency, burst amplitude, burst duration, and spike synchrony showed broadly consistent layer-dependent patterns, with progressive disruption observed in the progerin/A*β* network following A*β* addition **(Fig. S11)**. Together, these findings demonstrate that the combination of localised progerin expression and global A*β* perturbation produces progressive, layer-dependent disruption of network dynamics that is detectable longitudinally across the full six-layer hierarchy.

To track the evolution of individual node states over time, nodes in the progerin/A*β* network were classified at each timepoint as hypoactive, normal, or hyperactive relative to the control network firing rate distribution **(Fig. 3F)**. Thresholds were set at the control mean ± 2 SD per DIV, with nodes firing below 0.01 Hz considered silent and classified as hypoactive. Firing rate was computed per-electrode and averaged across each node’s electrodes, summarised as the per-DIV median. The elevated values relative to typical single-unit recordings likely reflect multi-unit activity captured by individual electrodes. Following A*β* addition, the network broadly transitioned into a hypoactive state. Over subsequent days, nodes in higher layers progressively recovered toward the normal activity range, while nodes in lower layers remained hypoactive for longer. From DIV 35 onward, the network returned to a broadly hypoactive state across all layers. This pattern is consistent with a spatially heterogeneous response to the combined disease perturbation, in which the distance from the progerin-expressing node, quantified as hierarchical layer and hop distance in the feedforward hierarchy **(Fig. 3A)**, may contribute to the observed spatial gradient of recovery. This gradient was statistically significant when quantified by layer throughout the recovery window (Spearman *ρ* = 0.23, *p* = 2.0 × 10−5), with a similar but weaker, non-significant trend by hop distance (*ρ* = 0.09, *p* = 9.0 × 10−2) **(Fig. S12)**.

### Transfer entropy reveals progressive reorganisation of functional connectivity and network topology following single-node progerin expression and amyloid-β Perturbation

Functional connectivity maps indicated predominantly feedforward directionality across layers in both control and progerin/A*β* networks, whereas within-layer connections showed more comparable transfer entropy in both directions. This was reflected in the sub-dominant/dominant transfer entropy ratio, which was low across layers (6.4 %, IQR 3.0 % to 14.1 %) but much higher within layers (53.1 %, IQR 33.4 % to 72.9 %). The asymmetry was consistent across conditions and recording timepoints **(Fig. 4A; Figs. S13–S14)**. Longitudinal tracking of TE asymmetry confirmed this hierarchical organisation, with lower layers predominantly acting as sources of information flow and higher layers as sinks, though nodes across all layers exhibited both behaviours over time **(Fig. 4B)**. This was consistent with the structural connectivity established by the platform design **(Fig. 1B–C)**, confirming that the asymmetric microchannel architecture successfully enforced directional information flow in both conditions. This hierarchical source-sink organisation mirrors the feedforward and feedback pathway structure described in primate visual cortex (33, 34).

**Figure 4.**
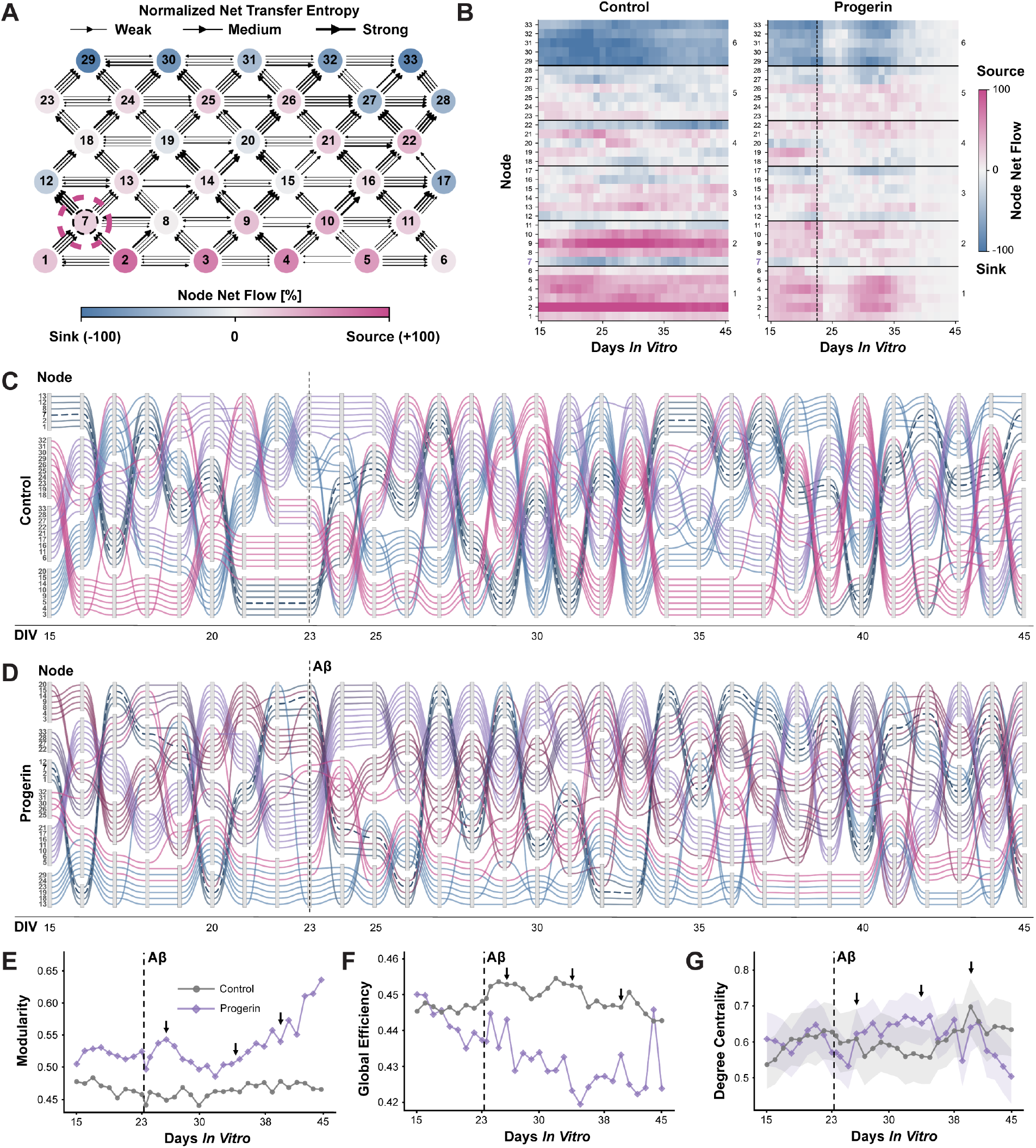
Transfer entropy revealed progressive reorganisation of functional connectivity and network topology in response to focal and global disease-relevant perturbations. **(A)** Representative snapshot of directional functional connectivity across the 33-node network at DIV 30, derived from normalised net transfer entropy (TE). Arrow thickness indicates the strength of net TE across each microchannel connection, and node colour indicates net information flow, with pink nodes acting as sources and blue nodes acting as sinks. The progerin-expressing node (node 7, dashed circle) is indicated. Inter-layer connections displayed predominantly strong feedforward directionality, while within-layer connections were more variable in direction and strength, consistent with the underlying structural connectivity. **(B)** Heatmaps of normalised net transfer entropy per node over time in control (left) and progerin/A*β* (right) networks, with nodes organised by layer (highest layers at top). Positive values (pink) indicated net source behaviour and negative values (blue) indicated net sink behaviour. Both networks displayed a complex source-sink organisation across all layers and timepoints, with lower layers tending toward net source behaviour and higher layers toward net sink behaviour, consistent with the feedforward hierarchy of the platform. **(C–D)** Alluvial plots of community structure over time for control **(C)** and progerin/A*β* **(D)** networks, derived using Louvain modularity. Node lines are coloured by layer and grouped into communities at each selected timepoint. Both networks maintained comparable numbers of communities, fluctuating between four and six, prior to A*β* addition. Qualitative inspection suggested more frequent reassignment of individual nodes between communities in the progerin/A*β* network compared to control following A*β* perturbation, potentially indicative of reduced community stability. **(E)** Modularity over time for control and progerin/A*β* networks. Modularity in the progerin/A*β* network was modestly elevated relative to control prior to A*β* addition. A steep increase was observed immediately following A*β* addition, followed by a decline coinciding with the media change, after which modularity increased progressively and remained markedly elevated compared to control through DIV 45. **(F)** Global efficiency over time for control and progerin/A*β* networks. Following A*β* addition, global efficiency in the progerin/A*β* network gradually declined over time relative to control. **(G)** Weighted degree centrality over time for control and progerin/A*β* networks (mean ± SEM). The two networks displayed comparable degree centrality prior to A*β* addition, after which the progerin/A*β* network showed a relative increase compared to control before declining below control levels from approximately DIV 38 onward. Media changes at DIV 26, 34, and 40 are indicated by arrows in **(E–G)**.

Community structure, referring to groups of nodes that communicate more strongly with each other than with the rest of the network, was assessed using Louvain modularity. Both networks maintained comparable numbers of communities, fluctuating between four and six, prior to A*β* addition **(Fig. 4C–D)**. Inspection of node community trajectories suggested more frequent reassignment of individual nodes between communities in the progerin/A*β* network compared to control **(Fig. 4C–D, Fig. S15)**, potentially indicative of reduced community stability. Interestingly, this apparent instability in the progerin/A*β* network coincided with a transient steep increase in modularity, a measure of the degree to which the network is organised into distinct communities. Modularity subsequently declined coincident with the media change before gradually increasing again and remaining markedly elevated compared to controls through DIV 45 **(Fig. 4E)**. Global efficiency, defined as the average inverse shortest path length between all node pairs and ranging from 0 to 1, gradually declined over time in the progerin/A*β* network relative to control **(Fig. 4F)**. Finally, degree centrality was quantified as the normalised, weighted total strength of a node’s incoming and outgoing connections to all other nodes. The two networks displayed comparable degree centrality prior to A*β* addition, after which the progerin/A*β* network showed a relative increase compared to control before declining toward comparable levels again around DIV 38 **(Fig. 4G)**.

Collectively, these findings demonstrate that nodal progerin expression and global A*β* perturbation produce progressive and spatially heterogeneous reorganisation of functional network topology, detectable simultaneously at the level of individual nodes, communities, and whole-network graph metrics. The 33-node architecture makes it possible to resolve how individual nodes transition between functional states over time.

### Multiparametric classification reveals progressive network divergence following nodal accelerated ageing and amyloid-*β* Perturbation

To characterise the multi-parametric electrophysiological signature distinguishing control from progerin/A*β* network nodes, t-Distributed Stochastic Neighbor Embedding (t-SNE) was applied to feature vectors extracted from all network nodes at each recorded timepoint **(Fig. 5A)**. At DIV 15, control and progerin/A*β* nodes occupied overlapping regions of the t-SNE space, suggesting broadly similar electrophysiological profiles during early network maturation. Progressive separation between conditions emerged from DIV 20 onward, prior to A*β* addition at DIV 23, with control and progerin/A*β* nodes occupying increasingly distinct regions of the feature space over time. By DIV 35–40, the two conditions were almost completely separated, with progerin/A*β* nodes forming a tight cluster distinct from the control nodes. Within the progerin/A*β* network, nodes at higher hop distances from the progerin-expressing node tended to occupy positions closer to the control cluster at later timepoints **(Fig. 5A; Figs. S16–S17)**, suggesting a spatial gradient in the electrophysiological divergence from controls that decays with increasing topological distance from the progerin-expressing node. This spatial structuring of feature space could be quantified directly, with pairwise feature-vector distances correlating with both layer and hop distance from the progerin-expressing node **(Fig. S18)**.

**Figure 5.**
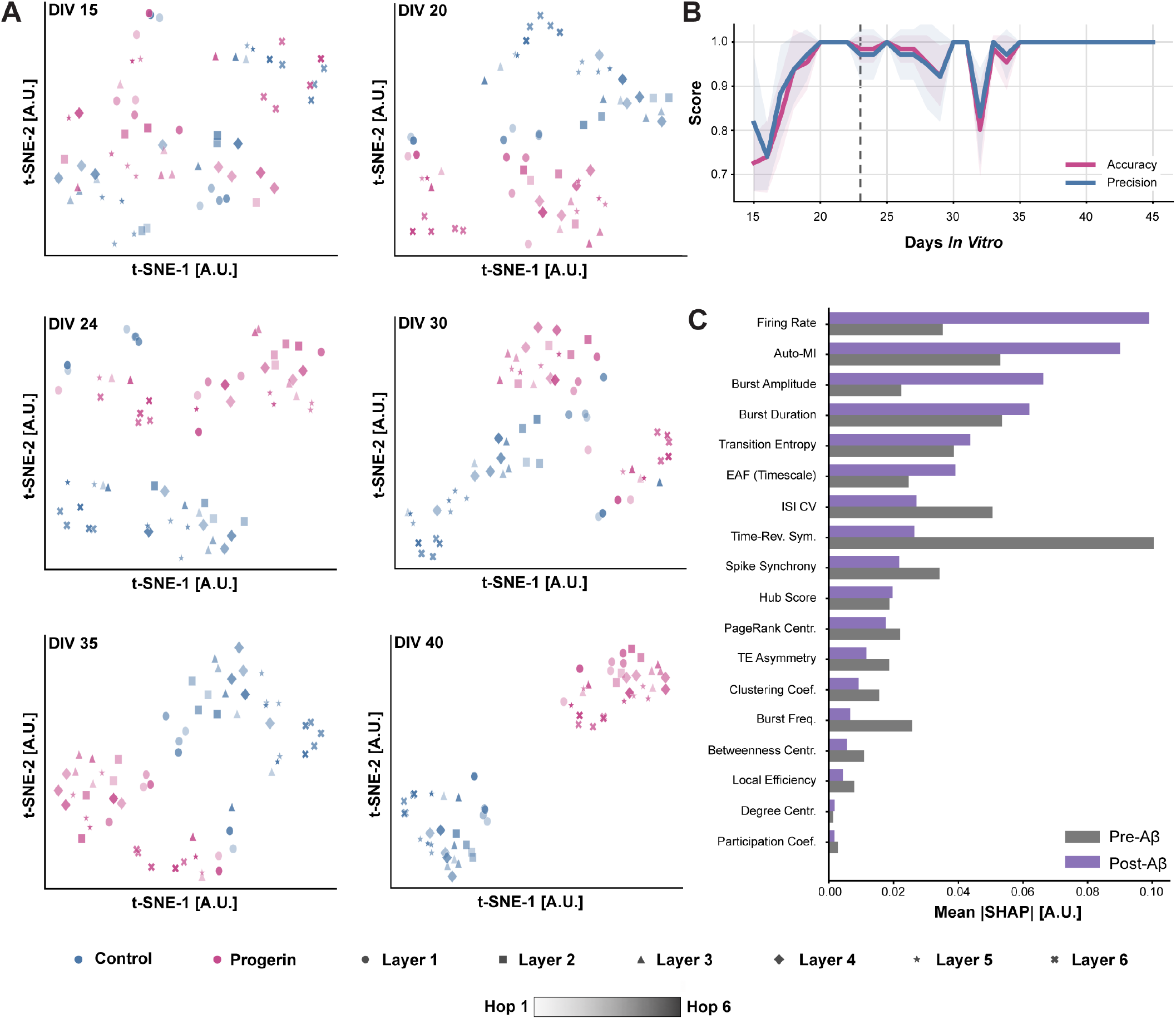
Multiparametric electrophysiological classification distinguishes control from progerin/A*β* network nodes and reveals progressive divergence over time. **(A)** t-SNE projections of multiparametric electrophysiological feature vectors for all network nodes at six timepoints (DIV 15–40), coloured by condition (blue: control, pink: progerin/A*β*) with colour saturation indicating hop distance from the progerin-expressing node (lighter: lower hop distance, darker: higher hop distance), and shaped by layer (Layer 1–6). Progressive separation between conditions emerged from DIV 20 onward, preceding A*β* addition at DIV 23, and became increasingly pronounced over time. At later timepoints, progerin/A*β* nodes at higher hop distances tended to occupy positions closer to the control cluster, suggesting a spatial gradient in the electrophysiological divergence from controls that decays with increasing topological distance from the progerin-expressing node. **(B)** Longitudinal classification performance (accuracy and precision) of a per-DIV stratified 5-fold cross-validated Random Forest classifier distinguishing control from progerin/A*β* nodes (mean ± SD across folds). Classification performance exceeded chance from DIV 15 onward, reached near-perfect accuracy by DIV 20, and recovered rapidly following transient dips coinciding with media changes at DIV 23, 26, 34, and 40. Dashed vertical line indicates timing of A*β* addition. **(C)** SHAP feature importance for a Random Forest classifier trained to distinguish control from progerin/A*β* nodes, shown separately for the pre-A*β* (DIV *<* 23, grey) and post-A*β* (DIV ≥ 23, purple) periods. Firing rate, Auto-MI, and burst-related features dominated post-A*β*, while time-reversal symmetry was the most prominent feature pre-A*β*.

The separability over time was investigated by training a Random Forest classifier, which distinguishes control from progerin/A*β* nodes. First, the longitudinal classification performance confirmed that control and progerin/A*β* nodes were distinguishable throughout the recording period **(Fig. 5B)**. Classification accuracy and precision exceeded chance level from DIV 15 onward, reaching near-perfect performance by DIV 20 and remaining consistently high thereafter. Transient dips in classification performance were observed coinciding with media changes at DIV 23, 26, 34, and 40, consistent with the transient perturbations in network dynamics observed in the firing rate and CV_ISI_ analyses **(Fig. 3B– C)**. Following each dip, classification performance recovered rapidly, and from DIV 37 onward, all metrics reached and maintained perfect or near-perfect scores. Together, these findings demonstrate that the multiparametric electrophysiological signature of the progerin/A*β* network diverges progressively from controls over time, is detectable prior to A*β* addition, and becomes increasingly pronounced following the combined perturbation.

The importance of distinct features was analysed by applying Shapley Additive Explanations (SHAP) to the Random Forest classifier. It revealed distinct feature importance profiles before and after A*β* addition **(Fig. 5C; Fig. S19)**. Prior to A*β* addition, time-reversal symmetry was the most prominent feature, alongside burst duration, Auto-MI and CV_ISI_. Following A*β* addition, firing rate, auto-mutual-information (Auto-MI), and burst duration and amplitude became the dominant features, reflecting the more pronounced and widespread changes in network dynamics observed after the A*β* perturbation.

## Discussion

Understanding how pathology spreads through the hierarchical circuits of the human brain requires models that combine architectural complexity with the resolution to track activity across every connection in the network. Here, a 33-node, six-layer human neural circuit platform integrated with continuous HD-MEA recordings was developed to address this challenge. Spheroids of 50 cells were assembled across 33 defined nodes, with electrodes routed within every connecting microchannel, enabling activity to be tracked at high spatiotemporal resolution across a known structural scaffold continuously over weeks. Transfer entropy-based functional connectivity analysis, graph-theoretic mapping, and machine learning-based feature analysis enabled detailed characterization of disease-induced network dynamics. Beyond demonstrating that disease-relevant perturbations alter network dynamics, these approaches resolved precisely where, when, and in what order these changes emerge across the full circuit hierarchy.

Structural characterisation confirmed strong directional axonal outgrowth across inter-layer connections, consistent with the Tesla valve geometry. Fluorescence imaging revealed a significant bias toward feedforward axonal outgrowth, though neurites of both fluorescent labels were observed within inter-layer channels. This may reflect the spontaneous formation of long-range projections spanning multiple layers, which are well established in the neocortex and are thought to play important functional roles in hierarchical information processing (35–37). Transfer entropy analysis confirmed that functional connectivity was strongly directional and hierarchically organised across the six layers, consistent with the structurally imposed feedforward architecture, validating the platform as a model of feedforward cortical circuit organisation.

The combined perturbation produced a temporally ordered sequence of network-level changes. Before A*β* addition, signatures of focal cellular stress were already detectable. In neurodegeneration, networks are proposed to initiate compensatory reorganisation early to preserve function before pathological burden overwhelms these mechanisms (5, 38, 39). Consistent with this, multiparametric classification confirmed early divergence between conditions, with control and progerin/A*β* nodes distinguishable from DIV 15 onward and near-perfect separation achieved by DIV 20. The dominant pre-A*β* feature was time-reversal symmetry, a measure of the degree to which network dynamics deviate from thermodynamic equilibrium (40, 41). Its prominence suggests that progerin expression may alter the temporal structure of network activity in subtle ways that precede overt changes in firing rate or burst properties, consistent with evidence that reduced temporal irreversibility is an early signature of AD-relevant network dysfunction (42).

Following A*β* addition, the layer-dependent firing rate hierarchy progressively collapsed, and layer-dependent CV_ISI_ differentiation was similarly lost, suggesting a breakdown of the distinct activity profiles that each layer had developed during maturation (5, 39). Correspondingly, firing rate, Auto-MI, and burst-related features emerged as the dominant classifiers, reflecting the more pronounced changes in network dynamics following the global perturbation. This is consistent with the observed tendencies toward transient hyperactivity and increased bursting followed by hypoactivity in the perturbed networks, mirroring trajectories reported following amyloid accumulation in both transgenic animal models and iPSC-derived systems (4, 43, 44). Aberrant bursting has been closely linked to amyloid-driven network dysfunction, where reduced inhibitory signaling and disrupted glutamate clearance drive early network hyperexcitability, before progressive synaptic loss leads to network silencing (45). Critically, this disruption was spatially heterogeneous. Nodes in higher layers recovered toward normal activity ranges before those in lower layers, and nodes at higher hop distances from the progerin-expressing node tended to cluster closer to controls in multiparametric feature space at later timepoints. This spatial gradient of recovery is consistent with topological distance from the disease core conferring a degree of resilience (38, 39, 46). Whether this reflects a specific protective effect of distance or simply the higher baseline activity of upper layers remains an open question for future studies with replicate networks.

Graph-theoretic analysis indicated a pattern consistent with adaptive reorganisation and progressive failure. A transient increase in modularity was observed immediately following A*β* addition, suggesting an initial compensatory attempt to maintain network segregation (47, 48), while global efficiency gradually declined over the recording period, reflecting a progressive loss of network integration. This was accompanied by more frequent reassignment of individual nodes between communities in the progerin/A*β* network relative to control, suggesting reduced temporal stability of network organisation, consistent with the disease-related reorganisation of community allegiance reported in AD functional brain networks (49). In neurodegeneration, this pattern has been interpreted as reflecting a process in which network communication becomes increasingly rerouted through a small number of highly connected hub nodes, placing growing metabolic demands on them and ultimately predisposing them to failure (50, 51). Notably, mean weighted degree centrality was comparable between conditions prior to A*β* addition but became transiently elevated in the progerin/A*β* network following A*β* addition, before converging toward control levels by approximately DIV 38, consistent with a period of increased connectivity strength coinciding with the compensatory phase of network reorganisation. Whether this reflects selective hub formation or a broader increase in connectivity density cannot be resolved from mean degree centrality alone, and remains an open question for future work.

The present study represents a proof-of-principle with a single network per condition. The observed differences between conditions are considerable, but require validation in replicate networks to establish reproducibility. The per-DIV stratified cross-validation used for classification is performed within a single chip and therefore validates the classifier’s ability to distinguish nodes within that recording rather than generalising across independent biological replicates. Furthermore, while progerin expression is a well-validated model of accelerated neural ageing (30, 31), it recapitulates specific aspects of pathological ageing rather than the full complexity of physiological neural senescence. Modularity optimisation is additionally subject to a resolution limit, whereby small communities may be merged into larger ones (52), which should be considered when interpreting the community structure results reported here. Transient reductions in firing rate and CV_ISI_ were also observed in both conditions following media changes, with recovery in subsequent recordings, likely reflecting changes in temperature during the exchange procedure. Notably, the magnitude of these transient perturbations was modest, consistent with the stable recording environment maintained by the Inkudock system (53, 54). Between media changes, network activity remained stable across consecutive recording blocks, demonstrating the suitability of the platform for longitudinal electrophysiological studies over extended periods.

This platform establishes a scalable framework for spatially resolved, longitudinal interrogation of disease-relevant network dynamics in human neural circuits. Natural extensions include the use of patient-derived iPSC neurons to study disease-specific genetic backgrounds, layer-specific neural subtypes to more faithfully recapitulate cortical laminar organisation, and pharmacological perturbations to enable direct assessment of therapeutic interventions. Beyond pharmacological perturbations, the platform is amenable to additional disease-relevant stressors such as inflammatory stimuli, oxidative stress, and combinations of proteinopathies, enabling systematic dissection of how multiple pathological factors interact at the circuit level. The approach is broadly applicable to neurological conditions where focal pathology propagates through interconnected neural circuits, including Parkinson’s disease, ALS, and other conditions characterised by the combination of ageing and proteinopathy (5, 51). The ability to track how individual nodes transition between functional states across a known circuit hierarchy offers a route toward identifying the earliest electrophysiological signatures of neurodegeneration *in vitro* and testing therapeutic interventions aimed at halting disease progression.

## Materials and Methods

### Experimental Design

The experimental workflow is illustrated in **Figure 6A** and the timeline in **Figure 6D**. Excitatory neurons, inhibitory neurons, and astrocytes were seeded into AggreWell plates to form uniform spheroids comprising 25 excitatory neurons, 25 inhibitory neurons, and 10 astrocytes. One spheroid per network was transduced with a lentiviral vector at a 1:1000 dilution to induce progerin expression in approximately 20% of cells, introducing a site of locally accelerated ageing within the network. Following a 24-hour aggregation period, spheroids were transferred into the 33-node microfluidic platform pre-mounted on Maxwell Max-One Plus PLM HD-MEA chips (26,400 electrodes), with the progerin-expressing spheroid seeded into a designated node in layer 2 of the feedforward hierarchy. Networks were allowed to mature for 15 days prior to the onset of continuous electrophysiological recordings.

**Figure 6.**
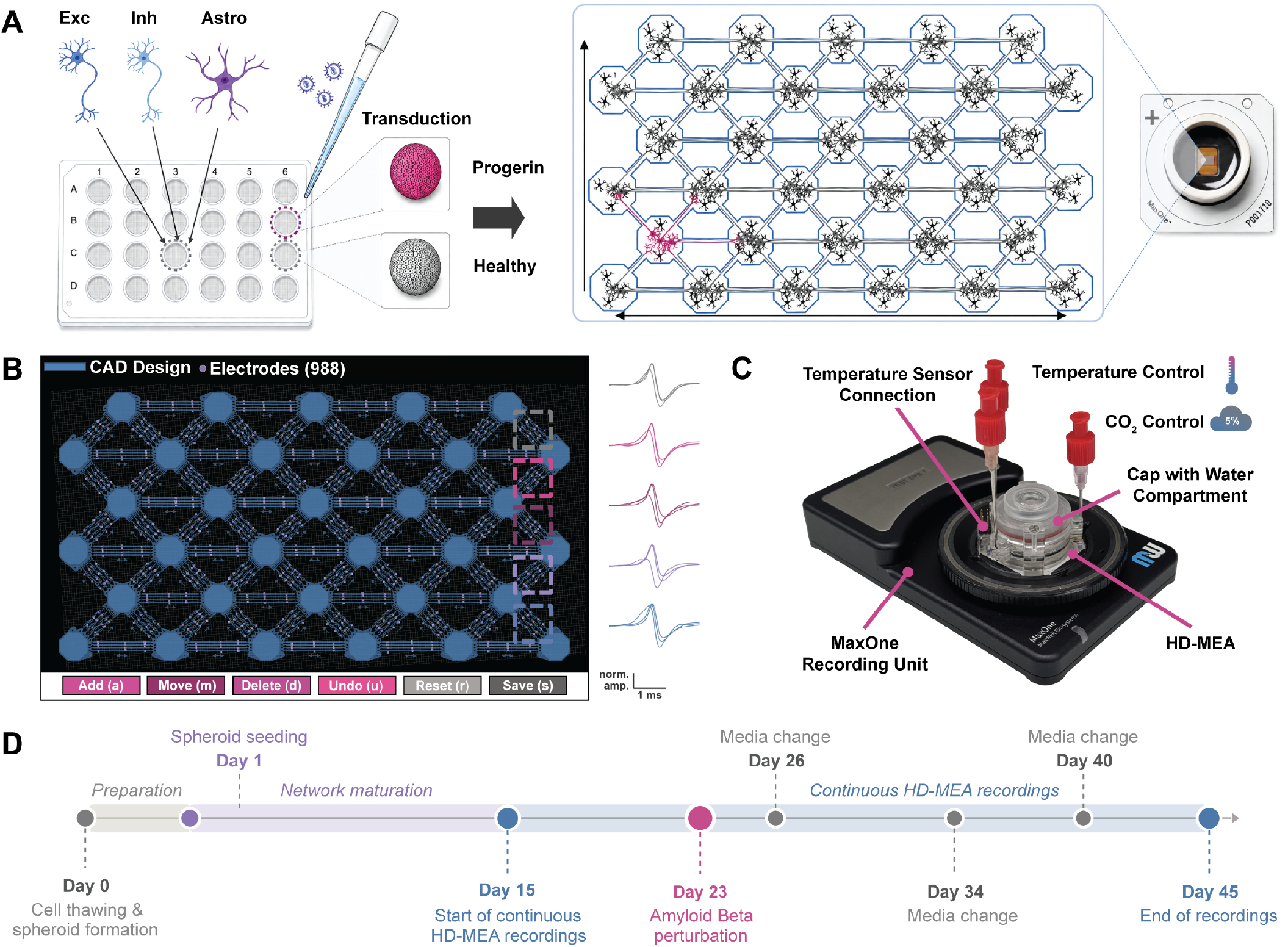
Integrated experimental workflow for longitudinal HD-MEA recording of a 33-node human neural circuit model of AD-relevant neurodegeneration. **(A)** Overview of the experimental workflow. Excitatory neurons, inhibitory neurons, and astrocytes were seeded into AggreWell plates to form uniform spheroids of 50 cells, comprising 25 excitatory neurons, 25 inhibitory neurons, and 10 astrocytes. One spheroid was transduced with a lentiviral vector to induce progerin expression (pink). Spheroids were transferred into the 33-node microfluidic platform, with the progerin-expressing spheroid seeded into a designated node (pink, dashed circle). These networks were cultured on a Maxwell HD-MEA chip (26,400 electrodes) for longitudinal electrophysiological recording. **(B)** Automated electrode selection interface, showing the CAD design of the 33-node layout overlaid on the HD-MEA electrode array. Three electrodes per microchannel (proximal, central, and distal) were placed automatically using a custom pipeline, yielding 924 routed electrodes per chip. The GUI allowed manual inspection and correction of electrode positions prior to recording. Representative spike waveforms from selected electrodes are shown. **(C)** Environmental control setup for longitudinal recordings as introduced and validated in previous work (53). The HD-MEA chip was kept in a CO_2_-controlled chamber at 5% CO_2_. The temperature was maintained at around 36°C in the cell culture medium during recording, accounting for heat dissipation from the active electronic circuitry of the HD-MEA. The medium temperature was selected to prevent condensation on the cell culture lid during day-long recordings by ensuring a warmer culture lid with respect to the medium. A cell culture lid with a water compartment at the interface was used to address the humidity gradient and thus minimise evaporation of cell culture medium, which is required to prevent drifts in ionic concentrations. **(D)** Experimental timeline from cell thawing to end of recordings. Spheroids were formed and seeded on Day 0–1, with continuous HD-MEA recordings initiated at DIV 15 following a network maturation period. Amyloid-*β* perturbation was applied to the progerin/A*β* network at DIV 23, with media changes performed at DIV 26, 34, and 40. Recordings were concluded at DIV 45.

Electrode selection was performed using a custom automated pipeline that placed three electrodes per microchannel, yielding 924 routed electrodes per chip, with manual inspection and correction performed via a custom graphical user interface (GUI) prior to recording **(Fig. 6B)**. Recordings were maintained under controlled environmental conditions at 36 ^◦^C and 5% CO_2_ using the Inkudock system combined with a MEA cap including an intermediate water compartment to eliminate evaporation **(Fig. 6C)** (53, 54).

Continuous HD-MEA recordings were initiated at DIV 15 and maintained until DIV 45. At DIV 23, amyloid-*β* was added globally to the progerin/A*β* network, while the control network received a media change only. Subsequent media changes were performed at DIV 26, 34, and 40 for both networks. With each media change, the water compartment of the cap was refilled.

### Microstructure Design and Fabrication

PDMS microstructures were designed using AutoCAD (Autodesk) and fabricated by WunderliChips GmbH (Zurich, Switzerland). The platform comprised 33 spheroid chambers arranged in a six-layer feedforward hierarchy, with four parallel microchannels connecting each adjacent chamber pair. Chamber openings were 200 µm in diameter, with a chamber depth of approximately 200 µm. Inter-layer unidirectional microchannels were 300 µm in length and 4 µm in height, while within-layer bidirectional channels were 450 µm in length at the same height. The platform was designed to fit within the active electrode area of the Maxwell MaxOne HD-MEA (3.85 × 2.1 mm), with channel spacing accounting for the 17.5 µm electrode pitch to ensure that individual microchannels did not span more than one electrode column.

Directional connectivity between layers was enforced using Tesla valve-inspired microchannel geometries, which redirect neurites growing in the non-preferred direction back toward their chamber of origin (8). Sawtooth structures were integrated on the postsynaptic side of unidirectional channels to further misdirect neurites attempting to enter the channels from the unintended direction. Droplet-shaped pillars were placed at the channel entrances to provide structural support for the PDMS roof and to guide neurites toward the channel entrances rather than allowing them to grow around the pillar structures.

### Chip Preparation

Prior to cell seeding, PDMS microstructures were assembled onto either Maxwell MaxOne Plus PLM HD-MEA chips or glass-bottom dishes (Ibidi, catalog no. 81218-200), depending on the experiment.

For glass-bottom dishes, surfaces were plasma-treated for 60 s using a Tergeo plasma cleaner (PIE Scientific, USA) with an air/O_2_/H_2_O program at 35 W with a 50% duty cycle, an oxygen flow rate of 5 sccm and a water flow rate of 10 sccm. Dishes were immediately transferred to a biosafety cabinet and UV-sterilized for 20–30 min. Poly-D-lysine (PDL, Gibco, A3890401, 0.1 mgmL^−1^) was applied to cover the surface and incubated for 45 min at room temperature, after which surfaces were rinsed three times with ultrapure water and dried under nitrogen flow.

For Maxwell MaxOne Plus PLM HD-MEA chips, surfaces were rinsed three times with ultrapure water and dried under nitrogen flow prior to PDL coating at the same concentration and incubation conditions as described above.

In both cases, PDMS microstructures were carefully placed onto the coated surface and secured by desiccation for 2 min to promote adhesion. Surfaces were then filled with prewarmed phosphate-buffered saline (PBS) and desiccated for a further 10–15 min until no visible air bubbles remained within the microstructures. PBS was subsequently replaced with neural medium and substrates were stored in a humidified incubator at 37 ^◦^C and 5% CO_2_ for a minimum of 2 h prior to cell seeding.

### Spheroid Formation and Seeding

Three distinct spheroid compositions were used depending on the experiment. For quantification of axon directionality and neurite width, spheroids comprised NGN2-induced excitatory neurons expressing either GFP or RFP, seeded in alternating layers. For optimisation of amyloid-*β* perturbation conditions, spheroids comprised NGN2-induced excitatory neurons only. For all disease model experiments, composite spheroids were formed as described below.

Spheroids of 50 cells were formed using AggreWell™400 96-well plates (StemCell Technologies, catalog no. 200-0563). Human iPSC-derived excitatory neurons, inhibitory neurons, and astrocytes were thawed and co-seeded to form spheroids comprising 25 excitatory neurons, 25 inhibitory neurons, and 10 astrocytes per spheroid.

Human iPSC-derived excitatory neurons were kindly provided by Novartis, generated using a doxycycline-inducible NGN2 differentiation protocol (55) and cryopreserved in FBS with 5% DMSO prior to delivery. Human GABAergic inhibitory neurons were generated using a single-step transposon-based induction protocol with Ascl1/Dlx2 (56), and were kindly provided by Prof. L. Niels Cornelisse and Juraj J. Ondris at Amsterdam UMC. Human astrocytes were generated using the Sigma-iSOX9 cell line (57), and were kindly provided by Prof. Catherine Verfaillie and Dr. Karan Ahuja at KU Leuven. All cell types were stored in liquid nitrogen until use.

Cells were cultured in neural medium consisting of Neurobasal Plus Medium (Gibco™, A3582901) supplemented with 2% B27 Plus (Gibco™, A3582801), 1% N2 (Gibco™, 17502048), 0.1% BDNF (PeproTech, 450-02), 0.1% GDNF (PeproTech, 450-10), 1% GlutaMax (Gibco™, 35050061), and 1% Penicillin-Streptomycin (Sigma-Aldrich, P4333). For composite spheroids, the medium was additionally supplemented with 0.1% CNTF (PeproTech, 450-13) throughout the culture period, and Fetal Bovine Serum (FBS, Sigma-Aldrich, F9665) was added to a final concentration of 0.5% during seeding only. Rock Inhibitor (Sigma-Aldrich, Y27632) was added at 0.1% during seeding to enhance cell viability. Doxycycline (Fisher Scientific, NC0424034) was added at 0.2% up to DIV 10 to support completion of NGN2-induced neural differentiation. From DIV 4 onward, half the medium volume was replaced with fresh neural medium every 3–4 days.

After 24 hours in the AggreWell plates, spheroids were transferred to the microstructure chambers. Spheroids were first dissociated from the AggreWell substrate by gentle pipetting and transferred to the hydrophobic surface of a petri dish lid to prevent attachment. Individual spheroids were then selected and deposited into the individual chambers of the microstructure using a 10 µl pipette under a stereomicroscope in a biosafety cabinet.

### Induction of Progerin Expression

To induce progerin expression, cells were transduced using a lentiviral vector carrying a GFP-tagged progerin construct (pHR-SIN-GFP-PG, KC-CL-101), allowing GFP fluorescence to serve as a reporter for successful transduction, as previously described (32). Lentiviral particles were produced using HEK293T cells transiently transfected with the transfer plasmid alongside packaging plasmids (CMV-Gag-Pol, Harvard dR8.91) and envelope plasmid (pVSV-G, Clontech 631530). Viral supernatants were collected 48 h post-transfection, clarified by centrifugation at 500*g* for 10 min at 4°C, filtered through a 0.45 µm membrane, and concentrated 10-fold overnight at 4°C using Lenti-X Concentrator (Takara, 631232). Concentrated viral particles were pelleted at 1500*g* for 45 min at 4°C, resuspended in ice-cold PBS, and stored as aliquots at − 80°C until use. For optimisation of transduction efficiency, a serial dilution of lentiviral particles was tested across dilutions of 1:100, 1:200, 1:500, 1:1500, and 1:5000. Based on these results, a final dilution of 1:1000 was selected, yielding approximately 20% GFP-progerin-positive cells per spheroid (**Fig. 2D–G)**. Twenty-four hours after seeding in AggreWell plates, half the medium volume was replaced with neural medium containing the lentivirus at twice the target concentration. Following a 3-hour incubation, the virus-containing medium was replaced with fresh neural medium.

### Amyloid Beta Perturbations

Beta-Amyloid Peptide (1-42, human) (Abcam, ab120301) was prepared as follows. A vial of 100 µg lyophilized peptide was rapidly thawed and dissolved in 100 µL of 1% NH_4_OH (Thermo Scientific, 458680025) in PBS, before further dilution to 1 mgmL^−1^ with 900 µL PBS. The solution was vortexed gently for 30 s to mix without inducing aggregation, yielding a stock concentration of 22.153 µM. The solution was aliquoted into 20 µL aliquots, snap-frozen in liquid nitrogen, and stored at − 80 ◦C until use (58).

To determine an appropriate working concentration, a titration of A*β* concentrations was performed, as described in **Fig. 2I–K**. Titration experiments were performed in 18-well Ibidi chips (Ibidi, catalog no. 81818). Based on these results, a final working concentration of 0.2 µM was selected to induce A*β* aggregate formation while preserving network viability over prolonged recording periods. At DIV 23, aliquots were thawed and diluted in neural medium to the target concentration, and half the medium volume in the progerin/A*β* network was replaced with the A*β*-containing medium. The control network received a media change only. Immunocytochemistry was conducted 18 days after perturbation to assess perineuronal A*β* aggregate formation **(Fig. 2H)**.

### Immunocytochemistry & Confocal Microscopy

Cells were fixed with 4% paraformaldehyde (PFA) for 15 min at room temperature, followed by three 5-min washes with phosphate-buffered saline (PBS, Gibco™, 10010-015). Permeabilization was performed using 0.5% Triton X-100 (Sigma-Aldrich, 066K0089) in PBS for 5 min, after which cells were washed twice with PBS for 5 min each. Nonspecific binding was blocked by incubating samples in 5% goat serum (Abcam, ab7481) in PBS for 1 h at room temperature under gentle shaking at 50 rpm. Cells were then incubated overnight at 4 ^◦^C with primary antibodies diluted in fresh blocking solution (see **Table 1** for antibodies and concentrations). The following day, primary antibody solution was removed and samples were washed three times with PBS for 5 min each. Secondary antibodies diluted in blocking solution were then applied for 3 h at room temperature under gentle shaking, protected from light. Nuclei were counter-stained with Hoechst 33342 (1:2000; Invitrogen, H3570) in PBS for 30 min at room temperature, protected from light. Samples were washed three times with PBS for 5 min each, once with ultrapure water, and immersed in fresh ultrapure water for imaging.

**Table 1.** Antibodies and concentrations used for Immunocytochemistry.

| Marker | Catalogue Number | Concentration |
| --- | --- | --- |
| β3-Tubulin | ab41489 | 1/500 |
| Synapsin | ab254349 | 1/200 |
| VGAT | 131 011 (Synaptic Systems) | 1/200 |
| McSA1 | MM-0015-P (MédiMabs) | 1/1000 |
| Goat anti-Rabbit 488 | A32731 (Invitrogen) | 1/1000 |
| Goat anti-Mouse 555 | A32727 (Invitrogen) | 1/1000 |
| Goat anti-Chicken 647 | ab150171 | 1/1000 |

*All antibodies were purchased from Abcam unless otherwise stated.*

All images were acquired using an Olympus™ FV3000 Confocal Laser Scanning Microscope equipped with solid-state lasers at 405, 445, 488, 561, 594, and 640 × nm. Images were acquired using either a 20× objective (UPLFLN20XPH, NA 0.5) or a 60 × oil-immersion objective (UPLSAP060XS2, NA 1.3).

### Analysis of Neurite Directionality

Fluorescence images were processed using custom ImageJ macros. Regions of interest (ROIs) were drawn around the four microchannels in each image, and intensity and neurite width profiles were extracted for each lane along the full channel axis.

For intensity profiles, the maximum pixel value across the lane width was sampled at each axial position. Intensity values were normalised to the global per-image, per-channel maximum to account for variability in fluorescence intensity between imaging sessions and differences in fluorophore brightness or imaging gain settings.

For neurite width profiles, each lane was contrast-enhanced using CLAHE and binarised using Otsu thresholding. The Local Thickness plugin was applied to the binary mask, assigning to each foreground pixel the diameter of the largest inscribed circle at that location. The maximum local thickness across the lane width was sampled at each axial position to yield a width profile in micrometres.

Profiles were averaged across the four lanes per image and summarised over the 50–200 µm region from the channel entry. Statistical comparisons between desired and undesired directions were performed using independent-samples *t*-tests.

### Quantification of Amyloid-*β* Deposits and Neurite Morphology

Quantification of perineuronal A*β* deposits and neurite morphology was performed on confocal images acquired at 60 × magnification, using McSA1 and *β*III-tubulin staining respectively. Images were processed using custom ImageJ macros and converted to 8-bit format prior to analysis.

For A*β* deposit quantification, McSA1 images underwent rolling-ball background subtraction, Gaussian filtering, contrast optimisation, adaptive local thresholding using the Bernsen method (radius = 15 pixels, contrast threshold = 30), and hole filling. Images were converted to binary masks and particle density, aggregate count, and average aggregate area were measured using the Fiji Analyse Particles plugin.

For neurite morphology quantification, *β*III-tubulin images underwent rolling-ball background subtraction, contrast optimisation, non-linear gamma correction, Gaussian filtering, local contrast enhancement (CLAHE), Tubeness filtering, and Otsu thresholding. Images were converted to binary masks and neurite area was evaluated using the Fiji Analyse Particles plugin. Binary masks were further processed through the Skeletonize plugin and analysed using the Analyse Skeleton (2D/3D) plugin to quantify total neurite length, endpoint density, and junction number.

Data analysis and visualisation were performed in RStudio. Statistical comparisons across A*β* concentrations were performed using one-way ANOVA followed by Tukey’s post hoc test, with comparisons made against the control group only. Results are reported as mean ± SEM.

### Analysis of Progerin Expression

Progerin expression was evaluated across lentiviral dilutions of 1:100, 1:200, 1:500, 1:1500, and 1:5000 in spheroids of 50 excitatory neurons seeded in 18-well Ibidi chips (Ibidi, catalog no. 81818). At DIV 7, total nuclei were labelled using a LIVE/DEAD™ Cell Imaging Kit (488/570; Invitrogen, R37601). At DIV 9– 10, GFP-progerin-expressing spheroids were imaged by confocal Z-stack acquisition with a 5 µm step size.

Images were processed using custom ImageJ macros. Both the nuclear channel and GFP channel underwent a uniform preprocessing pipeline consisting of rolling-ball background subtraction, contrast optimisation, Gaussian filtering, Otsu thresholding, hole filling, and watershed segmentation. The percentage of GFP-progerin-positive cells was calculated relative to the total number of nuclei per spheroid. Data analysis and visualisation were performed in RStudio. Statistical comparisons were performed using one-way ANOVA followed by Tukey’s post hoc test, with comparisons made against the control group only.

### Analysis of Reactive Oxygen Species

Oxidative stress was assessed at DIV 35 in progerin-expressing spheroids using the CellROX™ Deep Red reagent (Invitrogen, C10422). Culture medium was replaced with neural medium containing 5 µM CellROX and incubated for 30 min, after which the staining solution was replaced with fresh phenol red-free neural medium. Live imaging was performed using an Olympus™ FV3000 CLSM with a 30 × silicone oil-immersion objective (UPLSAP030XS, NA 1.05).

CellROX signal was quantified as integrated density normalised to spheroid area. The CellROX channel was preprocessed using background subtraction. Spheroid area was determined from the transmitted light channel using the Fiji Analyse Particles plugin following background subtraction, Gaussian filtering, Otsu thresholding, hole filling, erosion, and dilation. Data analysis and visualisation were performed in RStudio. Statistical comparisons were performed using one-way ANOVA followed by Tukey’s post hoc test, with comparisons made against the control group only.

### Electrode Selection Pipeline

The MaxOne Plus chip features 26,400 electrodes on a fixed grid, but only up to 1,020 can be recorded simultaneously. A subset of recording electrodes must therefore be chosen per chip. An automated electrode selection pipeline that chooses electrodes inside every microchannel of the 33-node layout without manual intervention was developed **(Fig. S20)**. The pipeline takes two inputs: (i) the chip’s measured impedance map (exported as a 120 × 220 array), which reveals the microstructure features on the electrode grid (59), and (ii) the computer-aided design (CAD) layout of the chamber-microchannel geometry, supplied as a GDS file. It consists of three stages: registration, geometry segmentation, and selection. These stages can be run end to end, while each is implemented as an independent module, allowing user intervention after every step.

### Registration

The impedance map produced by the HDMEA is a two-dimensional grid of values, one per recording electrode, and can therefore be treated as a greyscale image whose pixels correspond directly to electrodes. The spatial resolution of the HD-MEA is often too low to resolve fine features of the microstructure like neighbouring microchannels. To overcome this challenge, the CAD design used to fabricate the microchannel architecture was aligned with the impedance map, enabling individual microchannels to be assigned to the recording electrodes. The impedance map was first normalised to [0,1] and converted to a binary mask by Otsu thresholding. Pixels corresponding to electrodes covered by the microstructure were grouped into valid connected components and small components under 0.05% of the image were discarded as thresholding noise. The map was then cropped to the bounding box enclosing all remaining connected components. The geometry of the microstructure was read from all layers of the CAD design, assuming no layer carried particular semantics, and was then scaled to the bounding box size while preserving its aspect ratio. The CAD design was aligned to the binary mask by maximizing their overlap, measured as the intersection over union (IoU). The CAD design was initially placed according to the aspect ratio of the impedance map, yielding two candidate alignments that differ by a 180° rotation. For each candidate, a grid search over translation parameters was performed, and the transformation with the highest overlap was retained. The alignment was then further refined by applying a similarity transformation whose parameters were optimized using Powell minimization of (1 - IoU).

### Geometry segmentation

The registered CAD design was rasterised at 6×6 supersampling and convolved with a normalised circular kernel whose radius was set to 0.8 times the chamber radius. Local maxima above a predefined threshold were identified as node centres. For each centre, the surrounding chamber body was isolated by morphological opening of the CAD mask, dilated by 0.4 times the chamber radius, and removed, yielding a mask containing only the microchannels.

### Selection

The microchannel mask was segmented into individual strands by per-pixel sub-pixel voting: each native pixel was assigned to the microchannel covering most of its 6 × 6 sub-pixels, with at least 10% of coverage, and tied pixels were left unassigned. The microchannels connecting two chambers formed a 4-microchannel bundle. Within each bundle, the electrodes were chosen using a joint combinatorial solver: for each of *k* target positions *z*_*i*_ along the shared bundle axis, one pixel per strand was selected to minimise

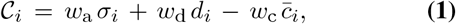

where *σ*_*i*_ is the spread of the chosen pixels’ projections onto the bundle axis (alignment), *d*_*i*_ is their mean projection’s distance to the target position *z*_*i*_, and 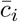 is their mean microchannel coverage. The weights *w*_a_, *w*_d_, *w*_c_ control the trade-off between alignment, position, and coverage. A minimum spacing constraint prevents electrodes from collapsing onto one pixel. Each microchannel received three electrodes, denoted `e0` and `e2` (the two ends, nearest the two connected chambers), and `e1` (mid-channel). The final number of electrodes routed per chip to perform the connectivity analysis was 924, within the routing budget of 1,020.

Due to the fact that the microstructure can sit rotated or tilted on the recording frame, bundles were indexed in the chip’s own reference frame rather than in image coordinates. The two principal axes of the chamber-centre cloud (via PCA) define the lattice’s column and row directions: each bundle, represented by its chamber-pair midpoint, was projected onto these axes and sorted row by row (bottom-left to top-right). Assigning indices in this intrinsic frame makes the labelling invariant to the chip’s placement and orientation. Microchannels within a bundle were ordered by their position perpendicular to the line joining the two chambers. The bundle identifier for every microchannel was stored in a single map carried in the geometry file, ensuring consistent chamber/microchannel/electrode labeling across all downstream stages. For each run, the electrode coordinates and geometry profile were stored for downstream processing. The per-chip processing time for the full three-stage procedure was less than 2 min.

The automated selection can be inspected and, where needed, refined through an interactive GUI before recording. The editor overlays the proposed electrodes on the chip binary map and supports modifying or rebuilding the electrode selection by moving, adding or deleting individual electrodes.

### Electrophysiological Recordings

Extracellular activity was recorded on a high-density CMOS microelectrode array (Maxwell MaxOne Plus PLM HD-MEA, MaxWell Biosystems) at a sampling rate of 20 kHz with an amplifier gain of 512 ×, using a fixed electrode selection covering the node-microchannels layout. Spontaneous activity was recorded over DIV15–DIV45 (31 consecutive days). Recording pipeline performance metrics are reported in **Fig. S21**. HD-MEA data were acquired in 15 min blocks and processed immediately after acquisition using a custom Python pipeline, with up to two recordings processed in parallel. Signals were high-pass filtered using a second-order Butterworth filter with a cutoff frequency of 200 Hz, followed by smoothing with a second-order Savitzky–Golay filter (60) using a 0.25 ms window. Spikes were detected using a smoothed nonlinear energy operator (SNEO) with lag *k* = 3. Within each 1.5 ms detection window, the largest peak exceeding 20 times the median absolute energy was classified as a spike. Detection parameters were selected based on previous work (61).

### Continuous long-term acquisition pipeline

Long-term recording was handled by a custom fault-tolerant pipeline that decoupled acquisition from spike detection. Two threads, whose recording and processing subprocesses were pinned to disjoint CPU cores, communicated through a queue: a recorder thread wrote each 15 min block to a local disk, while a processor thread detected spikes on completed blocks with the SNEO detector and parameters above (representative traces at active and inactive timepoints are shown in **Fig. S22**), storing the frame index, channel, electrode, and amplitude of each event; the queue absorbed transient backlogs so recording remained gap-free. Each per-block spike file was transferred to network storage with end-to-end integrity verification via a BLAKE3 checksum recomputed at the destination, with up to five re-transfers on mismatch. Additionally, the raw recording of a block was preserved alongside its spike waveforms every ∼ 8 h. Each block carried a timestamped filename from which the day *in vitro* was later derived (REC1 = DIV15, May 7 2026), and free-space guards, a watchdog, and a per-block log of durations, sizes, checksums, and any fallback conditions ensured continuity and a complete audit trail.

### Data Analysis

Activity, dynamical, connectivity and machine learning analyses were performed. To enable spatial analysis, recording electrodes were mapped to node indices leveraging the automated electrode selection geometry: for each microchannel, the endpoint electrodes adjacent to one node (`e0`) and the other node (`e2`) were retained and the mid-electrode (`e1`) was excluded. This made every analysed electrode attributable to a specific node. Each node was assigned a single canonical index (1 to 33, sorted from bottom-left to top-right position) used consistently across all analyses, so that a given index denotes the same physical node in every feature modality.

Each node was further assigned two complementary spatial coordinates on the lattice. The first is absolute *layer*, corresponding to the six physical rows of nodes (1-6, layer 1 = lowest physical row). The second, defined by the disease-model network, is a topological *hop distance*: the minimum number of node-to-node steps from each node to the progerin-expressing disease core node (1-6, hop 1 = closest nodes to the core). The core is the origin (*hop 0*), and the surrounding nodes form successive shells at increasing hop distance. To enable matched per-hop comparisons in the absence of a disease core, the control network was assigned the same homologous *hop 0* at the same lattice position, making both conditions directly comparable. The disease core node was excluded from layer and hop distance statistics in both modalities.

### Activity features

For every 15 min recording block, six per-node activity features were computed. Unless stated otherwise, features were computed per electrode and then averaged across the node’s electrodes. Electrodes with fewer than 5 spikes in a block were excluded from the corresponding feature.

- **Firing rate** (Hz): per-electrode spike count divided by block duration, averaged across the node’s electrodes.
- **ISI irregularity** (CV_ISI_): the coefficient of variation of the inter-spike intervals, 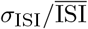, capturing spike-train irregularity. Values near 1 indicate Poisson-like firing, below 1 more regular, and above 1 more bursty firing.
- **Burst duration, burst frequency** and **burst amplitude**: bursts were detected with a MaxInterval algorithm (intra-burst ISI threshold 100 ms, merge gap 50 ms, minimum 3 spikes, minimum duration 10 ms). Burst duration is the mean burst length; burst frequency is the number of bursts per second; burst amplitude is the mean across bursts of the peak spike count within a sliding 10 ms window, expressed in Hz.
- **Spike synchrony**: mean pairwise spike-time tiling coefficient (STTC) (62) over all pairs of the node’s active electrodes (Δ*t* = 10 ms). For spike trains *A, B*:

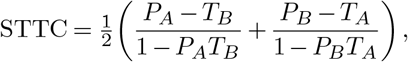

where *T*_*X*_ is the fraction of recording time within ± Δ*t* of a spike of train *X* and *P*_*X*_ the fraction of *X*’s spikes within ±Δ*t* of a spike of the other train.

Per-node values were retained at their native 15 min block resolution and summarised as the median per hour and per DIV, limiting the influence of transient block-level fluctuations. For each electrophysiological feature, the trajectory of each layer was calculated as the mean ± 95% confidence interval across all nodes within that layer.

### Dynamical features

On the same per-block basis, descriptors of each node’s activity were computed. Each node’s spikes were binned at 100 ms to form an activity time series, from which the following descriptors were computed: (i) the effective activity timescale (EAF), the first lag at which the series’ autocorrelation drops below 1*/e*. Lags up to 100 were considered and the metric is reported in seconds; (ii) the automutual information (AMI) between the series and its copy shifted by the EAF lag, estimated with a direct (plug-in) estimator on an 8 × 8 joint histogram, in bits. Each 15 min block provides *N* ≈ 9000 samples (100 ms bins) against *R* = 64 joint response classes (*N/R* ≈ 140, well above the *N/R* ≳ 32 guideline (63)), so residual sampling bias is negligible; (iii) the transition entropy, the Shannon entropy (in bits) of the distribution of consecutive state transitions of a Markov chain built over five discrete activity levels obtained by rank-binning the activity into quintiles (lowest to highest 20%), with higher values indicating more random, less predictable state switching; and (iv) the skewness of the series’ first differences (TRS), 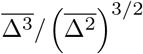, where Δ denotes the change in activity between consecutive 100 ms bins. Separately, a population activity series was formed for the whole network and for each layer and hop group by summing spike counts across all electrodes in 100 ms bins. Each series was modelled with a three-state Poisson hidden Markov model (HMM) (64–66). A three-state model was chosen for interpretability, giving discrete low-, intermediate- and high-activity regimes of the population. Each state emitted spike counts under a Poisson distribution with one rate parameter per state. The model was fitted with the Baum-Welch algorithm using the implementation’s default initialization, a convergence tolerance of 10−3, a maximum of 100 iterations, and a fixed random seed for reproducibility. The most likely state sequence was decoded with the Viterbi algorithm (67), states were ordered by ascending mean activity, and dwell time was defined as the duration of each continuous run in a state (number of consecutive 100 ms bins multiplied by the bin width). Runs truncated by recording boundaries were excluded from dwell-time statistics to avoid downward bias, while visit counts included all runs. For each state, the mean and standard deviation of dwell times, number of visits, and total and fractional time occupied are reported. Nodes or groups below the minimum spike or length requirement for a given metric were set to NaN.

### Transfer-entropy functional connectivity

Functional connectivity was estimated by transfer entropy (TE) (68), a model-free measure of directed, nonlinear and lagged statistical dependency between spike trains, quantifying how much the activity of one node reduces uncertainty about the future activity of another node. For a source process *X* and target process *Y*, the transfer entropy from *X* to *Y* is given as:

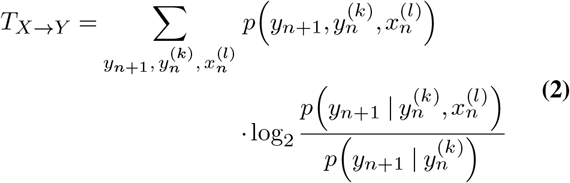

where 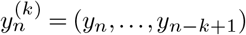 and 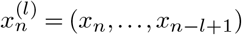 are the target and source history vectors of lengths *k* and *l*, and the sum runs over all (discrete) states of 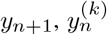 and 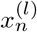. TE thus quantifies the reduction in uncertainty about the target’s next state contributed by the source’s past, beyond what the target’s own past already explains.

The functional connectivity of each network was represented as a directed, weighted graph whose nodes are chambers of the lattice layout and whose directed edge *i* → *j* carries the total significant TE flowing from node *i* to node *j*. The tested signal pairs are therefore the `e0`–`e2` endpoints of every microchannel, evaluated in both directions.

To optimise computational cost, TE was computed on a single 15 min recording per day rather than all blocks. Each day’s recording was selected from an afternoon–evening window, predominantly 16:00–22:00, and a single one in the morning right before A*β* addition. This yields one directed TE network per DIV per condition (DIV 15–45; 31 graphs). Computational performance metrics for the TE pipeline are reported in **Fig. S23**. Spike trains were binarised at a 0.25 ms bin width Δ*t* (5 samples at 20 kHz; Eq. 4). For each microchannel, bivariate pairwise TE was estimated between its source (`e0`) and target (`e2`) electrodes in both directions using the discrete estimator of the Java Information Dynamics Toolkit (JIDT) (69) in bits (base 2).

Transfer entropy was estimated using a source–target delay *u* swept over 1–8 bins (0.25 ms to 2.0 ms), keeping the delay of maximal TE per ordered pair, a target history of *k* = 4 bins (1.0 ms) and a single-bin source embedding (*l* = 1), both with unit embedding delays (*k*_*τ*_ = *l*_*τ*_ = 1). The delay parameter generalises Eq. 2 by evaluating transfer entropy over a range of source–target delays, such that the source history is taken *u* bins before the target history 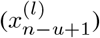. For significance, 100 permutations of the source train were performed at each delay, and the *p* value of the delay of maximal TE was retained and corrected across all ordered pairs by the Benjamini–Hochberg false-discovery-rate (FDR) method at *q* ≤ 0.05 (70). Only pairs with ≥ 5 spikes on both electrodes were tested.

The bin width Δ*t*, the delay window *u*, and the target history *k* were grounded in the physics of axonal propagation, published conduction velocities for cortical axons, and the cultures’ own measured propagation delays (Eqs. 3–6). A spike recorded at the source electrode `e0` reappears at the target `e2` after a conduction delay *τ*, set by the endpoint separation *d* and the conduction velocity *v*. TE recovers this delay as the lag *u* of maximal information transfer (Eq. 3). The bin width (Eq. 4) is the finest resolution at which the binary trains remain adequately populated. The 0.25 ms to 2.0 ms window is fixed by the geometry and the measured speed. The `e0`–`e2` separations span 99 µm to 350 µm (median 149 µm control, 137 µm progerin), and the conduction velocity, measured independently of TE by cross-correlating the two electrodes’ binned spike trains and taking the peak lag, is *v* ≈ 0.30 m*/*s (0.32 ± 0.03 control, 0.35 ± 0.02 m*/*s progerin; mean ± SD across days), consistent with published values for unmyelinated cortical axons on HD-MEAs (≈ 0.2–0.8 m*/*s (71)). The implied conduction delays therefore span 0.33 ms to 1.17 ms (Eq. 5), all inside the sweep. The target history *k* = 4 spans 1.0 ms (Eq. 6), which is on the order of the neuronal absolute refractory period, and ensures that the estimated TE reflects information contributed by the source. This parameter choice enabled the TE estimation between all electrode pairs for a 15 min recording to be computationally tractable.

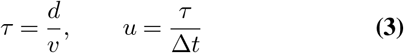

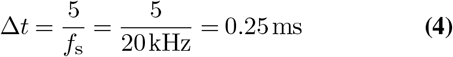

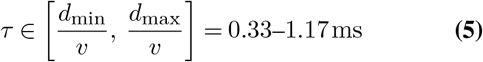

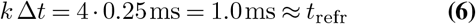

### Graph-theoretic network analysis

For each daily recording, a directed, weighted graph was constructed on the 33-chamber lattice. A directed edge *i* → *j* was included when the transfer entropy *T*_*ij*_ was significant after Benjamini– Hochberg false-discovery-rate correction (*α* = 0.05) (70), and was weighted by *T*_*ij*_ in bits. Because each pair of chambers is bridged by a bundle of parallel microchannels, the transfer entropies of all significant microchannels linking the same ordered pair were summed, so that an edge weight represents the total significant directed information flow between the two chambers. Edge orientation was assigned according to the physical microchannel geometry. Within-layer bundles were represented by bidirectional edges, whereas across-layer bundles retained only the dominant direction of information flow inferred from transfer entropy analysis, which was directed from lower to higher network layers in 98.5% of bundles in both network types.

Every measure was either computed in a directed form on the graph *G* or in an undirected form on a symmetrized graph *G*^*u*^, in which reciprocal edges were combined by summing their weights 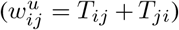. For across-layer edges, retained in a single direction, the contribution is that direction’s weight alone. For path-based measures, information-flow weights were converted to distances by inversion: weighted shortest-path length and global efficiency used *d*_*ij*_ = ⟨*T* ⟩*/T*_*ij*_, the inverse transfer entropy scaled by the recording’s mean edge weight ⟨*T*⟩, so that a typical edge has unit length and path-based quantities are comparable across recordings with differing overall transfer-entropy magnitude. Local efficiency used the same inversion, with ⟨*T*⟩ taken over the edges of each node’s neighbourhood subgraph and betweenness centrality used the unscaled inverse *d*_*ij*_ = 1*/T*_*ij*_.

Graph measures were computed with NetworkX (72). Per-node features comprised total degree and strength, total degree centrality (in+out degree normalised by *n* − 1), betweenness centrality, PageRank, the weighted clustering coefficient computed using the Onnela form (73) on the undirected graph *G*^*u*^, local efficiency with the harmonic mean of the inverse shortest-path lengths within each node’s neighbourhood subgraph, a transfer-entropy asymmetry index (out-strength minus in-strength), a hub score 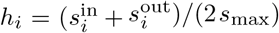, where 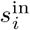 and 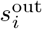 are node *I* ‘s in- and out-strength and *s*_max_ is the largest single-direction strength in the network, and the participation coefficient with respect to the community partition. Network-level features comprised density, degree assortativity (for *G*, the out–in assortativity: the Pearson correlation, over all directed edges *i* → *j*, between the out-degree of source node *i* and the in-degree of target node *j*), global efficiency, characteristic path length (mean weighted shortest-path length), diameter, modularity, and small-worldness *σ*.

Three graph metrics were reported longitudinally. Degree centrality was taken as the weighted total strength of each node as the sum of its incoming and outgoing significant transfer entropy on the directed graph, normalised within each recording by the maximum node strength so that trajectories are comparable across days. Global efficiency was computed on the undirected graph as the mean of the inverse shortest-path length over all *n*(*n* − 1) ordered pairs (74), with edge weights converted to distances by inverting the transfer entropy. Pairs lying in different connected components contribute 1*/*∞ = 0, so the measure remains defined when some chambers carry no significant edges. Modularity was reported undirected with the Louvain modularity (75) on the symmetrised graph.

Community structure was estimated using the Louvain method (75) (resolution 1.0). Small-worldness was quantified as *σ* = (*C/C*_rand_)*/*(*L/L*_rand_) (76), where *C* and *L* are the mean clustering coefficient and characteristic path length of the unweighted topology, relative to 100 degree-preserving random graphs generated by double-edge swap for *G*^*u*^ and by the directed configuration model for *G*; *L* was evaluated on the largest (strongly) connected component. Communities were computed with `python-louvain` 0.16.

### Machine learning

Activity, dynamical and graph/TE features were merged on (condition, DIV, node) at the canonical node index, resulting in a dataset 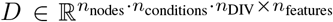 with *n*nodes = 33, *n*conditions = 2, *n*_DIV_ = 31 and *n*_features_ = 18 (six activity, four dynamical and eight graph/TE features), containing 2046 complete samples. Features were *z*-score normalised before any embedding or model fitting. For the cross-validated classifiers, this normalisation was fit on the training partition of each fold only, to prevent information leakage. Low-dimensional structure was visualised with *t*-distributed stochastic neighbour embedding (*t*-SNE) (77) (two components, PCA initialisation, automatic learning rate and perplexity 30, reduced for small subsets), computed separately per DIV (at six representative timepoints: DIV 15, 20, 24, 30, 35, and 40), per absolute layer and per hop distance from the disease core.

Condition separability over time was quantified by training a random-forest classifier (78, 79) (300 trees, balanced class weights, disease-model nodes as the positive class) to discriminate control from disease-model nodes *within each DIV* under stratified 5-fold cross-validation, reporting accuracy and precision. Feature contributions were quantified with SHAP values (80) from a random-forest classifier fit to each phase; the mean absolute SHAP value per feature was computed separately for the pre-A*β* (DIV*<* 23) and post-A*β* (DIV ≥ 23) phases. Machine-learning analyses used scikit-learn 1.9.0 and SHAP 0.52.0.

### Spatial dependency analysis

To investigate spatial dependence in the merged features for each DIV, a partial Mantel test was used. Each feature was first z-score normalised, and the dimensionality of the feature vector was reduced by retaining the first eight principal components. Partial Mantel correlations were then computed separately for hop distance and layer distance from the diseased core:

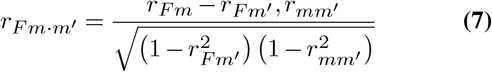

where *m* ∈ {layer, hop} denotes the distance metric of interest, *m*^′^ ∈ {layer, hop} \ *m* denotes the distance metric being controlled for, *F* is the PCA-reduced feature representation, and *r* is the Pearson correlation between the corresponding distance matrices. Statistical significance was assessed using 10,000 node-label permutations.

## DATA AVAILABILITY

The data supporting the findings of this study and the code used for evaluation will be publicly available upon acceptance.

## AUTHOR CONTRIBUTIONS

The author contributions follow the [CRediT](https://credit.niso.org/) system. **J.K**. and **A.B**.: Data curation, Formal analysis, Investigation, Methodology, Software, Visualization, Writing – original draft. **G.A**.: Data curation, Formal analysis, Investigation, Software, Visualization. **L.J**. and **B.M**.: Methodology, Resources. **E.D**. and **Y.R**.: Data curation, Formal analysis, Investigation, Visualization. **V.V**.: Methodology, Software. **M.P**. and **K.C**.: Funding acquisition, Resources. **N.W.H**.: Conceptualization, Methodology, Data curation, Formal analysis, Funding acquisition, Investigation, Project administration, Supervision, Visualization, Writing – original draft. All authors contributed to Writing – review & editing.

## FUNDING

This research was supported by ETH Zürich and the Swiss National Science Foundation (SNSF) [project number 182779].

### ACKNOWLEDGEMENTS

The authors thank Prof. János Vörös for access to laboratory facilities and support throughout this work. We are grateful to Prof. Lennart Niels Cornelisse and Juraj J. Ondris at Amsterdam UMC for providing inhibitory neurons, and to Prof. Catherine Verfaillie and Dr. Karan Ahuja at KU Leuven for providing astrocytes and for their guidance on astrocyte differentiation. We thank Novartis for providing NGN2-derived excitatory neurons.

## COMPETING FINANCIAL INTERESTS

The authors declare that the research was conducted in the absence of any commercial or financial relationships that could be construed as a potential conflict of interest.

## DECLARATION OF GENERATIVE AI AND AI-ASSISTED TECHNOLOGIES IN THE WRITING PROCESS

During the preparation of this work the authors used *Claude* Sonnet 4 (Anthropic, San Francisco, CA, USA) in order to improve language and readability. After using this tool, the authors reviewed and edited the content as needed and take full responsibility for the content of the publication.

## Supplementary Materials

**Figure S1.**
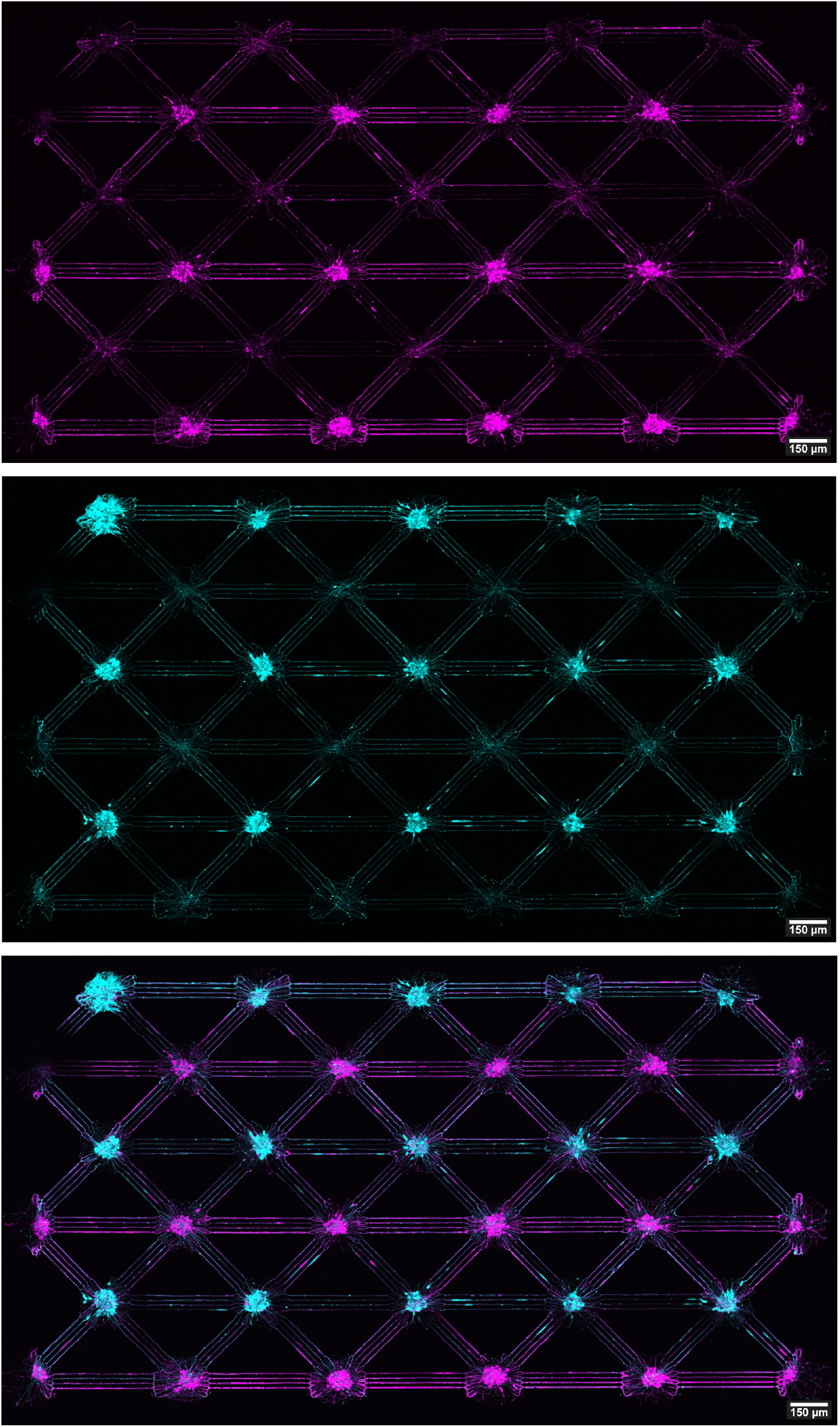
Bidirectional 33-node network used for quantification of axon directionality and neurite width. Fluorescence overview of a 33-node network with bidirectional straight channels both within and between layers, used as the control for quantification of maximum axon fluorescence intensity and neurite width in **Figure 1D–F** of the main paper. **(A)** GFP-expressing neurons (magenta). **(B)** RFP-expressing neurons (cyan). **(C)** Merged image showing both populations.

**Figure S2.**
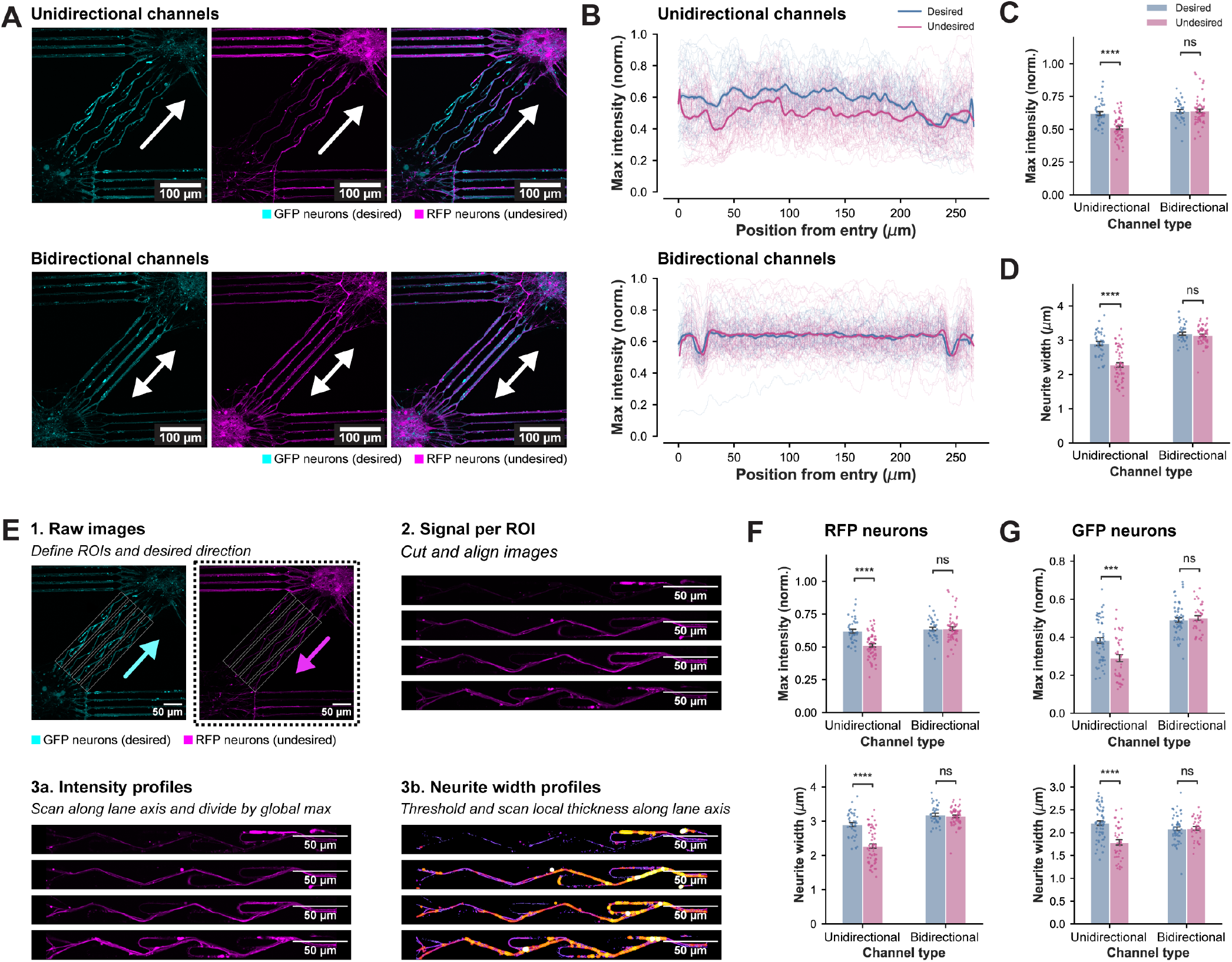
Directional axonal connectivity is maintained through DIV 35, and image analysis pipeline for quantification of axon directionality and neurite width, supplementary to Figure 1. **(A)** Representative fluorescence images of unidirectional (top) and bidirectional (bottom) channels at DIV 35, showing GFP neurons (cyan, desired direction) and RFP neurons (magenta, undesired direction). White arrows indicate the preferred direction of axonal outgrowth. **(B)** Full intensity profiles along the channel axis (0– 270 µm from channel entry) for desired (blue) and undesired (pink) directions in unidirectional (top) and bidirectional (bottom) channels at DIV 35, showing individual profiles (thin lines) and mean (thick lines). **(C)** Unidirectional channels exhibited significantly higher axon growth in the preferred versus non-preferred direction at DIV 35, while bidirectional channels showed no significant directional bias. \*\*\*\**p <* 0.0001; ns, not significant. **(D)** Axon bundles travelling in the undesired direction were significantly narrower than those in the desired direction in unidirectional channels at DIV 35. No significant difference in neurite width was observed in bidirectional channels. \*\*\*\**p <* 0.0001; ns, not significant. **(E)** Overview of the image analysis pipeline. Step 1: raw fluorescence images with regions of interest (ROIs) drawn around individual channels, with the desired direction defined per channel (cyan arrow, GFP neurons; magenta arrow, RFP neurons). Step 2: individual lanes are cropped and aligned along the channel axis. Step 3a: intensity profiles are extracted by scanning along the lane axis and recording the maximum pixel value across the lane width at each position, normalised to the global per-image, per-channel maximum. Step 3b: neurite width profiles are extracted by applying CLAHE contrast enhancement, Otsu thresholding, and the Local Thickness plugin to assign each foreground pixel the diameter of the largest inscribed circle, with the maximum local thickness sampled at each axial position. **(F)** Quantification of maximum fluorescence intensity (normalised, top) and neurite width (bottom) in desired versus undesired directions for RFP neurons in unidirectional and bidirectional channels. \*\*\*\**p <* 0.0001; ns, not significant. **(G)** Corresponding quantification for GFP neurons. \*\*\**p <* 0.001; \*\*\*\**p <* 0.0001; ns, not significant.

**Figure S3.**
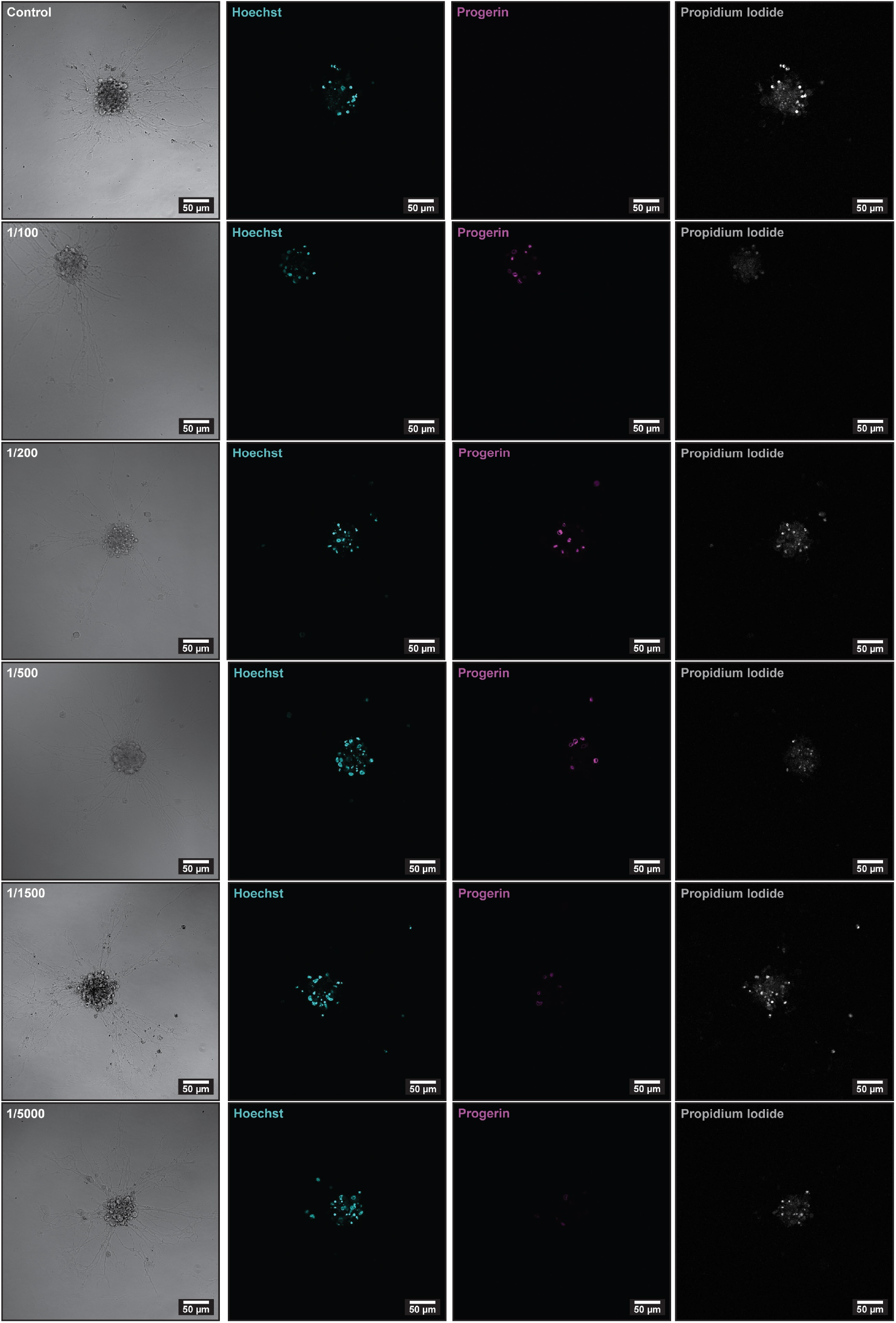
Representative images of progerin expression across lentiviral dilutions. Representative images of spheroids for each condition tested, from control to 1:5000 lentiviral dilution (rows, top to bottom). Columns show transmitted light (TD), Hoechst nuclear staining (cyan, used for quantification of total cell number per spheroid), GFP-progerin expression (magenta), and propidium iodide staining (gray, cell viability). A dose-dependent reduction in GFP-progerin-positive cells is visible with increasing dilution, consistent with the quantification shown in **Figure 2D** of the main manuscript.

**Figure S4.**
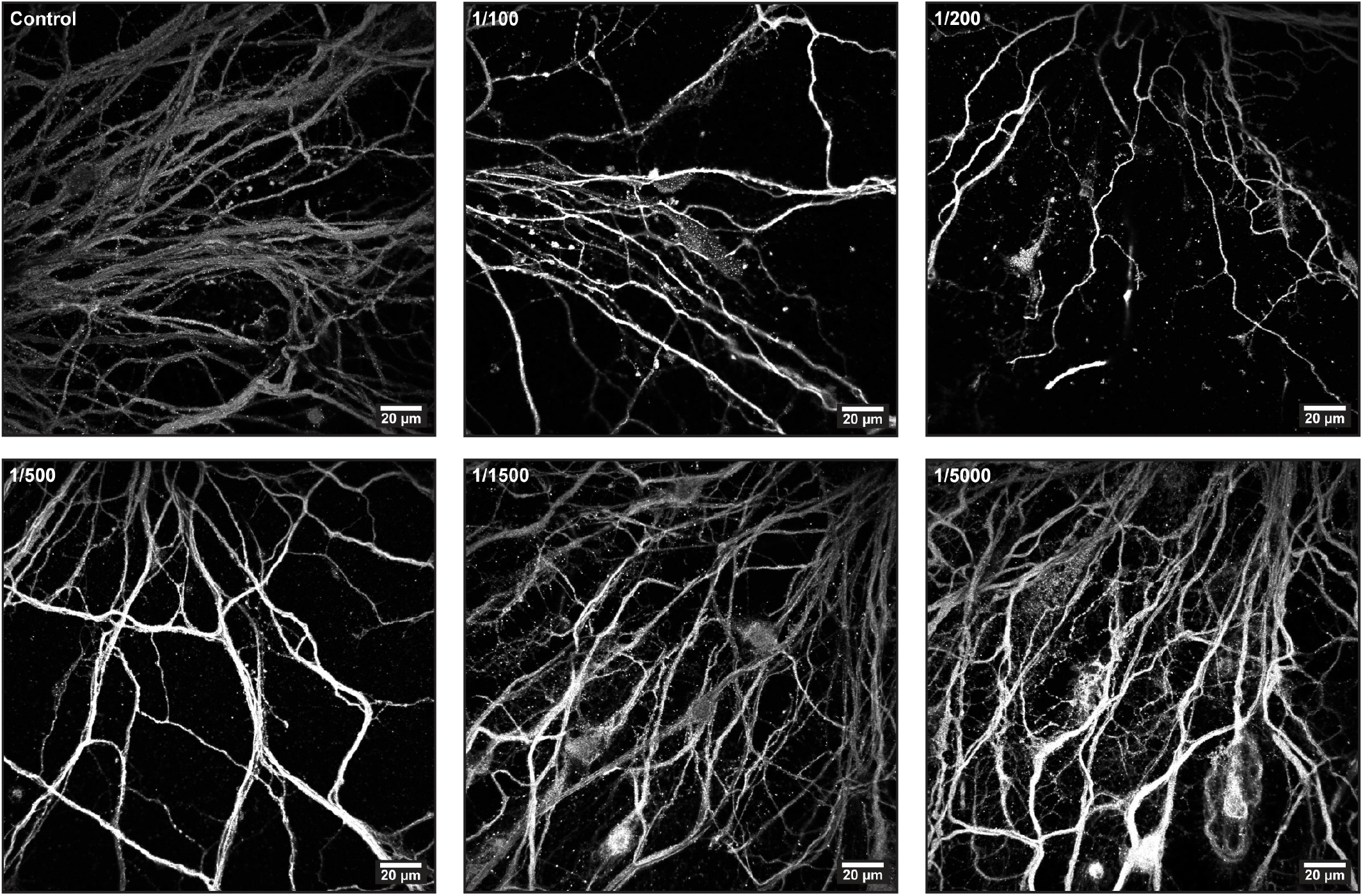
Representative images of neurite morphology across lentiviral dilutions. Representative confocal images of *β*3-tubulin-stained neurites in the spheroid halo for each condition tested, from control to 1:5000 lentiviral dilution. Images were used for quantification of neurite density shown in **Figure 2F** of the main manuscript.

**Figure S5.**
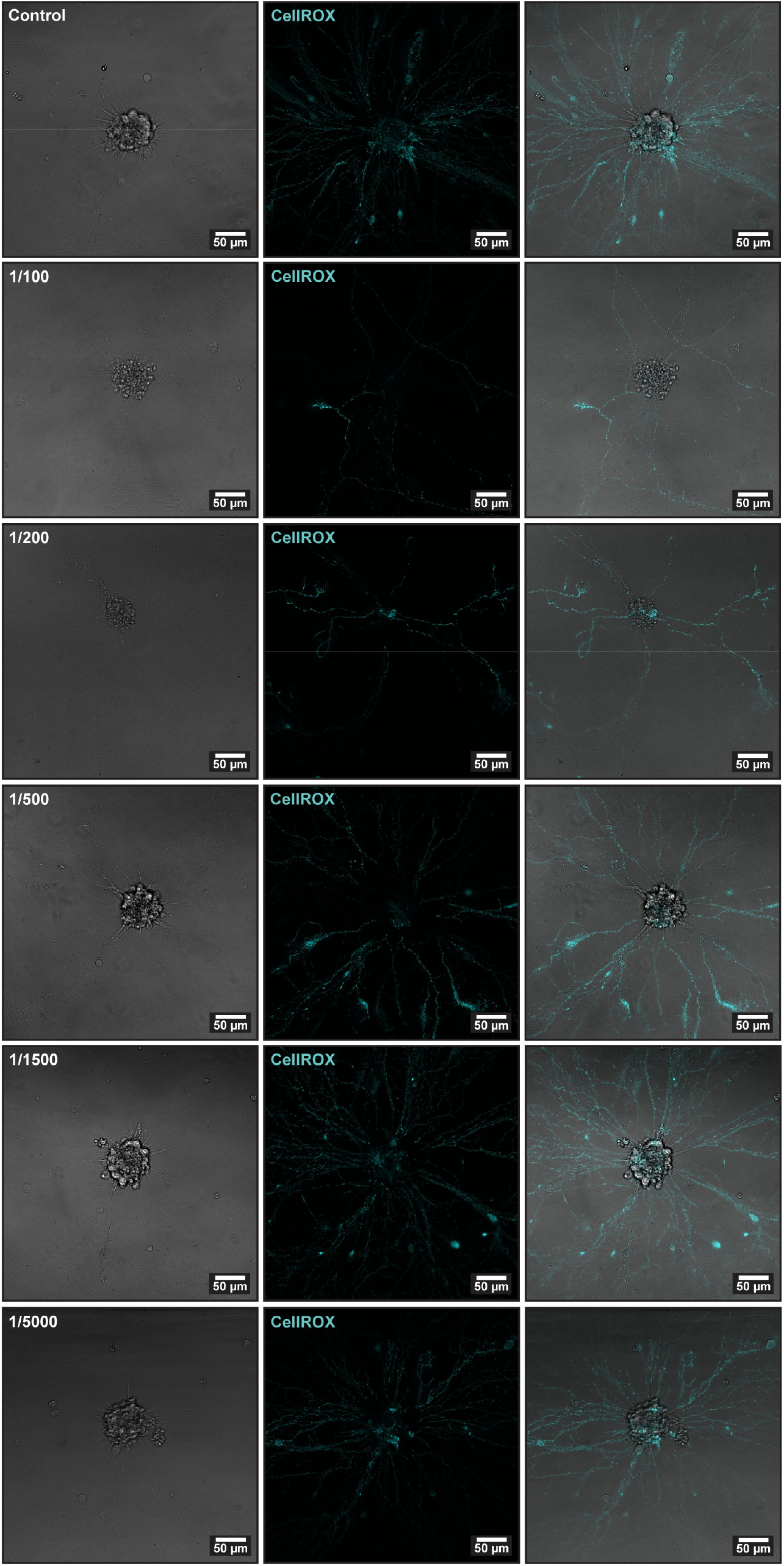
Representative CellROX images across lentiviral dilutions. Representative live images of spheroids stained with CellROX Deep Red for each condition tested, from control to 1:5000 lentiviral dilution (rows, top to bottom). Columns show transmitted light (TD), CellROX fluorescence (cyan), and merged images, respectively. CellROX signal reflects intracellular oxidative stress and was used for quantification of normalised integrated density shown in **Figure 2G** of the main manuscript.

**Figure S6.**
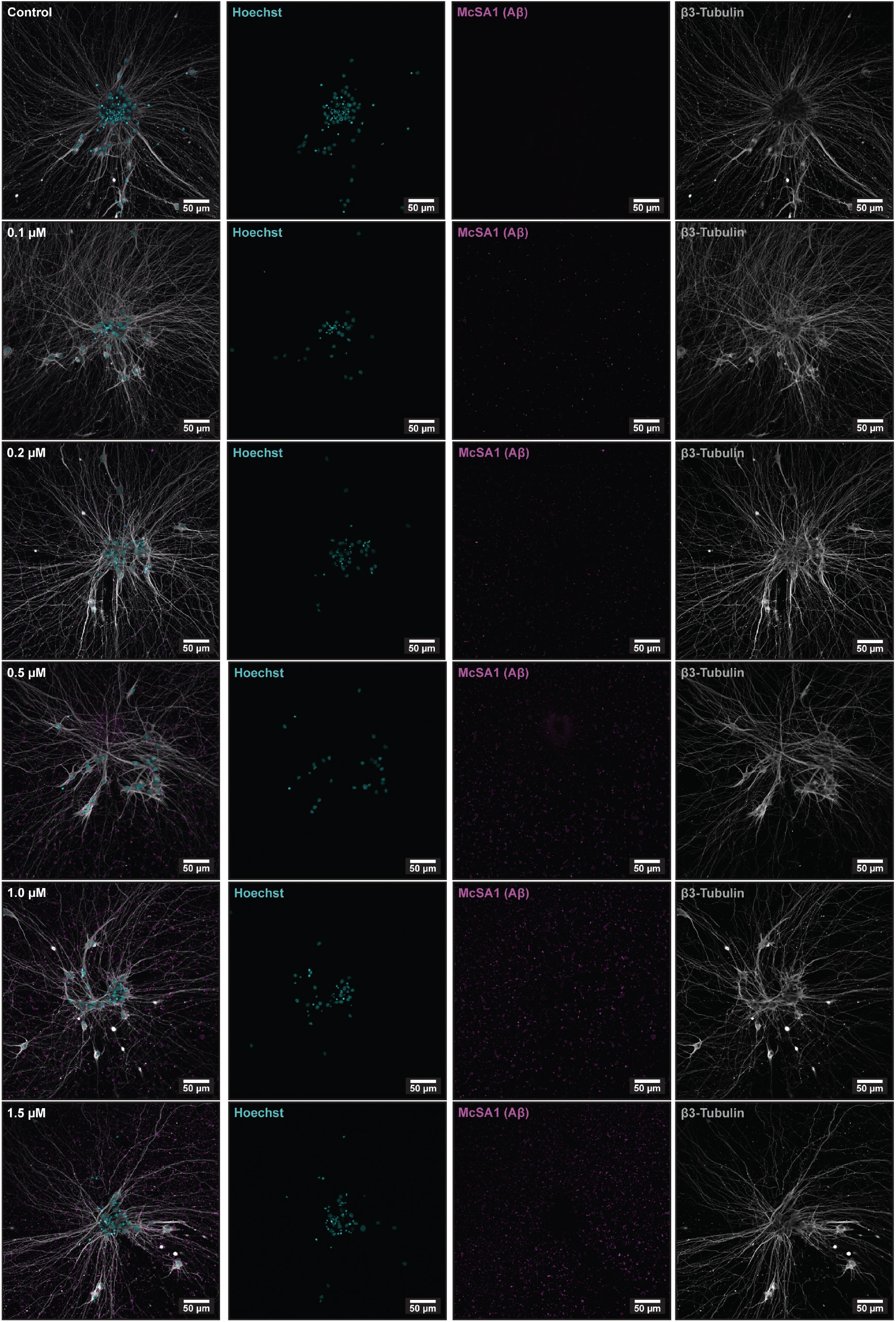
Representative immunofluorescence images of A*β* aggregate formation across concentrations at 20x magnification. Representative confocal images of spheroids fixed and stained 18 days after A*β* addition at DIV 23, for control and A*β* concentrations of 0.1, 0.2, 0.5, 1.0, and 1.5 µM. A dose-dependent increase in perineuronal A*β* aggregate formation is visible, consistent with the quantification shown in **Figure 2I–K** of the main manuscript.

**Figure S7.**
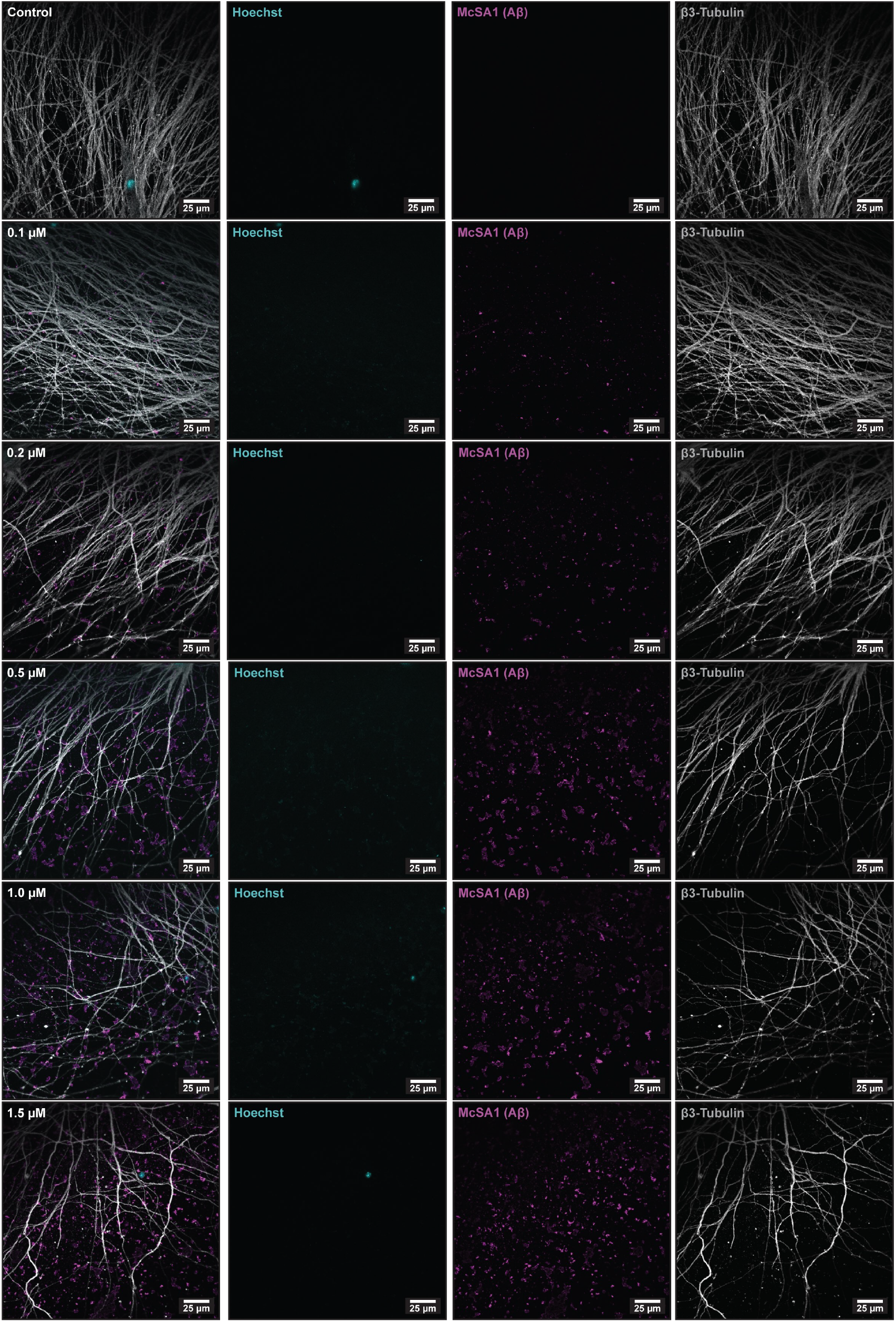
Representative immunofluorescence images of A*β* aggregate formation across concentrations at 60x magnification. Representative confocal images of spheroids fixed and stained 18 days after A*β* addition at DIV 23, for control and A*β* concentrations of 0.1, 0.2, 0.5, 1.0, and 1.5 µM. A dose-dependent increase in perineuronal A*β* aggregate formation is visible, consistent with the quantification shown in **Figure 2I–K** of the main manuscript.

**Figure S8.**
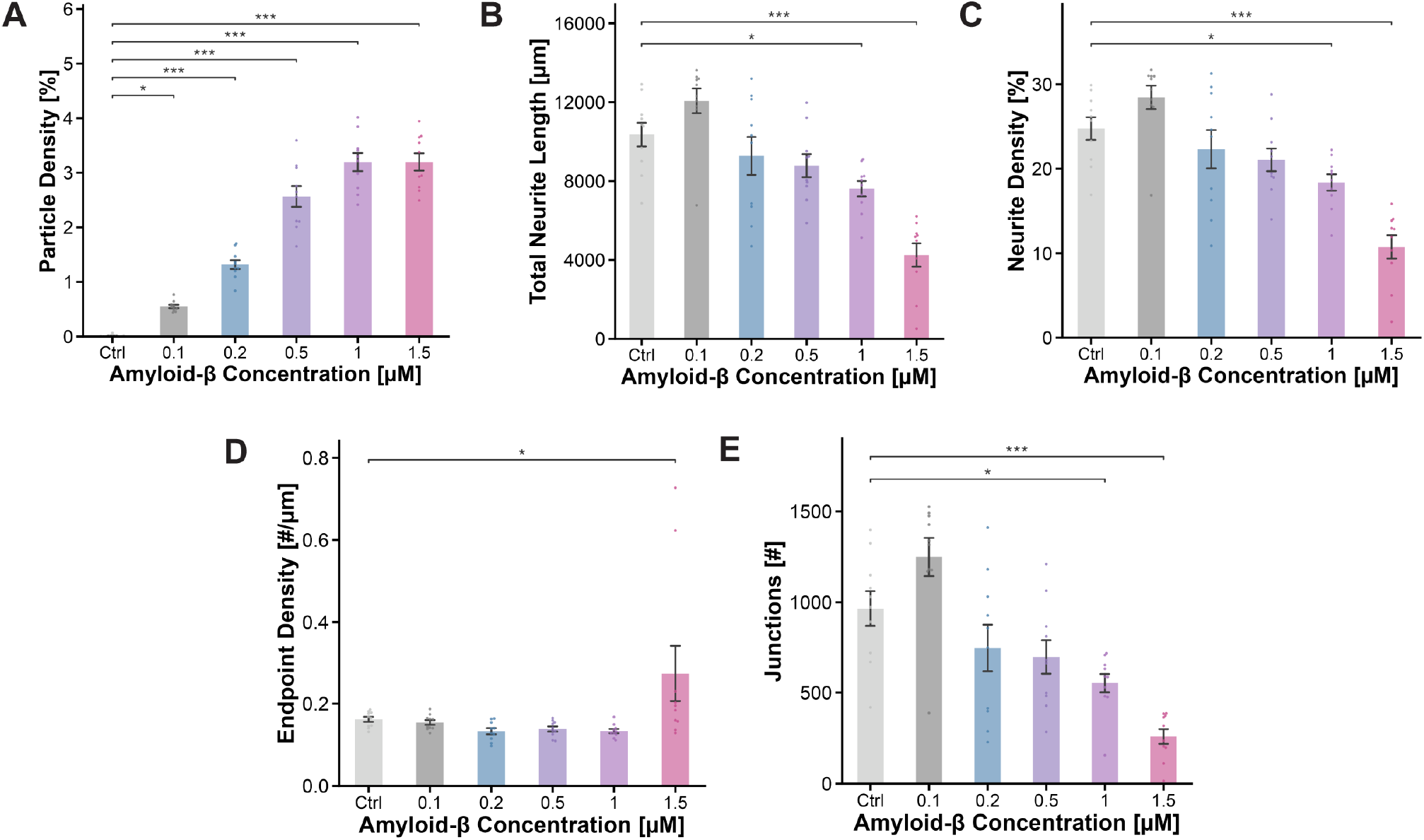
Extended morphological characterisation of amyloid-*β* concentration effects on neuronal networks, supplementary to Figure 2. **(A)** A*β* aggregate particle density increased significantly with concentration, with significant differences observed between the control and 0.1, 0.2, 0.5, 1, and 1.5 µM conditions. **(B)** Total neurite length declined with increasing A*β* concentration, with significant reductions at 1 and 1.5 µM compared to control. **(C)** Neurite density showed a similar concentration-dependent decline, with significant reductions at 1 and 1.5 µM compared to control. **(D)** Endpoint density showed no significant change across concentrations until 1.5 µM, where a significant increase was observed compared to control, consistent with neurite fragmentation at higher concentrations. **(E)** The number of neurite junctions declined significantly at 1 and 1.5 µM compared to control, indicating progressive disruption of neurite network complexity. Only statistical comparisons to the control condition are shown. Full pairwise statistics are reported in Supplementary Tables 17–26. \*\*\**p <* 0.001; \*\**p <* 0.01; \**p <* 0.05; ns, not significant.

**Figure S9.**
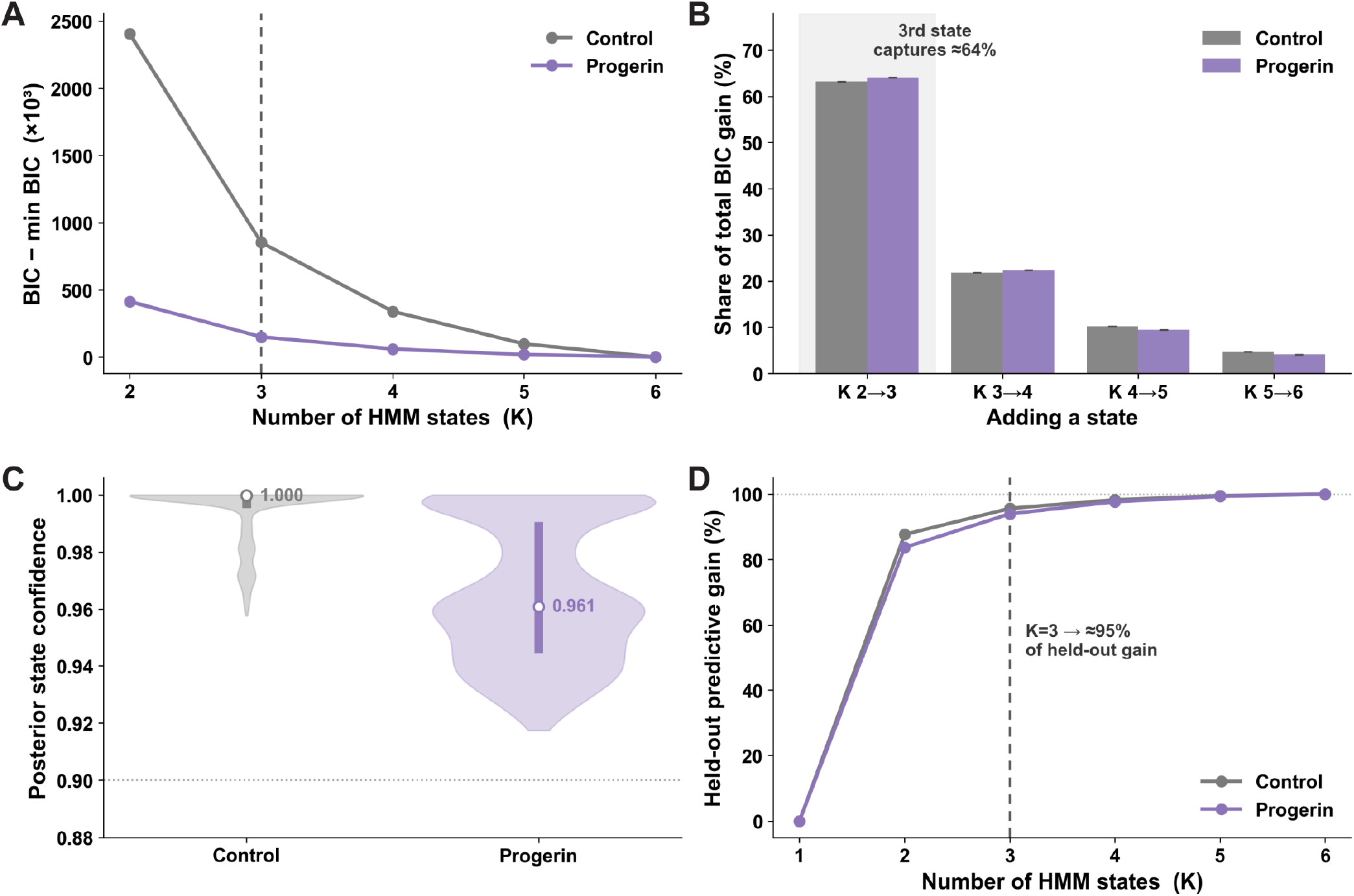
Poisson-HMM model selection and validation, supplementary to Figure 3. **(A)** BIC relative to the minimum BIC as a function of the number of hidden states (*K*) for control and progerin/A*β* networks. BIC decreases monotonically with *K*, with the largest gain observed at *K* = 3 (dashed line). **(B)** Share of total BIC gain contributed by each additional state. The transition from *K* = 2 to *K* = 3 accounts for approximately 64% of the total gain in both conditions, with strongly diminishing returns thereafter. **(C)** Posterior state assignment confidence for *K* = 3. Control networks show near-perfect assignment confidence (median 1.000), while progerin/A*β* networks show a median confidence of 0.961, both well above the 0.90 threshold (dotted line). **(D)** Held-out predictive gain as a function of *K*. At *K* = 3, approximately 95% of the maximum held-out predictive gain is already captured in both conditions, supporting *K* = 3 as the optimal and generalisable choice. A 3-state Poisson-HMM was fit to 100-ms population activity bins across 5477 hourly recordings (2818 control, 2659 progerin). The expectation-maximisation algorithm converged in 100% of fits.

**Figure S10.**
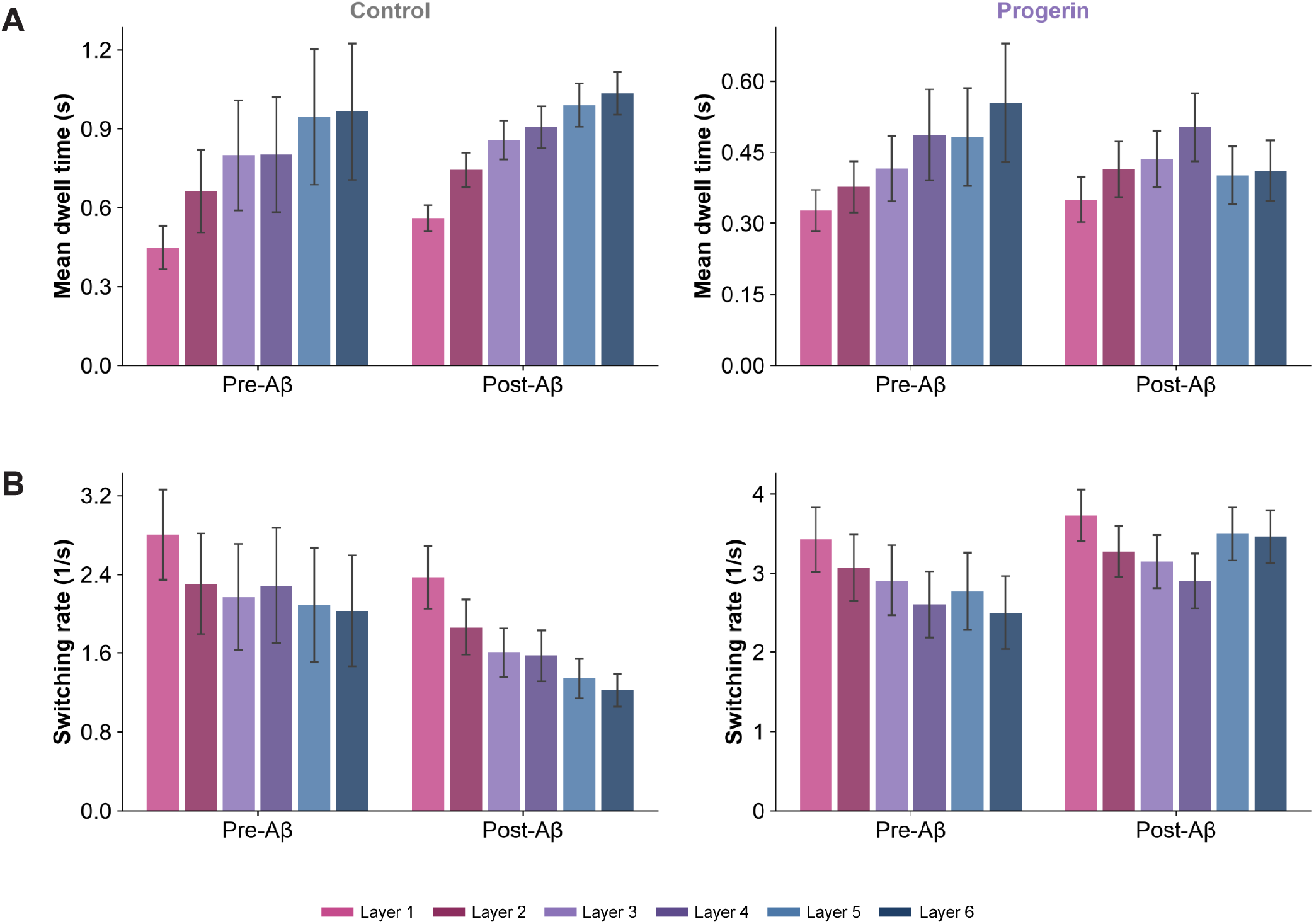
Layer-resolved dwell time and switching rate in control and progerin/A*β* networks, supplementary to Figure 3. **(A)** Mean dwell time by layer, before and after A*β* addition, for control (left) and progerin/A*β* (right) networks. In control networks, dwell time increased with layer both pre- and post-A*β*, with an overall increase following A*β* addition across all layers. The progerin/A*β* network showed a similar layer-dependent trend but with lower overall dwell times and a less pronounced increase following A*β* addition. **(B)** Switching rate by layer, before and after A*β* addition, for control (left) and progerin/A*β* (right) networks. Control networks showed a layer-dependent decrease in switching rate that became more pronounced following A*β* addition. In contrast, the progerin/A*β* network showed weaker layer dependence and an increase in switching rate across all layers following A*β* addition. Bars represent mean ± SEM across nodes within each layer.

**Figure S11.**
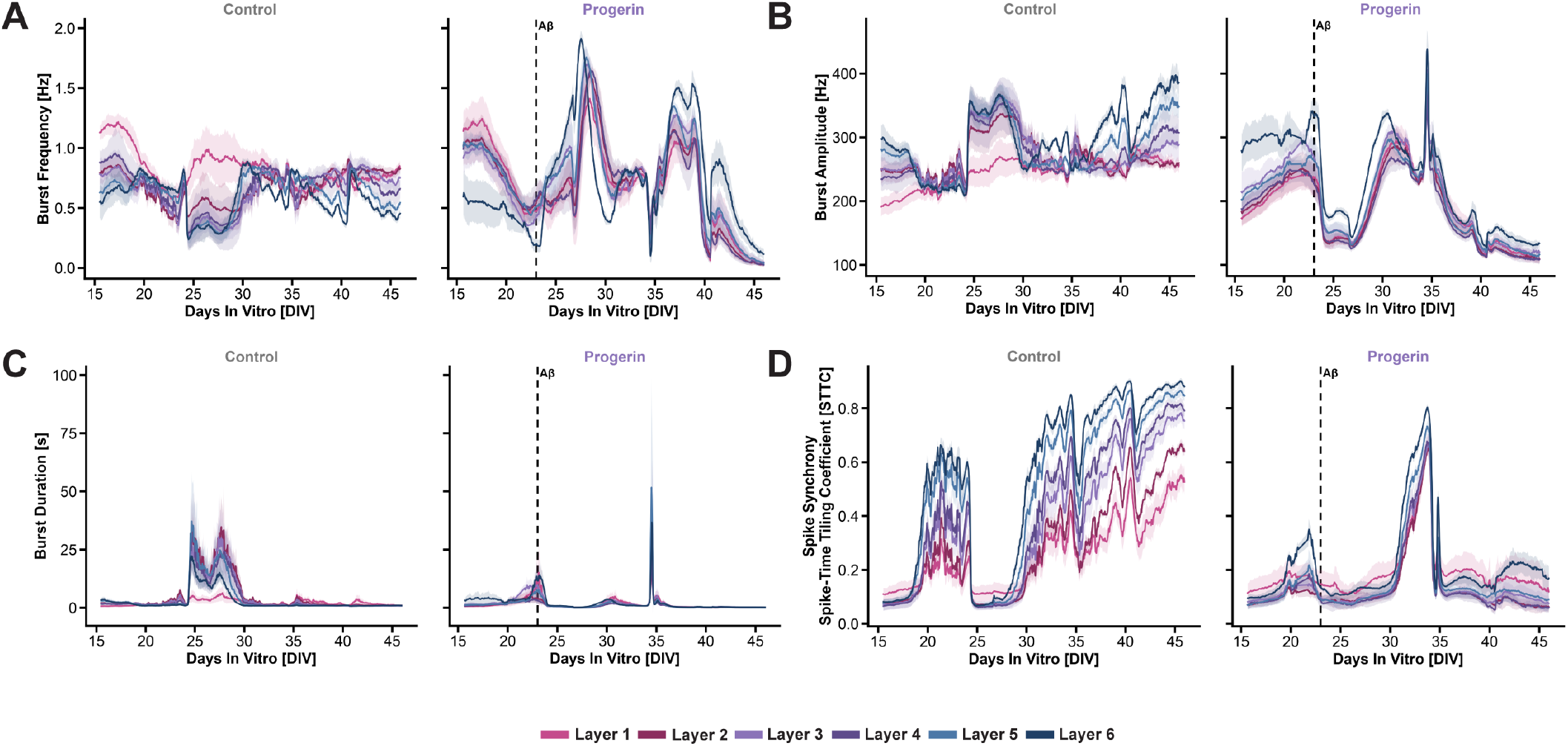
Longitudinal burst dynamics and spike synchrony across layers in control and progerin/A*β* networks, supplementary to Figure 3. **(A)** Burst frequency over time for control (left) and progerin/A*β* (right) networks, coloured by layer. A transient increase in burst frequency was observed immediately following A*β* addition at DIV 23 in the progerin/A*β* network, particularly in higher layers, before declining progressively. **(B)** Burst amplitude over time. Control networks displayed a gradual layer-dependent increase throughout the recording period, while the progerin/A*β* network showed a marked decline in burst amplitude following A*β* addition, with loss of the layer-dependent stratification observed in controls. **(C)** Burst duration over time. A pronounced transient increase was observed in control networks around DIV 24–26, coinciding with the media change, but was largely absent in the progerin/A*β* network. Spike synchrony, quantified as the Spike-Time Tiling Coefficient (STTC), over time. Control networks showed a progressive layer-dependent increase in synchrony throughout the recording period, while the progerin/A*β* network displayed markedly reduced and flatter synchrony following A*β* addition. Lines represent mean ± SEM across nodes within each layer. Dashed vertical line indicates timing of A*β* addition.

**Figure S12.**
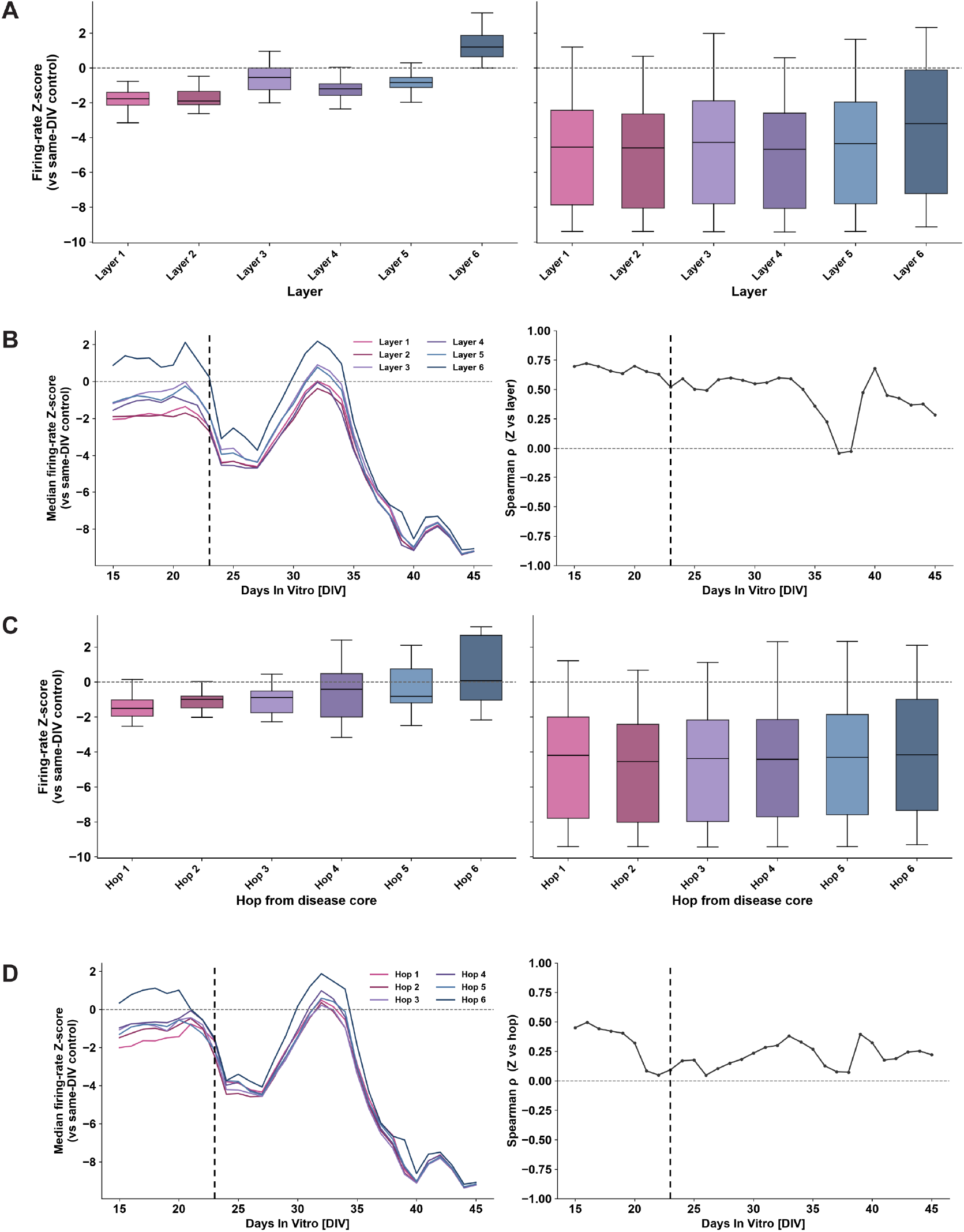
Hop distance and layer position both predict spatial gradients in disease recovery, supplementary to Figure 3. **(A)** Firing-rate Z-score (vs. same-DIV control) by layer, before (left) and after (right) A*β* application. Before A*β*, activity showed a clear gradient across layers (Spearman *ρ* = 0.66, ***, *p* = 3.4 × 10^−33^), with Layer 6 above control and Layers 1–2 most suppressed. After A*β*, this gradient largely collapsed (*ρ* = 0.09, *, *p* = 1.1 × 10^−2^), as all layers dropped to comparably low activity levels. **(B)** Median firing-rate Z-score by layer over time (left), with the corresponding layer-wise Spearman *ρ* tracked across DIV (right). The layer gradient was strong prior to A*β* addition, weakened transiently, and partially re-emerged during the recovery window (DIV 24–34: *ρ* = 0.23, ***, *p* = 2.0 × 10^−5^), with higher layers recovering toward control levels sooner than lower layers, before all groups declined together after DIV ∼35. **(C)** As in (A), but grouped by hop distance from the progerin-expressing disease core rather than by layer. A gradient was present pre-A*β* (*ρ* = 0.32, ***, *p* = 1.6 × 10^−7^) but was not statistically significant post-A*β* (*ρ* = 0.04, n.s., *p* = 2.5 × 10^−1^). **(D)** As in (B), but for hop distance. The hop-distance gradient was weaker than the layer gradient overall and did not reach significance during the recovery window (DIV 24–34: *ρ* = 0.09, n.s., *p* = 9.0 × 10^−2^), though nodes farther from the disease core showed a modest trend toward earlier recovery. Dashed vertical line indicates timing of A*β* addition.

**Figure S13.**
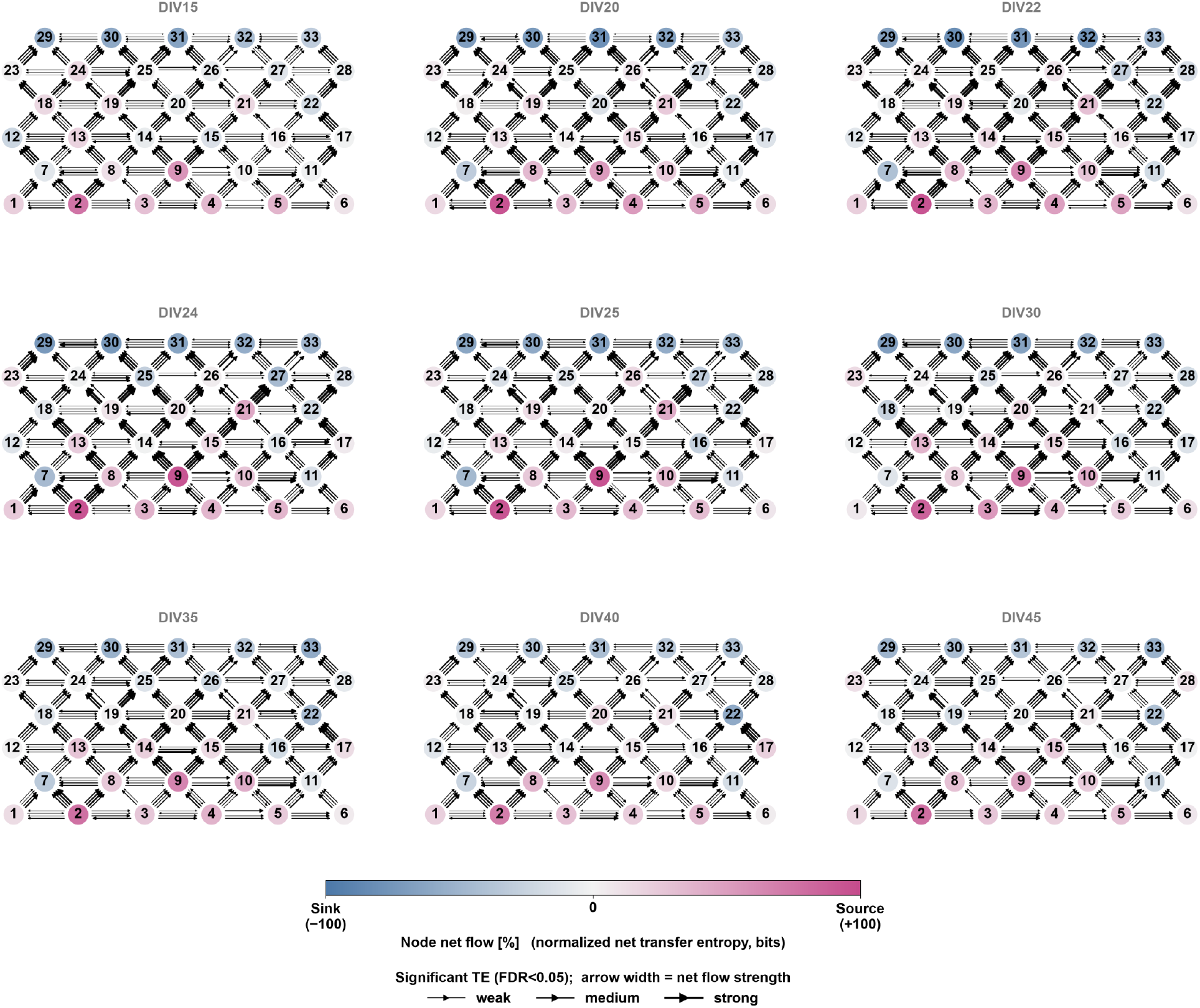
Longitudinal transfer entropy network maps for the control network, supplementary to Figure 4. Node-level functional connectivity maps at nine timepoints (DIV 15–45), showing significant directed connections (FDR *<* 0.05) as arrows, with arrow width proportional to net transfer entropy strength. Node colour indicates net information flow, with pink denoting net source behaviour and blue denoting net sink behaviour (normalised net transfer entropy, range −100 to +100%). The control network maintained relatively stable feedforward directionality and source-sink organisation throughout the recording period, with no marked reorganisation of functional connectivity over time.

**Figure S14.**
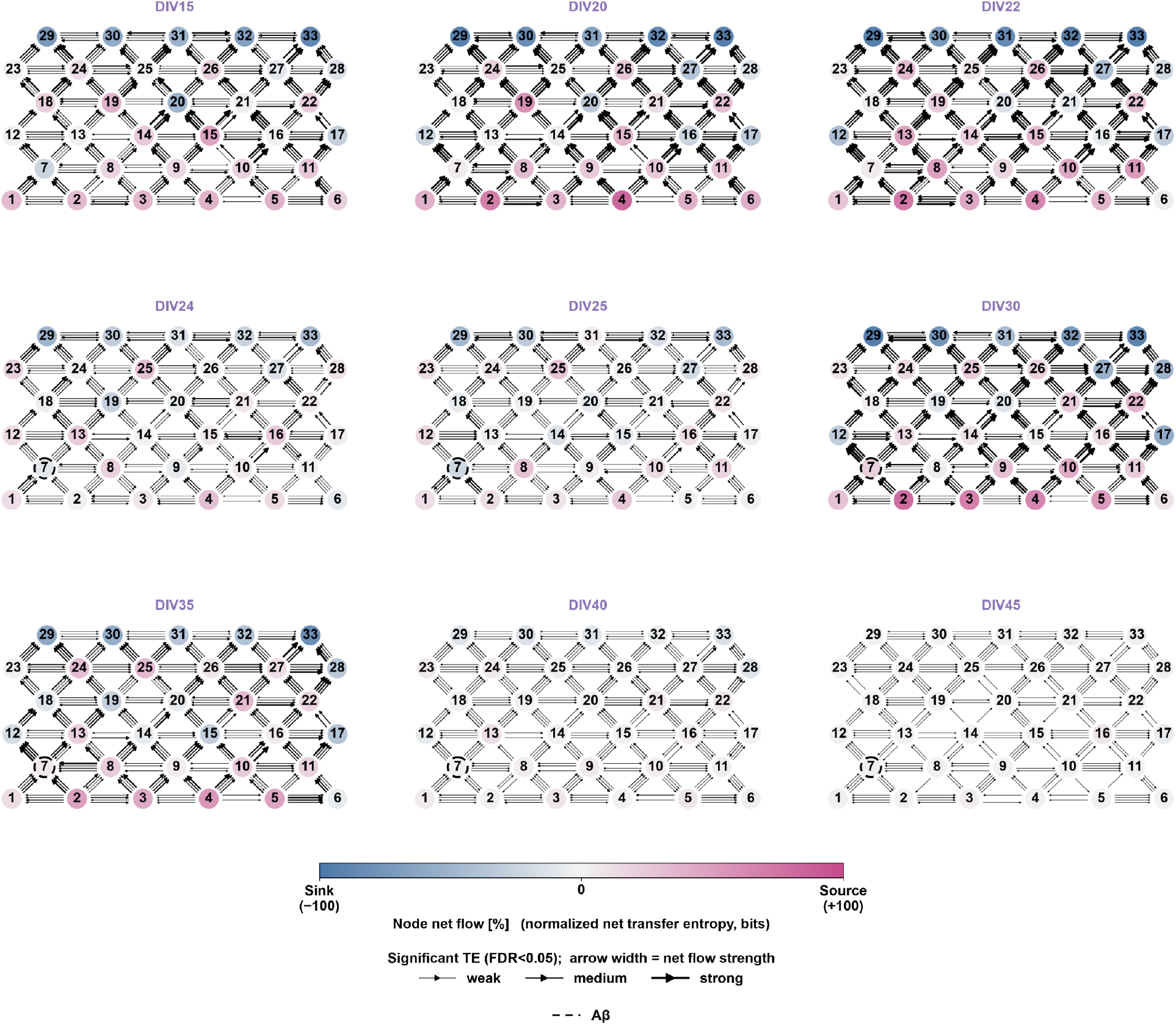
Longitudinal transfer entropy network maps for the progerin/A*β* network, supplementary to Figure 4. Node-level functional connectivity maps at nine timepoints (DIV 15–45), showing significant directed connections (FDR *<* 0.05) as arrows, with arrow width proportional to net transfer entropy strength. Node colour indicates net information flow, with pink denoting net source behaviour and blue denoting net sink behaviour (normalised net transfer entropy, range −100 to +100%). Prior to A*β* addition, the network displays active source-sink dynamics across multiple nodes. Following A*β* addition at DIV 23 (indicated by dashed node outlines), a progressive loss of significant connections and reduction in net information flow was observed, with the network approaching a near-silent state by DIV 40–45. Dashed node outlines indicate timepoints following A*β* addition.

**Figure S15.**
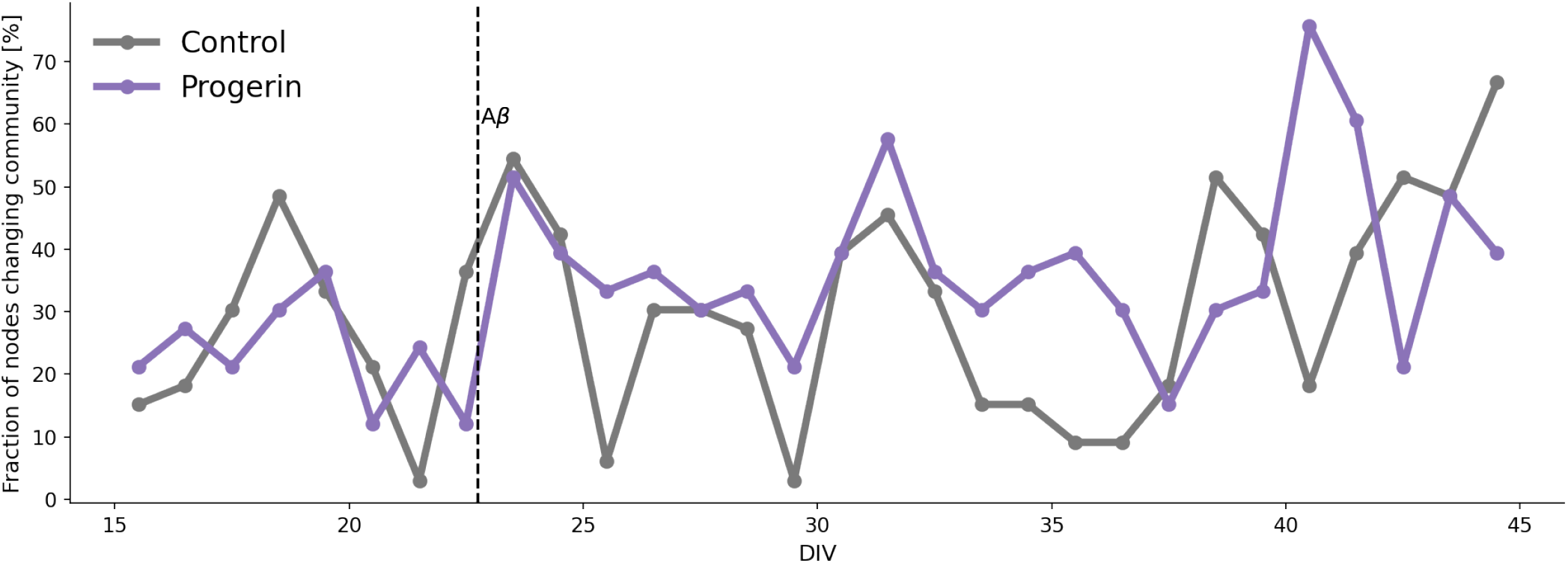
Node-level community membership instability in control and progerin/A*β* networks, supplementary to Figure 4. Fraction of nodes changing community assignment between consecutive recording days, for control (grey) and progerin/A*β* (purple) networks, over DIV. Community membership was comparably stable between conditions prior to A*β* addition. Following A*β* addition, the progerin/A*β* network showed increased and more variable node reassignment between communities relative to control.

**Figure S16.**
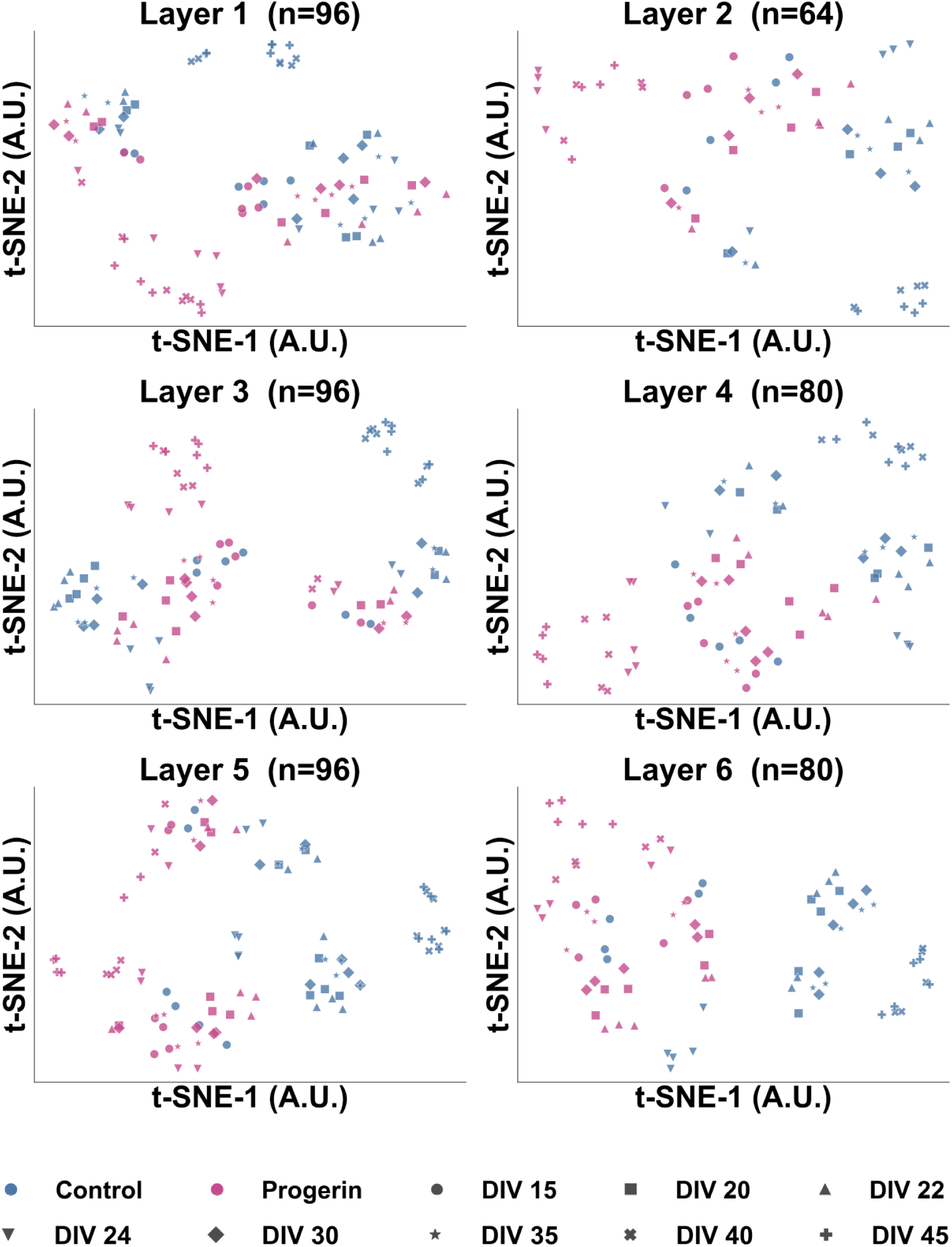
t-SNE projections of multiparametric electrophysiological feature vectors stratified by layer, supplementary to Figure 5. Each panel shows the t-SNE embedding for nodes within a single layer (layers 1–6), with points coloured by condition (blue: control, pink: progerin/A*β*) and shaped by timepoint (DIV 15–45). Within each layer, progressive separation between control and progerin/A*β* nodes was visible over time, consistent with the longitudinal divergence observed in the full network t-SNE in Figure 5A. Lower layers (1–2) showed earlier and more complete separation between conditions, while higher layers (5–6) displayed greater overlap between conditions at later timepoints, consistent with a spatial gradient of disease-relevant electrophysiological change that decays with increasing distance from the progerin node in layer 2. The number of observations per layer (*n*) reflects the number of nodes in that layer across all recorded timepoints.

**Figure S17.**
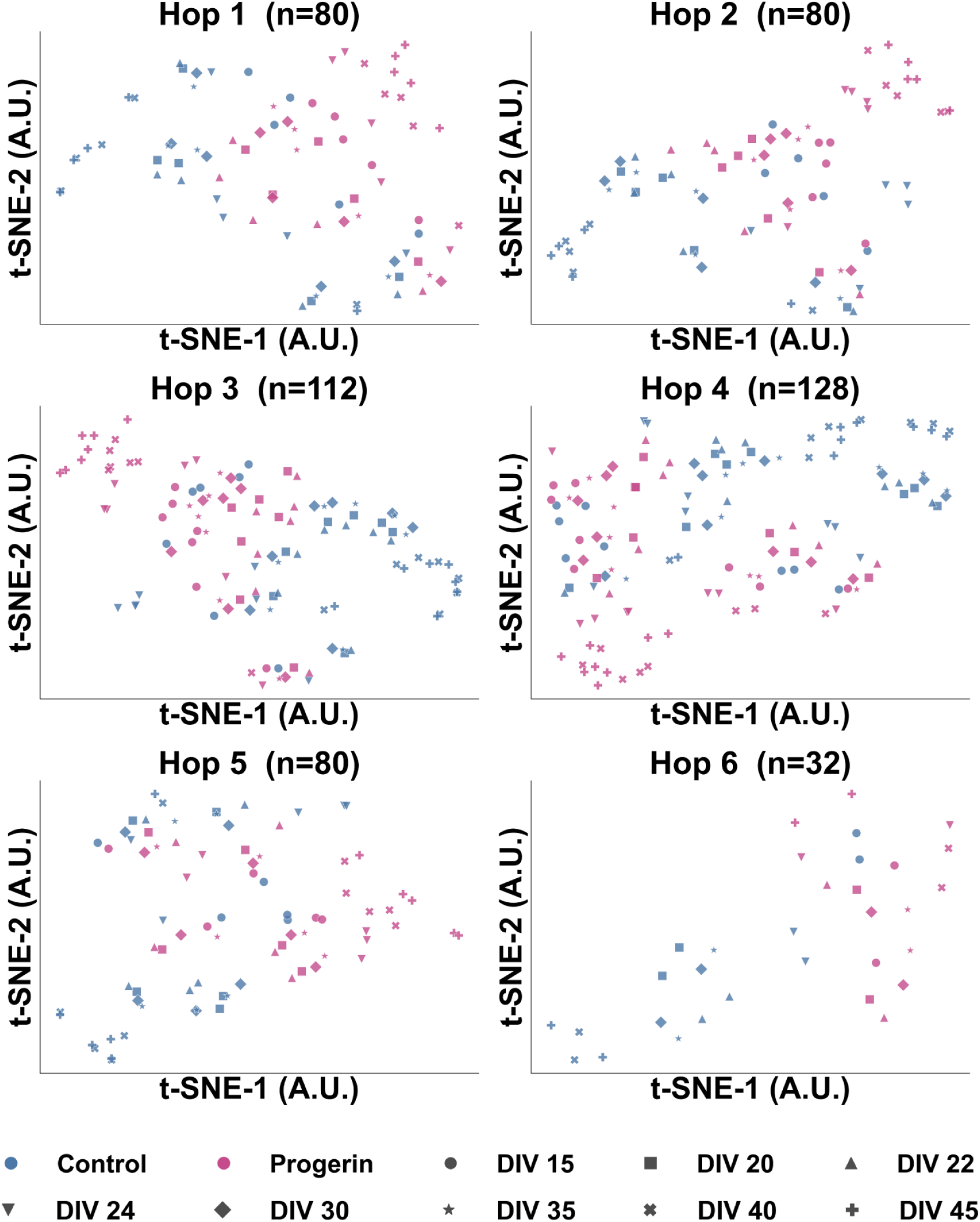
t-SNE projections of multiparametric electrophysiological feature vectors stratified by hop distance from the progerin node, supplementary to Figure 5. Each panel shows the t-SNE embedding for nodes at a given hop distance from the progerin-expressing node (hop 1–6), with points coloured by condition (blue: control, pink: progerin/A*β*) and shaped by timepoint (DIV 15–45). Nodes at hop distance 1, directly connected to the progerin node, showed early and pronounced separation between conditions. Separation between conditions became progressively less distinct with increasing hop distance, consistent with a spatial gradient of disease-relevant electrophysiological change that decays with topological distance from the disease core. At hop distance 6, the two conditions remained largely overlapping across all timepoints, suggesting that the most distal nodes retain electrophysiological signatures closer to the control condition throughout the recording period. The number of observations per hop distance (*n*) reflects the number of nodes at that distance across all recorded timepoints.

**Figure S18.**
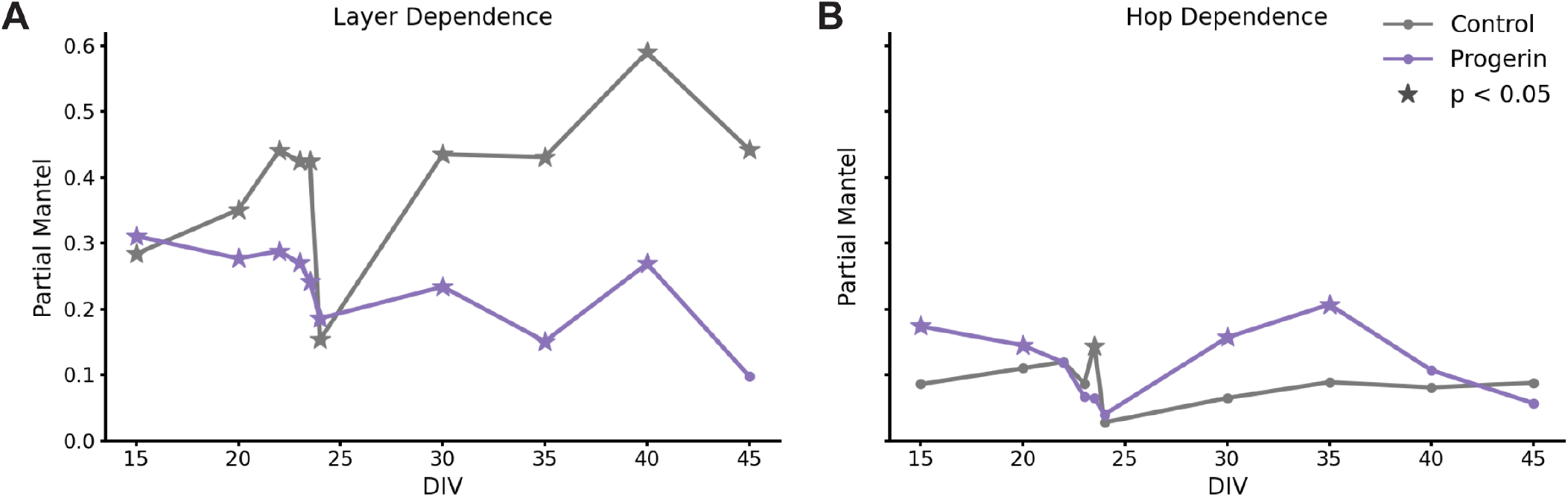
Multiparametric feature-space separation is structured by layer and hop distance from the disease core, supplementary to Figure 5. Partial Mantel correlations between pairwise multiparametric feature-vector distance (t-SNE feature space, Fig. 5A) and structural distance from the progerin-expressing node, for control (grey) and progerin/A*β* (purple) networks, over DIV. **(A)** Partial Mantel correlation between feature-vector distance and layer distance, controlling for hop distance. Control networks displayed a stronger and more variable layer-dependent structuring of feature space over time, while the progerin/A*β* network remained comparatively low and stable throughout the recording period. **(B)** Partial Mantel correlation between feature-vector distance and hop distance, controlling for layer distance. Correlations remained low in both conditions throughout the recording period, with the progerin/A*β* network showing a modest, transient increase relative to control around DIV 30–35. Stars indicate timepoints at which the partial Mantel correlation was statistically significant (*p <* 0.05).

**Figure S19.**
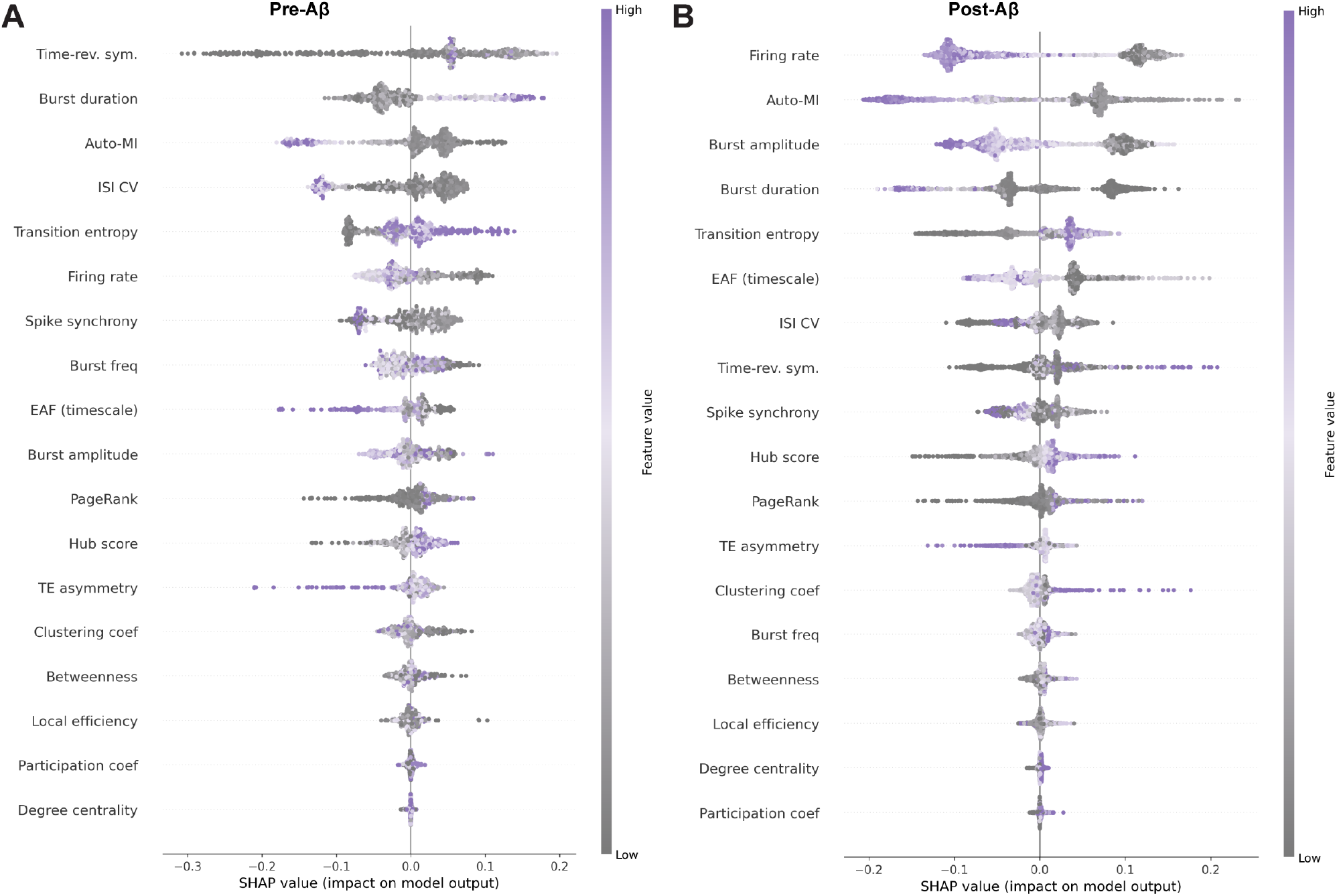
SHAP beeswarm plots showing per-feature contributions to the Random Forest classifier distinguishing control from progerin/A*β* nodes, supplementary to Figure 5. Each dot represents one node-timepoint observation. The horizontal position indicates the SHAP value, reflecting the magnitude and direction of each feature’s contribution to the model output, where positive values push toward the progerin/A*β* classification and negative values push toward the control classification. Point colour indicates the feature value, from low (grey) to high (purple). Features are ranked by mean absolute SHAP value. **(A)** Pre-A*β* period (DIV *<* 23). Time-reversal symmetry was the most prominent feature, followed by burst duration and Auto-MI. **(B)** Post-A*β* period (DIV ≥ 23). Firing rate and Auto-MI were the dominant features, followed by burst amplitude and burst duration.

**Figure S20.**
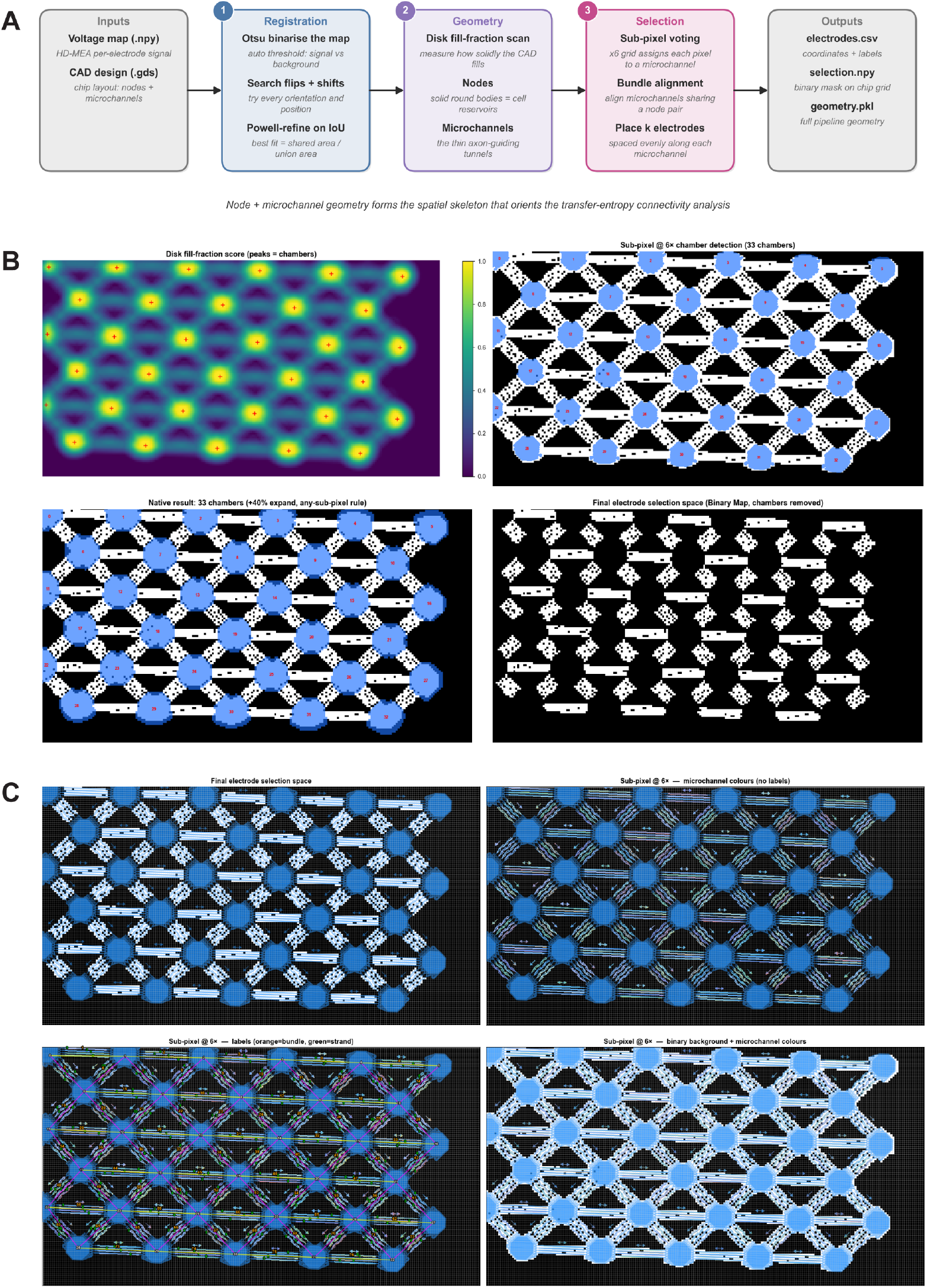
Automated electrode selection pipeline for HD-MEA recordings, supplementary to Figure 6. **(A)** Schematic overview of the three-stage pipeline. In the registration stage, the HD-MEA voltage map (.npy) and CAD design (.gds) are co-registered using Otsu binarisation, flip and shift search, and Powell-refined intersection over union (IoU) optimisation to identify the precise position and orientation of the microstructure on the chip. In the geometry stage, a disk fill-fraction scan identifies nodes (spheroid chambers) and microchannels from the CAD layout. In the selection stage, sub-pixel voting assigns electrodes to individual microchannels, bundle alignment groups electrodes sharing a node pair, and *k* electrodes are placed at evenly spaced positions along each microchannel. The pipeline outputs electrode coordinates, a binary selection mask, and the full pipeline geometry. **(B)** Intermediate outputs of the node identification stage. The disk fill-fraction score map (top left) shows peaks corresponding to spheroid chambers. Sub-pixel chamber detection at 6× resolution identifies all 33 chambers (top right). Chamber boundaries are expanded by 40% to define the node selection space (bottom left), after which chambers are removed from the final electrode selection space to restrict electrode placement to microchannel regions only (bottom right). **(C)** Intermediate outputs of the microchannel identification and electrode assignment stage. The final electrode selection space overlaid on the HD-MEA impedance map (top left). Microchannels identified at sub-pixel resolution and colour-coded by identity (top right). Bundle and strand labels assigned to each electrode (bottom left, orange: bundle, green: strand). Final electrode assignments overlaid on the binary microchannel map (bottom right).

**Figure S21.**
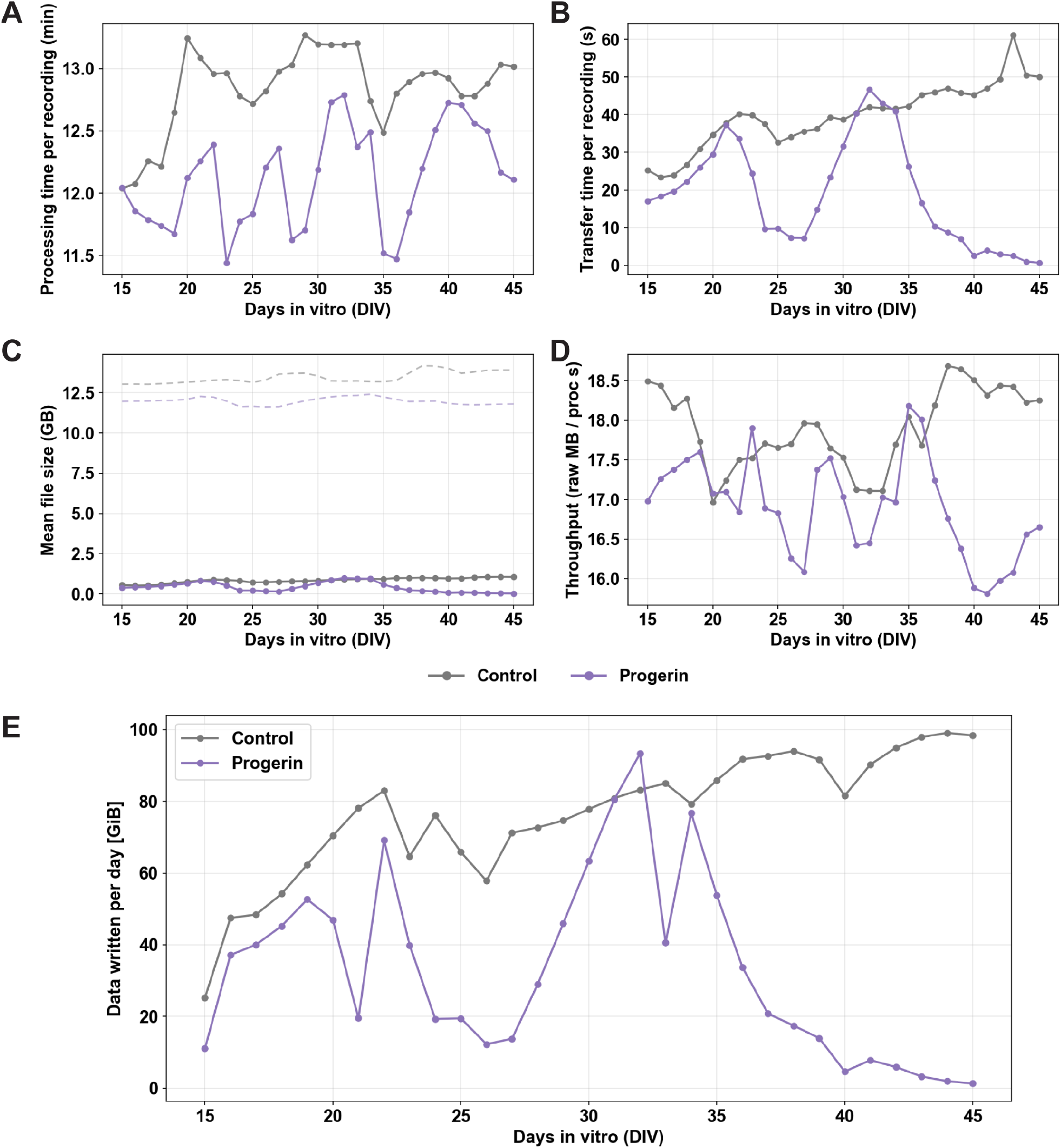
Continuous HD-MEA recording pipeline performance across the full recording period, supplementary to Figure 6. **(A)** Processing time per recording session for control and progerin/A*β* networks across DIV 15–45, remaining stable at approximately 12–13 min per session throughout the recording period for both conditions. The amount of data written was proportional to the number of detected spikes in **Fig. 3. (B)** Data transfer time per recording session. Transfer time in the progerin/A*β* network declines progressively from DIV 35 onward, approaching zero by DIV 40–45, consistent with the reduction in network activity observed in this condition **(Fig. 3). (C)** Mean file size per recording session for raw (dashed) and processed (solid) data. Raw files were approximately 12–13 GB per block, while processed files were less than 1 GB, demonstrating an approximately 20-fold reduction in data volume following spike detection and compression. **(D)** Processing throughput expressed as raw data processed per second (raw MB/proc s), remaining broadly stable across the recording period for both conditions. **(E)** Total data written per day for control and progerin/A*β* networks. The control network maintains consistently high data output throughout the recording period, while the progerin/A*β* network shows a progressive decline from DIV 35 onward, independently corroborating the reduction in network activity following combined nodal ageing and A*β* perturbation observed in Figure 3.

**Figure S22.**
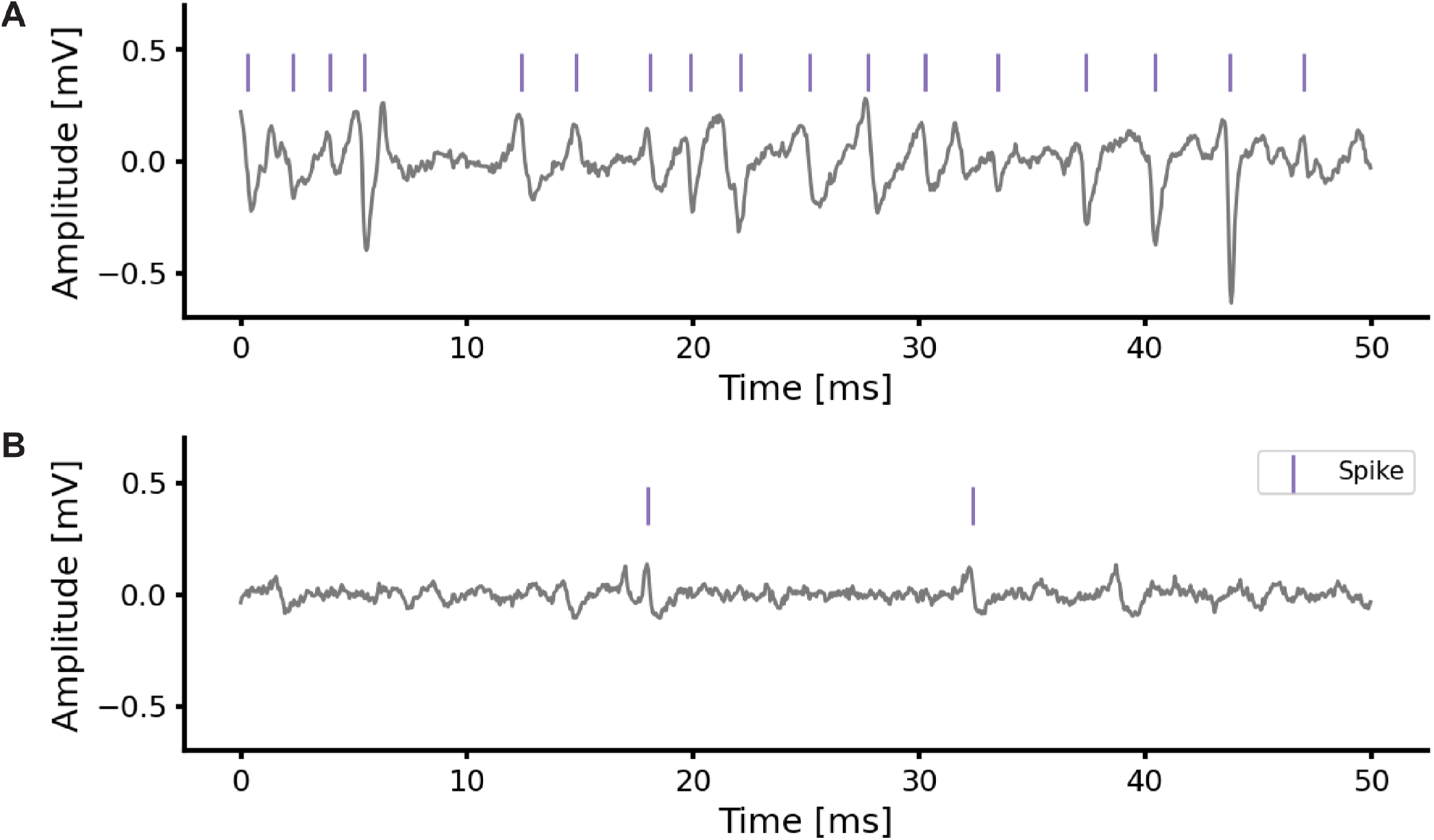
Representative raw electrode traces with detected spikes at active and inactive timepoints, supplementary to Figure 6. **(A)** Raw signal from a representative electrode during an active recording period, with detected spikes indicated by purple markers. The trace shows clear action potential waveforms with high signal-to-noise ratio and frequent spiking activity. **(B)** Raw signal from the same electrode during an inactive recording period. The trace shows low-amplitude noise with reduced spiking activity. Low amplitude action potentials are still detected using SNEO.

**Figure S23.**
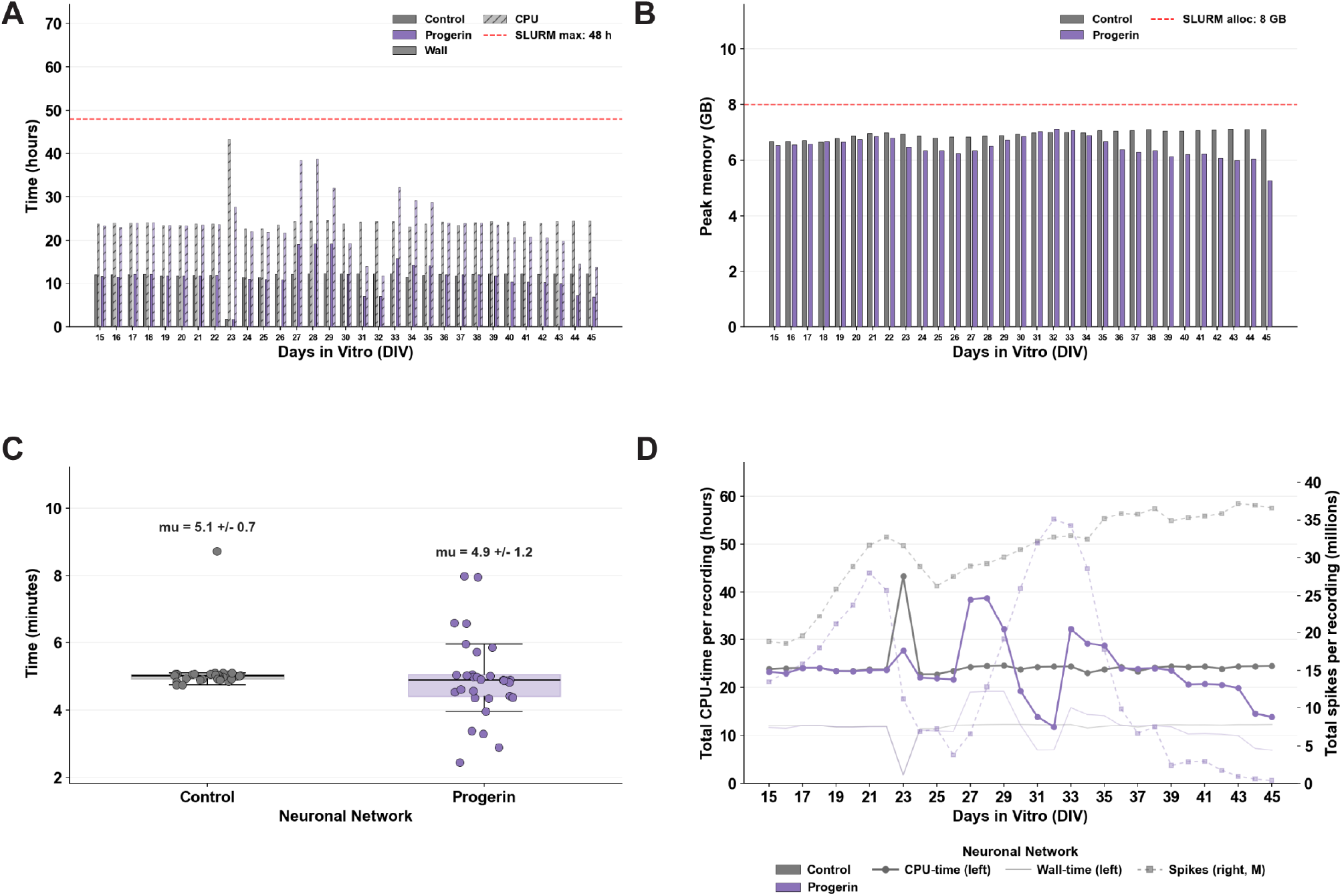
Transfer entropy computation performance across the recording period, supplementary to Figure 6. Transfer entropy was computed for all microchannel pairs across both networks using a 128-task parallel array on a SLURM cluster (2 CPUs, 8 GB memory allocated per task; bin size 0.25 ms, maximum lag 2.00 ms, maximum history target *k* = 1.00 ms, 100 permutations, FDR *α* = 0.05). **(A)** Wall time and CPU time per recording block for control and progerin/A*β* networks across DIV 15–45. CPU and Wall time remained within the 48-hour SLURM allocation limit. **(B)** Peak memory usage per recording block, remaining consistently below the 8 GB SLURM allocation for both conditions throughout the recording period. **(C)** Transfer entropy processing time per microchannel (both directions) for control (*µ* = 4.4 ± 1.6 min) and progerin/A*β* (*µ* = 3.7 ± 0.3 min) networks, demonstrating consistent and feasible per-channel computation times. **(D)** Total CPU time per recording session (left axis) and total spike count per recording session (right axis, millions) for control and progerin/A*β* networks across DIV 15–45. CPU time in the control network tracks closely with spike count, reflecting the activity-dependent computational cost of transfer entropy estimation. The progressive decline in both time and spike count in the progerin/A*β* network from DIV 35 onward independently corroborates the reduction in network activity observed in Figure 3.

### Supplementary Statistical Tables

#### Figure 1 – Platform Validation

##### Inhibitory Synapses

**Table 1:** Descriptive statistics of inhibitory synapse percentage across cell compositions. N indicates the number of fields of view analysed per condition, each corresponding to the neurite halo of an individual spheroid.

| Condition | Mean | SD | SE | N |
| --- | --- | --- | --- | --- |
| 50/50/0 | 37.879 | 25.483 | 8.494 | 9 |
| 50/50/20 | 14.019 | 6.761 | 2.390 | 8 |
| 50/50/50 | 29.824 | 21.413 | 7.138 | 9 |
| 80/20/0 | 5.267 | 2.116 | 0.705 | 9 |
| 80/20/20 | 12.907 | 7.069 | 2.356 | 9 |
| 80/20/50 | 11.203 | 9.709 | 3.236 | 9 |

**Table 2:** Statistical comparisons of inhibitory synapse percentage across cell compositions. Pairwise comparisons performed using one-way ANOVA with Tukey’s correction for multiple comparisons.

| Comparison | Adjusted p-value | Significance |
| --- | --- | --- |
| 50/50/0 vs. 50/50/20 | 0.02116 | * |
| 50/50/0 vs. 50/50/50 | 0.85756 | ns |
| 50/50/0 vs. 80/20/0 | 0.00036 | *** |
| 50/50/0 vs. 80/20/20 | 0.01033 | * |
| 50/50/0 vs. 80/20/50 | 0.00511 | ** |
| 50/50/20 vs. 50/50/50 | 0.26195 | ns |
| 50/50/20 vs. 80/20/0 | 0.82863 | ns |
| 50/50/20 vs. 80/20/20 | 0.99999 | ns |
| 50/50/20 vs. 80/20/50 | 0.99875 | ns |
| 50/50/50 vs. 80/20/0 | 0.01220 | * |
| 50/50/50 vs. 80/20/20 | 0.17180 | ns |
| 50/50/50 vs. 80/20/50 | 0.10328 | ns |
| 80/20/0 vs. 80/20/20 | 0.88256 | ns |
| 80/20/0 vs. 80/20/50 | 0.95675 | ns |
| 80/20/20 vs. 80/20/50 | 0.99988 | ns |

Significance levels: \*\*\* $p < 0.001$ , \*\* $p < 0.01$ , \* $p < 0.05$ , ns = not significant.

#### Figure 2 – Disease Model Validation

##### Progerin Expression

###### GFP-Progerin Expression

**Table 3:**
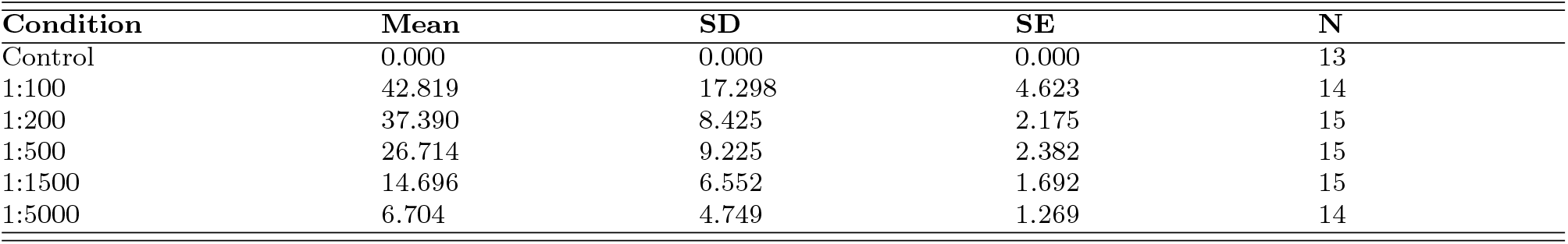
Descriptive statistics of GFP-progerin expression across viral dilutions. N indicates the number of spheroids analysed per condition.

**Table 4:** Statistical comparisons of GFP-progerin expression across viral dilutions. Pairwise comparisons performed using one-way ANOVA with Tukey’s correction for multiple comparisons.

| Comparison | Adjusted p-value | Significance |
| --- | --- | --- |
| Control vs. 1:100 | < 0.00001 | *** |
| Control vs. 1:200 | < 0.00001 | *** |
| Control vs. 1:500 | < 0.00001 | *** |
| Control vs. 1:1500 | 0.00111 | ** |
| Control vs. 1:5000 | 0.43114 | ns |
| 1:100 vs. 1:200 | 0.62358 | ns |
| 1:100 vs. 1:500 | 0.00019 | *** |
| 1:100 vs. 1:1500 | < 0.00001 | *** |
| 1:100 vs. 1:5000 | < 0.00001 | *** |
| 1:200 vs. 1:500 | 0.02822 | * |
| 1:200 vs. 1:1500 | < 0.00001 | *** |
| 1:200 vs. 1:5000 | < 0.00001 | *** |
| 1:500 vs. 1:1500 | 0.00886 | ** |
| 1:500 vs. 1:5000 | 0.00000 | *** |
| 1:1500 vs. 1:5000 | 0.20453 | ns |

Significance levels: \*\*\* $p < 0.001$ , \*\* $p < 0.01$ , \* $p < 0.05$ , ns = not significant.

##### Cells per Spheroid

**Table 5:** Descriptive statistics of cells per spheroid across viral dilutions. N indicates the number of spheroids analysed per condition.

| Condition | Mean | SD | SE | N |
| --- | --- | --- | --- | --- |
| Control | 55.692 | 16.403 | 4.549 | 13 |
| 1:100 | 25.400 | 14.436 | 3.727 | 15 |
| 1:200 | 47.067 | 10.423 | 2.691 | 15 |
| 1:500 | 50.267 | 14.048 | 3.627 | 15 |
| 1:1500 | 48.733 | 8.623 | 2.226 | 15 |
| 1:5000 | 50.786 | 12.897 | 3.447 | 14 |

**Table 6:**
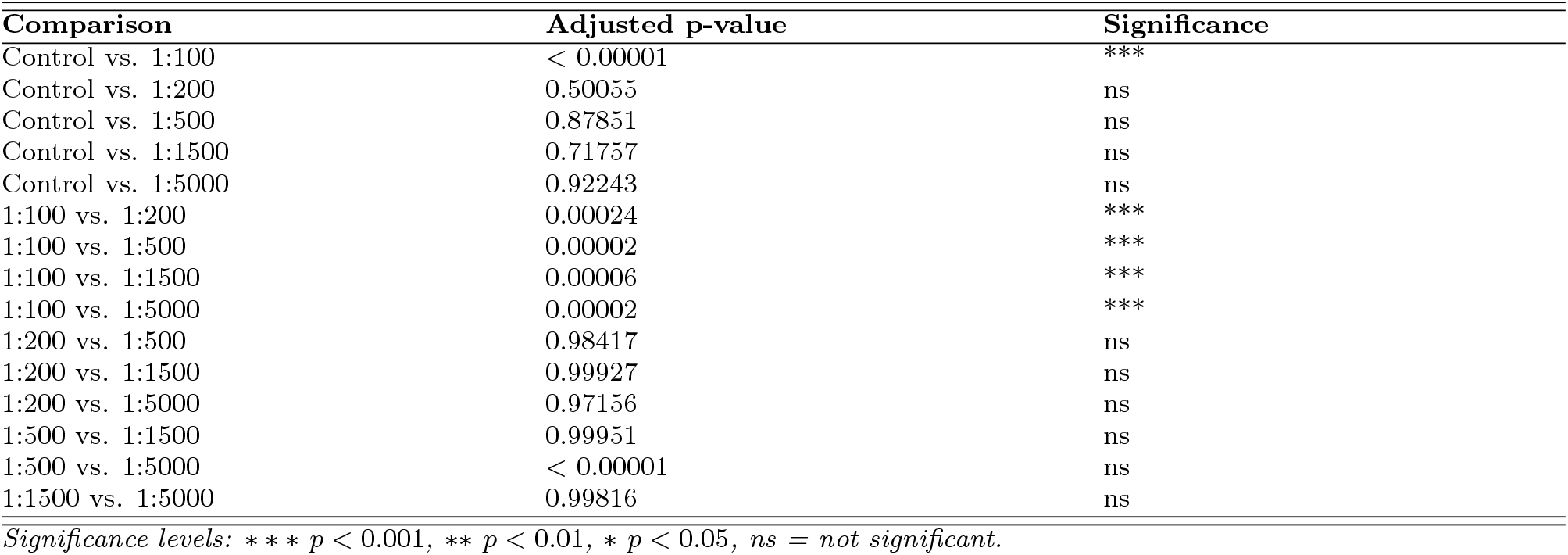
Statistical comparisons of cells per spheroid across viral dilutions. Pairwise comparisons performed using one-way ANOVA with Tukey’s correction for multiple comparisons.

##### Neurite Density

**Table 7:** Descriptive statistics of total neurite length across viral dilutions. N indicates the number of fields of view analysed per condition, each corresponding to the neurite halo of an individual spheroid.

| Condition | Mean | SD | SE | N |
| --- | --- | --- | --- | --- |
| Control | 7710.883 | 880.392 | 293.464 | 9 |
| 1:100 | 3479.346 | 1771.343 | 590.448 | 9 |
| 1:200 | 3198.999 | 1314.880 | 438.293 | 9 |
| 1:500 | 4390.835 | 1465.018 | 488.339 | 9 |
| 1:1500 | 6376.272 | 1893.449 | 631.150 | 9 |
| 1:5000 | 7065.601 | 1122.155 | 374.052 | 9 |

**Table 8:** Statistical comparisons of total neurite length across viral dilutions. Pairwise comparisons performed against the control condition only using one-way ANOVA with Tukey’s correction for multiple comparisons.

| Comparison | Adjusted p-value | Significance |
| --- | --- | --- |
| Control vs. 1:100 | < 0.00001 | *** |
| Control vs. 1:200 | < 0.00001 | *** |
| Control vs. 1:500 | 0.00007 | *** |
| Control vs. 1:1500 | 0.25386 | ns |
| Control vs. 1:5000 | 0.88416 | ns |

Significance levels: \*\*\* $p < 0.001$ , \*\* $p < 0.01$ , \* $p < 0.05$ , ns = not significant.

##### Oxidative Stress

**Table 9:** Descriptive statistics of normalised CellROX integrated density across viral dilutions. N indicates the number of spheroids analysed per condition.

| Condition | Mean | SD | SE | N |
| --- | --- | --- | --- | --- |
| Control | 42160.23 | 11919.26 | 3973.088 | 9 |
| 1:100 | 32083.93 | 17080.95 | 5693.650 | 9 |
| 1:200 | 35659.12 | 18638.41 | 6212.802 | 9 |
| 1:500 | 43730.08 | 13517.02 | 4505.675 | 9 |
| 1:1500 | 44471.76 | 12656.26 | 4218.754 | 9 |
| 1:5000 | 40778.62 | 13958.40 | 4935.039 | 8 |

**Table 10:** Statistical comparisons of normalised CellROX integrated density across viral dilutions. Pairwise comparisons performed against the control condition only using one-way ANOVA with Tukey’s correction for multiple comparisons.

| Comparison | Adjusted p-value | Significance |
| --- | --- | --- |
| Control vs. 1:100 | 0.57303 | ns |
| Control vs. 1:200 | 0.89061 | ns |
| Control vs. 1:500 | 0.99983 | ns |
| Control vs. 1:1500 | 0.99887 | ns |
| Control vs. 1:5000 | 0.99992 | ns |

Significance levels: \*\*\* $p < 0.001$ , \*\* $p < 0.01$ , \* $p < 0.05$ , ns = not significant.

##### Amyloid-*β* Concentration Effects

**Table 11:** Descriptive statistics of intraneuronal A*β* aggregate size across concentrations. N indicates the number of fields of view analysed per condition, each corresponding to the neurite halo of an individual spheroid.

| Condition | Mean | SD | SE | N |
| --- | --- | --- | --- | --- |
| Control | 1.444 | 1.120 | 0.354 | 10 |
| 0.1 μM | 1.430 | 0.071 | 0.022 | 10 |
| 0.2 μM | 1.788 | 0.109 | 0.034 | 10 |
| 0.5 μM | 2.762 | 0.264 | 0.083 | 10 |
| 1 μM | 2.633 | 0.218 | 0.069 | 10 |
| 1.5 μM | 2.991 | 0.593 | 0.188 | 10 |

**Table 12:**
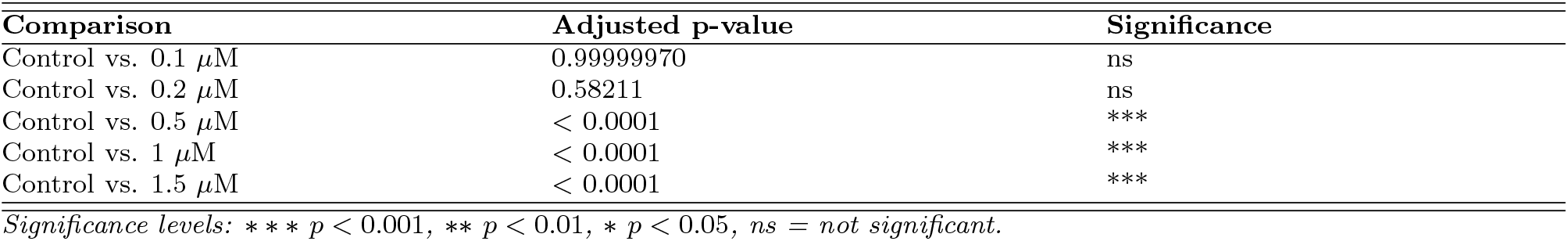
Statistical comparisons of intraneuronal A*β* aggregate size across concentrations. Pairwise comparisons performed against the control condition only using one-way ANOVA with Tukey’s correction for multiple comparisons.

**Table 13:** Descriptive statistics of intraneuronal A*β* aggregate count across concentrations. N indicates the number of fields of view analysed per condition, each corresponding to the neurite halo of an individual spheroid.

| Condition | Mean | SD | SE | N |
| --- | --- | --- | --- | --- |
| Control | 1.900 | 2.079 | 0.657 | 10 |
| 0.1 μM | 172.900 | 27.020 | 8.544 | 10 |
| 0.2 μM | 330.800 | 56.645 | 17.913 | 10 |
| 0.5 μM | 421.800 | 102.737 | 32.488 | 10 |
| 1 μM | 545.700 | 73.164 | 23.136 | 10 |
| 1.5 μM | 498.600 | 128.281 | 40.566 | 10 |

**Table 14:** Statistical comparisons of intraneuronal A*β* aggregate count across concentrations. Pairwise comparisons performed against the control condition only using one-way ANOVA with Tukey’s correction for multiple comparisons.

| Comparison | Adjusted p-value | Significance |
| --- | --- | --- |
| Control vs. 0.1 μM | < 0.0001 | *** |
| Control vs. 0.2 μM | < 0.0001 | *** |
| Control vs. 0.5 μM | < 0.0001 | *** |
| Control vs. 1 μM | < 0.0001 | *** |
| Control vs. 1.5 μM | < 0.0001 | *** |

**Table 15:** Descriptive statistics of neurite mesh area fraction across concentrations. N indicates the number of fields of view analysed per condition, each corresponding to the neurite halo of an individual spheroid.

| Condition | Mean | SD | SE | N |
| --- | --- | --- | --- | --- |
| Control | 11144.076 | 1916.565 | 606.071 | 10 |
| 0.1 μM | 12806.265 | 1965.861 | 621.660 | 10 |
| 0.2 μM | 10041.654 | 3217.302 | 1017.400 | 10 |
| 0.5 μM | 9472.841 | 1909.066 | 603.700 | 10 |
| 1 μM | 8267.976 | 1376.015 | 435.134 | 10 |
| 1.5 μM | 4841.546 | 1971.099 | 623.316 | 10 |

**Table 16:**
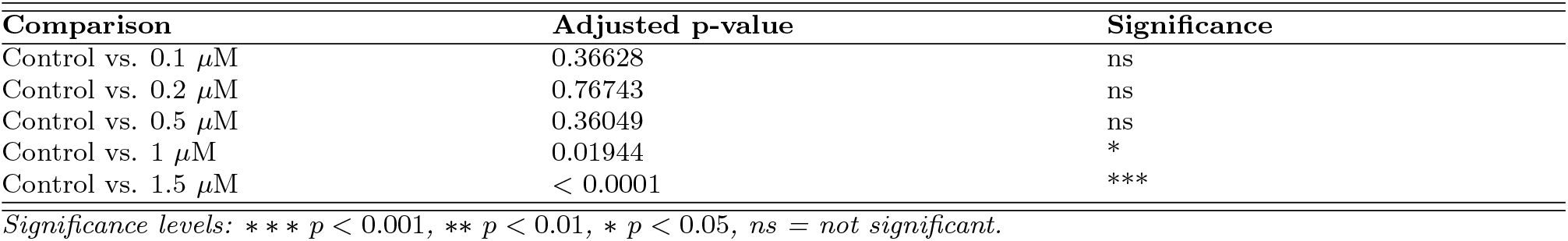
Statistical comparisons of neurite mesh area fraction across concentrations. Pairwise comparisons performed against the control condition only using one-way ANOVA with Tukey’s correction for multiple comparisons.

| Comparison | Adjusted p-value | Significance |
| --- | --- | --- |
| Control vs. 0.1 μM | 0.36628 | ns |
| Control vs. 0.2 μM | 0.76743 | ns |
| Control vs. 0.5 μM | 0.36049 | ns |
| Control vs. 1 μM | 0.01944 | * |
| Control vs. 1.5 μM | < 0.0001 | *** |

Significance levels: \*\*\* $p < 0.001$ , \*\* $p < 0.01$ , \* $p < 0.05$ , ns = not significant.

#### Figure S7 – Supplementary Disease Model Validation

##### Supplementary Amyloid-*β* Concentration Effects

**Table 17:** Descriptive statistics of intraneuronal A*β* aggregate density across concentrations. N indicates the number of fields of view analysed per condition, each corresponding to the neurite halo of an individual spheroid.

| Condition | Mean | SD | SE | N |
| --- | --- | --- | --- | --- |
| Control | 0.011 | 0.020 | 0.006 | 10 |
| 0.1 μM | 0.550 | 0.098 | 0.031 | 10 |
| 0.2 μM | 1.318 | 0.260 | 0.082 | 10 |
| 0.5 μM | 2.567 | 0.601 | 0.190 | 10 |
| 1 μM | 3.195 | 0.522 | 0.165 | 10 |
| 1.5 μM | 3.197 | 0.502 | 0.159 | 10 |

**Table 18:** Statistical comparisons of intraneuronal A*β* aggregate density across concentrations. Pairwise comparisons performed against the control condition only using one-way ANOVA with Tukey’s correction for multiple comparisons.

| Comparison | Adjusted p-value | Significance |
| --- | --- | --- |
| Control vs. 0.1 μM | 0.01960 | * |
| Control vs. 0.2 μM | < 0.0001 | *** |
| Control vs. 0.5 μM | < 0.0001 | *** |
| Control vs. 1 μM | < 0.0001 | *** |
| Control vs. 1.5 μM | < 0.0001 | *** |

**Table 19:** Descriptive statistics of neurite mesh total length across concentrations. N indicates the number of fields of view analysed per condition, each corresponding to the neurite halo of an individual spheroid.

| Condition | Mean | SD | SE | N |
| --- | --- | --- | --- | --- |
| Control | 10357.209 | 1904.716 | 602.324 | 10 |
| 0.1 $\mu\text{M}$ | 12069.865 | 1980.504 | 626.290 | 10 |
| 0.2 $\mu\text{M}$ | 9274.087 | 3037.282 | 960.473 | 10 |
| 0.5 $\mu\text{M}$ | 8779.909 | 1851.416 | 585.469 | 10 |
| 1 $\mu\text{M}$ | 7612.637 | 1227.756 | 388.251 | 10 |
| 1.5 $\mu\text{M}$ | 4250.100 | 1871.327 | 591.766 | 10 |

**Table 20:**
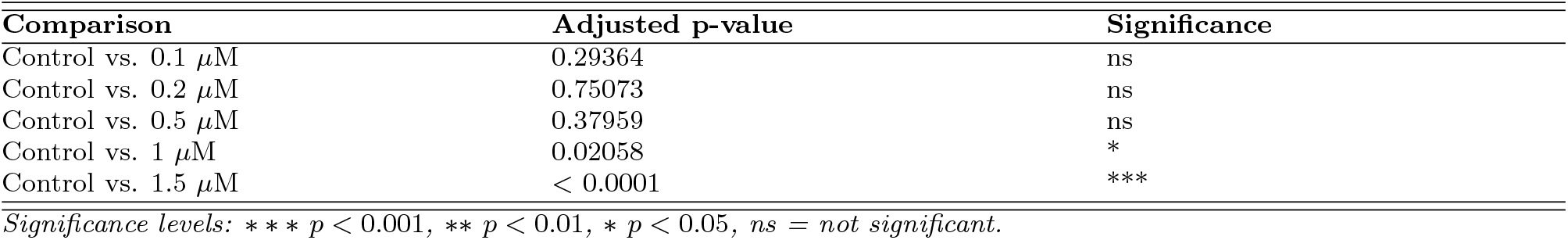
Statistical comparisons of neurite mesh total length across concentrations. Pairwise comparisons performed against the control condition only using one-way ANOVA with Tukey’s correction for multiple comparisons.

| Comparison | Adjusted p-value | Significance |
| --- | --- | --- |
| Control vs. 0.1 $\mu\text{M}$ | 0.29364 | ns |
| Control vs. 0.2 $\mu\text{M}$ | 0.75073 | ns |
| Control vs. 0.5 $\mu\text{M}$ | 0.37959 | ns |
| Control vs. 1 $\mu\text{M}$ | 0.02058 | * |
| Control vs. 1.5 $\mu\text{M}$ | < 0.0001 | *** |

Significance levels: \*\*\* $p < 0.001$ , \*\* $p < 0.01$ , \* $p < 0.05$ , ns = not significant.

**Table 21:**
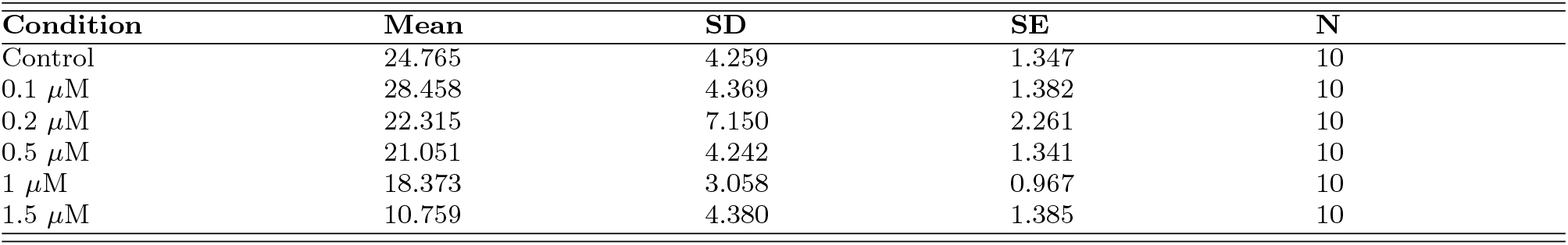
Descriptive statistics of neurite mesh density across concentrations. N indicates the number of fields of view analysed per condition, each corresponding to the neurite halo of an individual spheroid.

**Table 22:** Statistical comparisons of neurite mesh density across concentrations. Pairwise comparisons performed against the control condition only using one-way ANOVA with Tukey’s correction for multiple comparisons.

| Comparison | Adjusted p-value | Significance |
| --- | --- | --- |
| Control vs. 0.1 $\mu\text{M}$ | 0.36628 | ns |
| Control vs. 0.2 $\mu\text{M}$ | 0.76742 | ns |
| Control vs. 0.5 $\mu\text{M}$ | 0.36049 | ns |
| Control vs. 1 $\mu\text{M}$ | 0.01944 | * |
| Control vs. 1.5 $\mu\text{M}$ | < 0.0001 | *** |

Significance levels: \*\*\* $p < 0.001$ , \*\* $p < 0.01$ , \* $p < 0.05$ , ns = not significant.

**Table 23:**
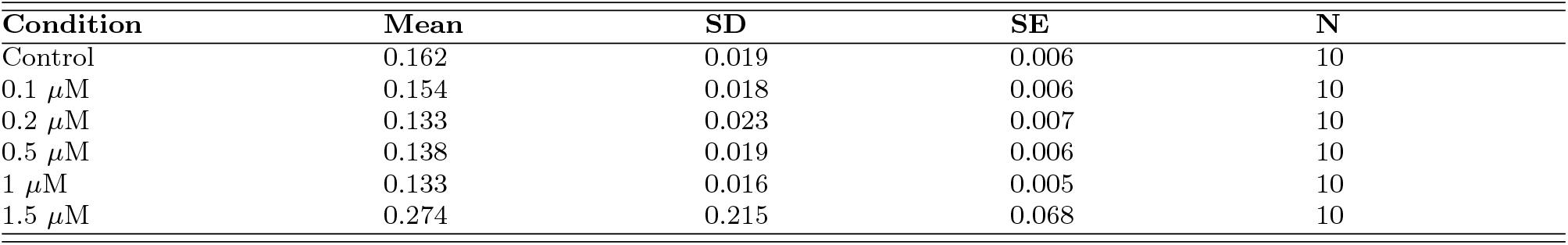
Descriptive statistics of endpoint density across concentrations. N indicates the number of fields of view analysed per condition, each corresponding to the neurite halo of an individual spheroid.

**Table 24:**
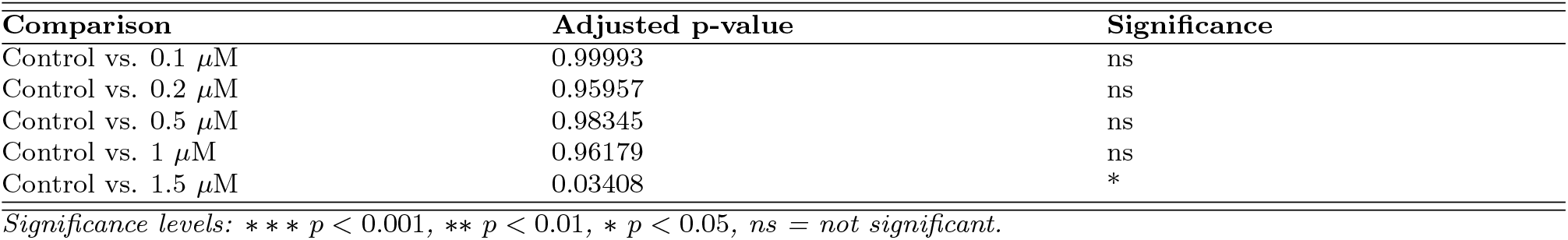
Statistical comparisons of endpoint density across concentrations. Pairwise comparisons performed against the control condition only using one-way ANOVA with Tukey’s correction for multiple comparisons.

**Table 25:** Descriptive statistics of number of junctions across concentrations. N indicates the number of fields of view analysed per condition, each corresponding to the neurite halo of an individual spheroid.

| Condition | Mean | SD | SE | N |
| --- | --- | --- | --- | --- |
| Control | 965.700 | 301.039 | 95.197 | 10 |
| 0.1 $\mu\text{M}$ | 1248.700 | 331.229 | 104.744 | 10 |
| 0.2 $\mu\text{M}$ | 748.700 | 403.957 | 127.742 | 10 |
| 0.5 $\mu\text{M}$ | 699.300 | 292.379 | 92.458 | 10 |
| 1 $\mu\text{M}$ | 553.600 | 163.734 | 51.777 | 10 |
| 1.5 $\mu\text{M}$ | 259.000 | 128.176 | 40.533 | 10 |

**Table 26:** Statistical comparisons of number of junctions across concentrations. Pairwise comparisons performed against the control condition only using one-way ANOVA with Tukey’s correction for multiple comparisons.

| Comparison | Adjusted p-value | Significance |
| --- | --- | --- |
| Control vs. 0.1 $\mu\text{M}$ | 0.14751 | ns |
| Control vs. 0.2 $\mu\text{M}$ | 0.39632 | ns |
| Control vs. 0.5 $\mu\text{M}$ | 0.19440 | ns |
| Control vs. 1 $\mu\text{M}$ | 0.01090 | * |
| Control vs. 1.5 $\mu\text{M}$ | < 0.0001 | *** |

Significance levels: \*\*\* $p < 0.001$ , \*\* $p < 0.01$ , \* $p < 0.05$ , ns = not significant.

## Notes

### Competing Interest Statement

The authors have declared no competing interest.

